# Bioinformatics Analysis of Non-Coding UTR Variants in *STAT1* Reveals Disruption of miRNA Binding, mRNA Stability, and Oncogenic Potential

**DOI:** 10.1101/2025.01.16.632379

**Authors:** Ebtihal Kamal, Mehad Ahmed

## Abstract

Single nucleotide polymorphisms (SNPs) are associated with a wide range of disorders, including diverse cancer types. In the context of cancer, alterations within non-coding regions, specifically untranslated regions (UTRs), have proven substantially important. SNPs localized in the 3 prime UTRs of the *STAT1* gene were assessed for their association with miRNAs using PolymiRTS, miRNASNP, and MicroSNIper. In addition, the 5 UTR SNPs were analyzed by SNP INFO for changes in the transcription factors binding sites. The significant SNPs were further analysed by Cscape to predict their oncogenic probability. The secondary structures of the wild type and the mutant mRNA were analysed by RNAfold.

Out of 605 SNPs analyzed, 14 UTR SNPs (six in the 3⍰ UTR region and eight in the 5⍰ UTR region) were identified with functional annotation ≤2a by RegulomeDB. The associations of 8 SNPs with miRNAs common between the 3 databases (PolymiRTS, miRNASNP, and MicroSNIper) were found.

Moreover we identified six SNPs (rs1197872838, rs3088307, rs45470392, rs1413522785, rs531009254, and rs190508584) destabilize the mRNA structure resulting in substantial change in free energy (ΔG). The SNP rs45470392 in 5⍰ UTR was predicted to alter the transcription factor binding sites. rs188557905, rs1220766131, rs1413522785, rs1168, and rs999207177 were predicted to be oncogenic SNPs in this study using bioinformatics tools.

Our findings highlight the impact of 3⍰ and 5⍰ UTR SNPs on miRNA, transcription, and translation of *STAT1*. These analyses suggest that these SNPs can have substantial functional importance in the *STAT1* gene. Future experimental validation could establish their potential role in the diagnosis and therapeutics of various diseases, including cancer.

## Introduction

The Signal Transducer and Activator of Transcription (STAT) protein family comprises transcription factors that are crucial in regulating physiological cellular processes, including proliferation, differentiation, apoptosis, and angiogenesis, in addition to structuring the epigenetic landscape of immune cells [1–3].

Signal transducer and activator of transcription 1 (STAT1) is a dormant cytoplasmic transcription factor that acts as an essential component in the interferon (IFN)-signaling pathway, activated via various triggers, including type I, II, and III interferons as well as interleukin-27 and inteinterleukin-27 (4–6). STAT1 mediates a variety of cellular functions in response to stimulation by cytokines, growth factors, and hormones [4].

The role and significance of STAT1 in cancer biology have been studied for a decade. The majority of evidence shows activating STAT1 plays a tumor suppressor role in cancer cells [3–7]. Nevertheless, results from some experiments and clinical studies suggest that STAT1 also exerts tumor promoter effects [6,8,9]. In some malignant phenotypes, STAT1 can function either as an oncoprotein or tumor suppressor in the same cell type. Thus, the function of STAT1 in cancer biology and the mechanism behind the oncogenic or tumor-suppressive function of STAT1 remain mysteries [4].

Single nucleotide polymorphisms (SNPs) are defined as gene variations that have a higher frequency than other variations. Phenotypic changes are directly linked to SNPs in the coding region [10–12]. Moreover, SNPs regulate gene expression through a variety of methods and can be identified in 5’– and 3’-untranslated regions (UTRs), exons, introns, and gene promoters [13]. MicroRNAs are small, noncoding RNAs with 18–24 nucleotides. They bind to messenger RNAs’ 3⍰ untranslated regions (3⍰UTRs), where they inhibit translation and ultimately to the decay of the mRNAs [14,15]. SNPs in the 3’ UTR miRNA binding sites have contributed to another dimension of potential genetic variation implicated in the complicated mechanism of oncogenesis (16–20).

A genome-wide association study revealed the majority of the SNPs have been identified in non-coding regions and are linked to various forms of cancer [16]. Non-coding SNP localization within the UTRs is considered to be especially deleterious. It has been established that the recognized variations in these regions effect the structure of miRNA, thereby affecting its interactions with other miRNAs and the resultant protein [17–19].

The introduction of various bioinformatics tools and software has rendered it feasible to study and analyse SNPs in the non-coding regions. These online tools, which were used to identify SNPs associated with miRNA, are PolymiRTS, miRNASNP, and MicroSNIper. Results of these methods indicated the SNPs have the capacity to alter the sequence’s microRNA targets. Various diseases, including cancer, may arise from damaged microRNAs, which also play a significant role in an array of regulatory pathways [20].

Considering the importance of UTR variations in several studies and their correlation with an array of disorders, this research aimed

Since limited studies have focused on the effects of UTR SNPs in STAT1gene, specifically their role in regulating transcription factor interactions and miRNA binding sites. We aim in this study to investigate UTR SNPs in STAT1 gene using a systematic in silico approaches; determining the functional significance of SNPs; predicting the likely influence of UTR variations on creating or disrupting miRNA binding sites. Furthermore, this study aimed to figure out how STAT1 UTR SNPs affect transcription factors binding regions and mRNA structure.

## 2. Methods

### 2.1. Retrieval of UTR SNPs and Annotation Using RegulomeDB

NCBI was accessed for STAT1 on 20 December 2024 for retrieval of 3 prime and 5 prime UTR SNPs. Functional annotation and scoring of UTR variants was performed through RegulomeDB (https://regulomedb.org/regulome-search). RegulomeDB identifies regulatory roles of non-coding SNPs through integration of high-throughput data sets from various sources, including ENCODE and GEO. The function of RegulomeDB is to assign scores to SNPs so that the functional SNPs can be identified. RegulomeDB scores of STAT1 non-coding variants were obtained by entering rs IDs of individual SNPs. The UTR variants were then analyzed via various bioinformatics tools (Figure 1).

**Figure 1.**
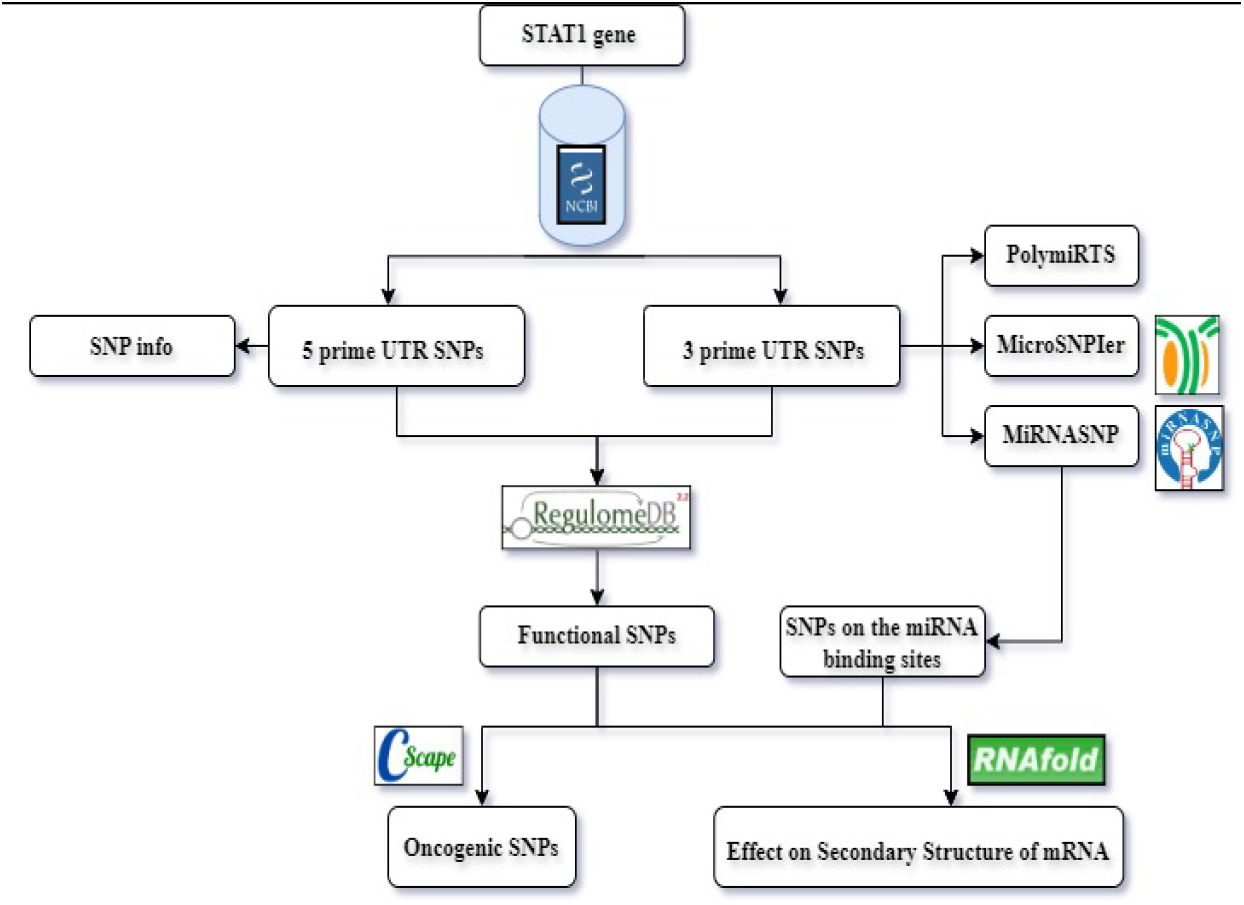
The workflow of the study

### 2.2 Assessment of Effect of 3 prime UTR SNPs on miRNA Binding Sites

The SNPs were assessed for their association with miRNA using three tools: polymorphism in miRNAs and their target sites (PolymiRTS), miRNASNP, and MicroSNIper databases. The PolymiRTS database [21] can identify the location of non-coding SNPs, whether they are in miRNA target sites or seed regions of miRNA. For analyzing the consequences of non-coding SNPs, the rs ID of each variant was submitted to the PolymiRTS database (https://compbio.uthsc.edu/miRSNP/miRSNP_detail_all.php). For prediction of loss and gain function of SNPs in pre-miRNA, mature miRNA, or located in the miRNA target sequences, miRNASNP was employed. A list of targets was produced with SNP-miRNA/duplexes, energy change, and gain/loss caused by SNP in 3 prime UTR of the gene [22]. MiRNASNP is available at (http://bioinfo.life.hust.edu.cn/miRNASNP/, accessed on 10 October 2024). MicroSNIper program was also used to determine the interaction between SNPs and miRNAs. [23] It is available at (http://vm24141.virt.gwdg.de/services/microsniper/), accessed on 10 October 2022.

### 2.3. Determination of Effect of SNPs on Secondary Structure of mRNA

The SNPs annotated as 2a and 2b after Regulome DB analysis and variants associated with miRNA were selected for further analysis. Using the RNAfold server, within Vienna RNA package [24], the minimum free energies (MFE) and mRNA secondary structures of wild-type and SNPs were obtained This package is based on minimum free energy algorithms. The results were obtained using wild-type and SNP sequences that were obtained from NCBI.

### 2.4. Analysis of 5prime UTR SNPs on Transcription Factor Binding Sites (TFBS)

The impact of 5 prime UTR variants on transcription factor binding sites was evaluated through SNPinfo (FuncPred) (https://snpinfo.niehs.nih.gov/snpinfo/snpfunc.html), which was also used to determine the potential functional effect of regulatory SNPs [25]. A list of SNP rs IDs was uploaded for batch analysis with default settings. The output was a list of SNPs with possible functional effect

### 2.5. Prediction of cancer-associated SNPs

The association of nsSNP with the development of cancer was predicted using CScape tools. CScape is a combinatorial web portal for determining the oncogenic status of single mutation in coding as well as noncoding region of genome [26] Mutations are classified as neutral or cancer driver based on P values ranging from 0–1. Value >0.5 is predicted as oncogenic. Whereas <0.5 is considered as neutral. CScape is available at http://cscape.biocompute.org.uk/

## 3. Results

### 3.1. Retrieval of UTR SNPs of STAT1 from NCBI and Scoring on RegulomeDB

A total of 999 UTR SNPs, including 838 in the 3⍰ UTR (Supplementary Table 1) and 161 in the 5⍰ UTR (Supplementary Table 2), were acquired from the NCBI database. Distribution of the SNPs was shown in Figure 2a. To check whether the SNPs are pathogenic, likely pathogenic, or acting as a risk factor for a disease, the retrieved UTR SNPs were subjected to functional evaluation through the RegulomeDB server. Out of all SNPs, 503 in the 3⍰ UTR-located and 102 in the 5⍰ UTR-located were found on RegulomeDB; all of them distributed in Figure 2b. A sum of 14 SNPs in the UTR (6 in 3⍰ UTR and 8 in 5⍰ UTR regions) was annotated as ≤2a by RegulomeDB having variable RegulomeDB scores as mentioned in figure 2c. All the SNPs in 3 and 5 UTR and their RegulomeDB rank are presented in supplementary table S3, S4, respectively.

**Table S1.**
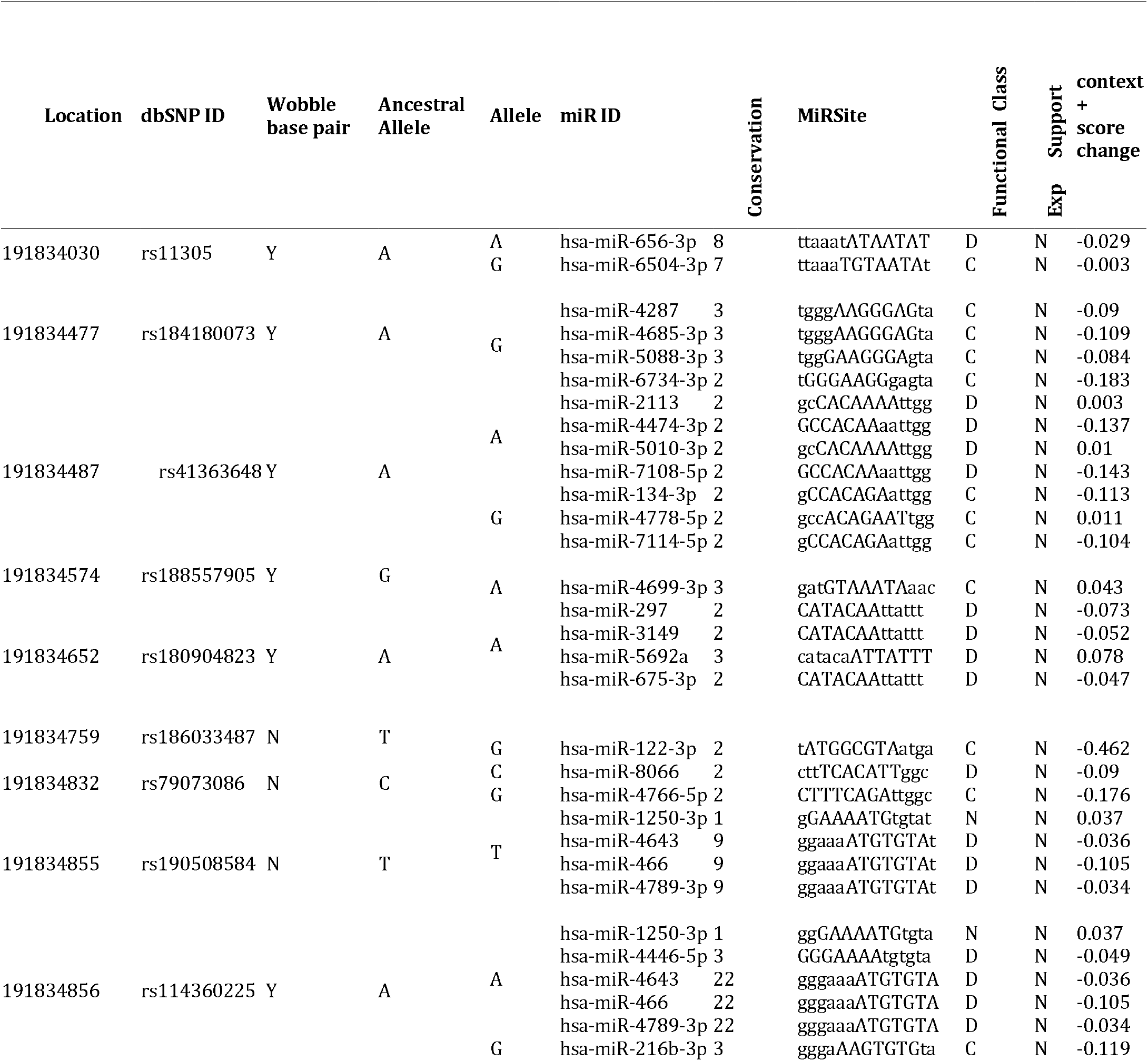
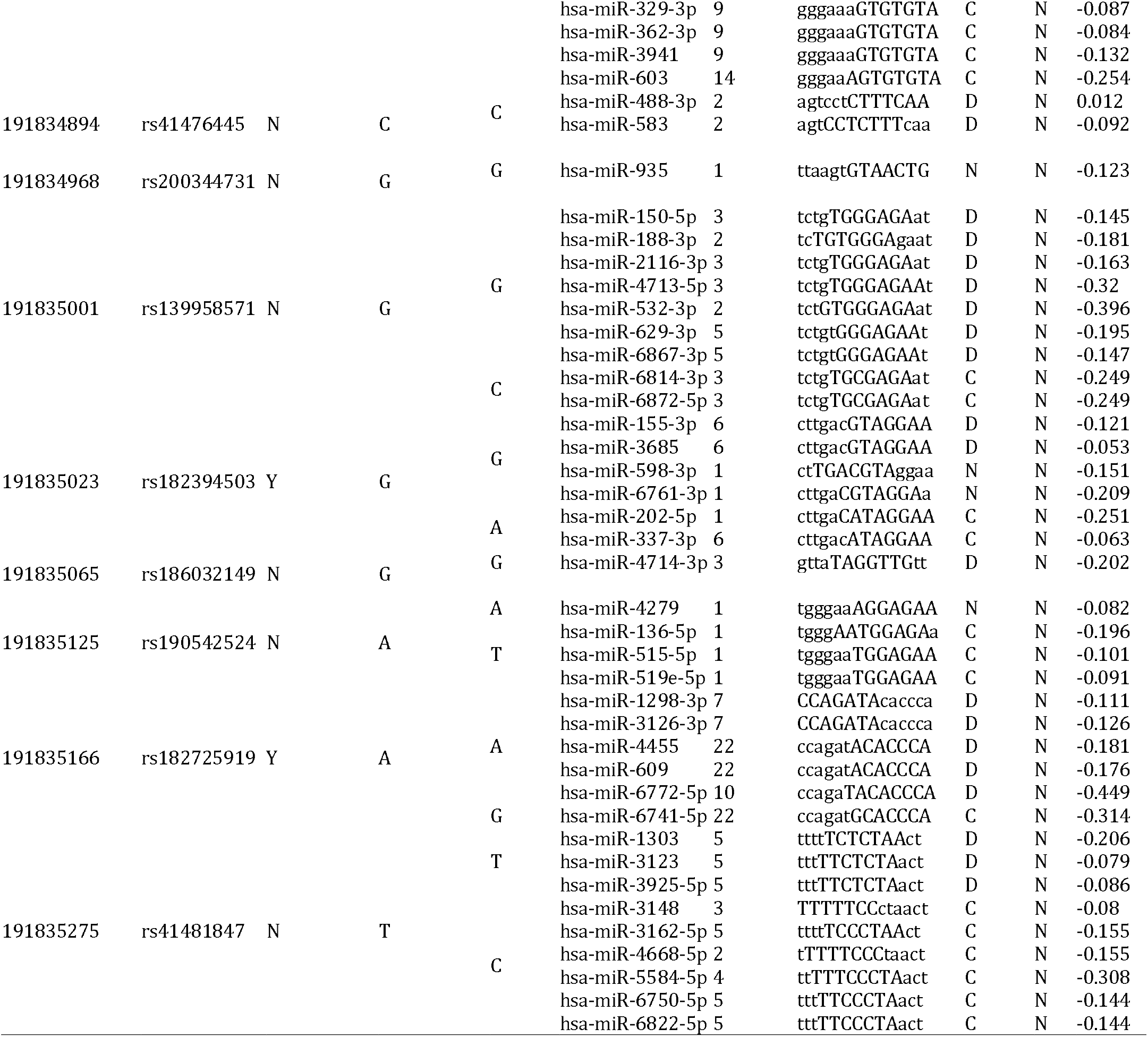
Location, alleles, miRNA ID, miRNA binding site and function of the 3 UTR from PolymiRTS analysis.

**Table 2.**
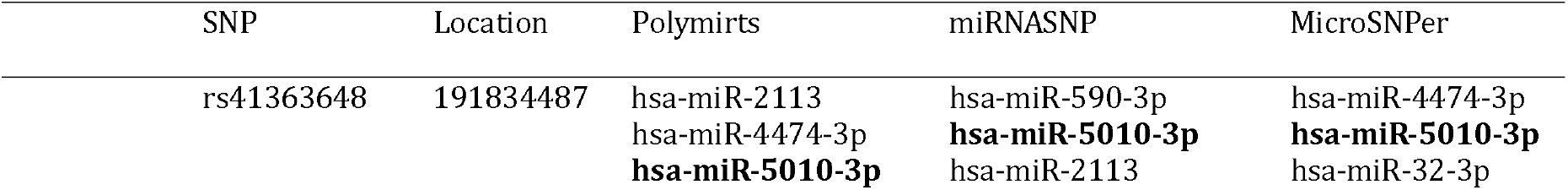
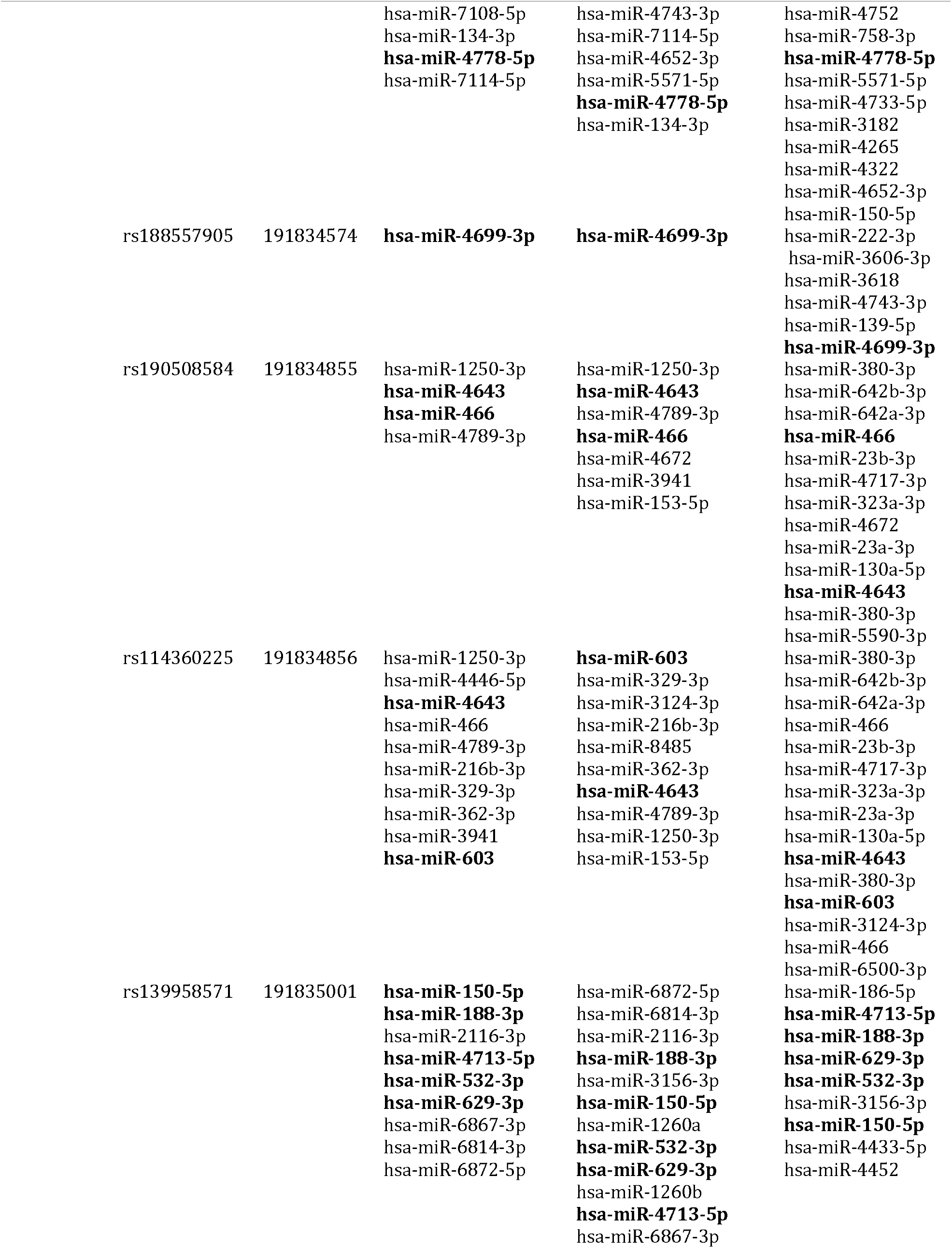
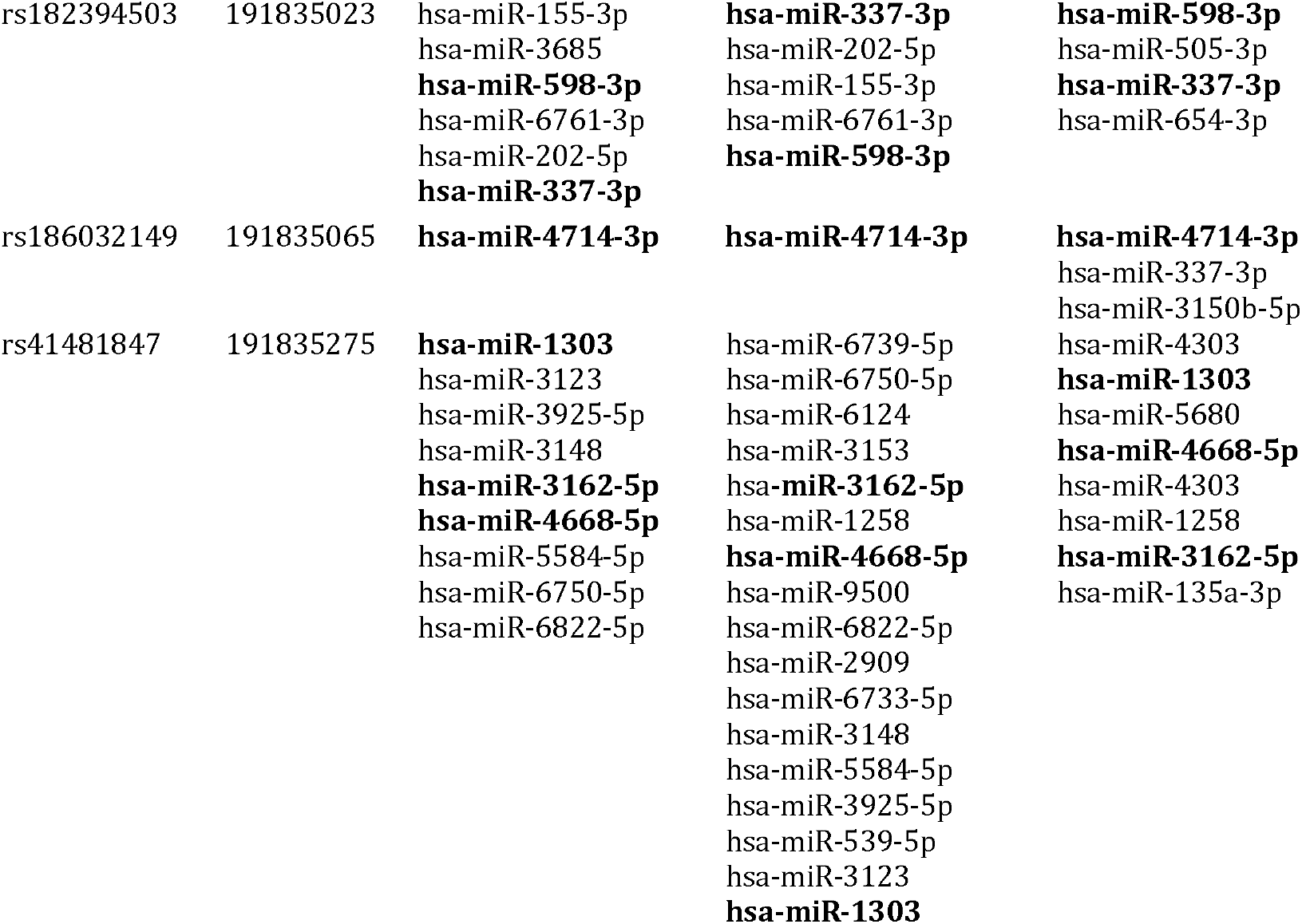
Association of the miRNA with 3 UTR SNPs predicted through PolymiRTS, MiRNASNP, and MicroSNIper.

**Figure 2.**
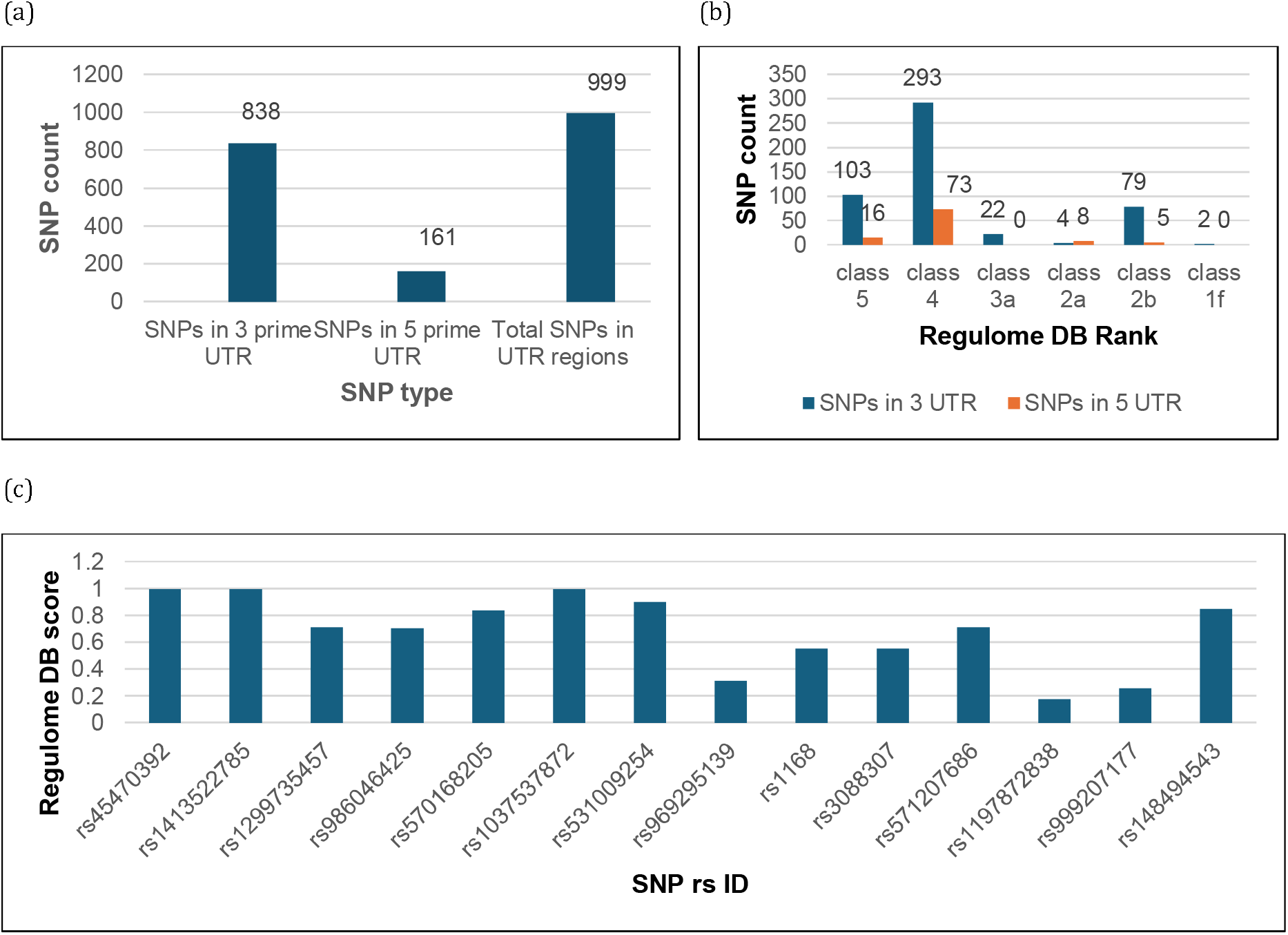
(a) Total count of 3⍰ and 5⍰ UTR variants of STAT1 recovered from NCBI; (b) annotation and scoring of UTR SNPs obtained through Regulome DB; (c) Regulome DB scores of SNPs annotated as, 1f and 2a.

### 3.2. Association of UTR SNPs with miRNAs

The influence of 3’ UTR SNPs obtained from NCBI on miRNA was assessed using PolymiRTS, miRNASNP, and MicroSNIper. The objective of utilizing PolymiRTS was to identify SNPs which affect miRNA target regions by or disrupting or creating a new miRNA target regions. Conservation scores, ancestral alleles, altered alleles, context, and score alterations are shown (Table 1). The role of SNP at the PolymiRTS has been classified into four categories. D represents the disruption of a conserved miRNA site, N signifies the disruption of non-conserved miRNA, C indicates the generation of novel miRNA, and O is employed when ancestral alleles remain undetermined. PolymiRTS identified the correlation of 17 variations with miRNAs, accompanied by disruption scores. Twelve SNPs were identified that disrupt the conserved region of miRNA, with scores ranging from 2 to 10. Five SNPs were identified that disrupt the non-conserved miRNA, exhibiting a score range of 0–16. Twelve SNPs contributed to the synthesis of novel miRNA.

MiRNASNP is used to determine the losses and gains of miRNAs. According to miRNASNP, 40 SNPs contributed to the loss of miRNA target sites, whereas 30 SNPs resulted in the gain of target sites. A total of 29 SNPs were modifying miRNA through both gain and loss of target sites. Supplementary Table 5. The overall findings from these three tools revealed that eight SNPs are functionally significant, with the greatest probability of modifying the miRNA target sequence. The miRNAs that were common across all tools are highlighted in bold font (Table 2).

### 3.4. Impact of UTR SNPs on Secondary Structure of mRNA

A total of 22 SNPs (6 in the 3’ UTR and 8 in the 5’ UTR regions) were classified as ≤2a (2a and 1f) by RegulomeDB. Additionally, the 8 SNPs that modify miRNA binding sites, identified across the three databases (PolymiRTS, miRNASNP, and MicroSNIper), underwent further analysis using RNAfold to assess their effects on mRNA structure. RNAfold analysis indicated an increase in minimum free energy (MFE) and destabilization of mRNA structure in both wild-type and mutant-type SNPs for 14 out of 22 SNPs, while the remaining 8 SNPs either did not induce significant alterations in MFE and mRNA structure or resulted in decreased MFE with stabilization of the secondary structure. Among the 14 SNPs, 2 resided in the 3⍰ UTR (Table 3), 3 were situated in the 5⍰ UTR (Table 4), and 1 SNP exhibited relationships with miRNA, being consistently recognized across the three databases (PolymiRTS, miRNASNP, and MicroSNIper) (Table 5). A substantial rise in energy was detected for the SNPs in the 3’ UTR, with rs1197872838 and rs3088307 showing ΔG values rising from –21.20 (kcal/mol) to –19.00 (kcal/mol) and from – 50.40 (kcal/mol) to –47.10 (kcal/mol), respectively. This indicates that the SNPs destabilize the mRNA structure. Other SNPs exhibit either marginal or no variation in energy, with minimal or no changes to the secondary structure (table 3).Significant alterations in ΔG were seen in three SNPs located in the 5⍰ UTR: rs45470392, rs1413522785, and rs531009254. The ΔG ranged from –40.90 to –38.80 (kcal/mol), –41.90 to –41.40 (kcal/mol), and –35.20 to – 34.10 (kcal/mol), respectively. Consequently, the mRNA structure is destabilized due to the elevated energy (Table 4). A significant alteration in mutant structures is noted, particularly in rs1413522785 and rs531009254. A reduction in the MFE of the mutant mRNA was noted in rs1299735457 (from –43.10 kcal/mol to –43.80 kcal/mol) and rs1037537872 (from –39.00 kcal/mol to –40.30 kcal/mol), indicating structural stability. No alteration in the minimal free energy for the remaining three SNPs (rs986046425, rs570168205, and rs969295139).

**Table 3.**
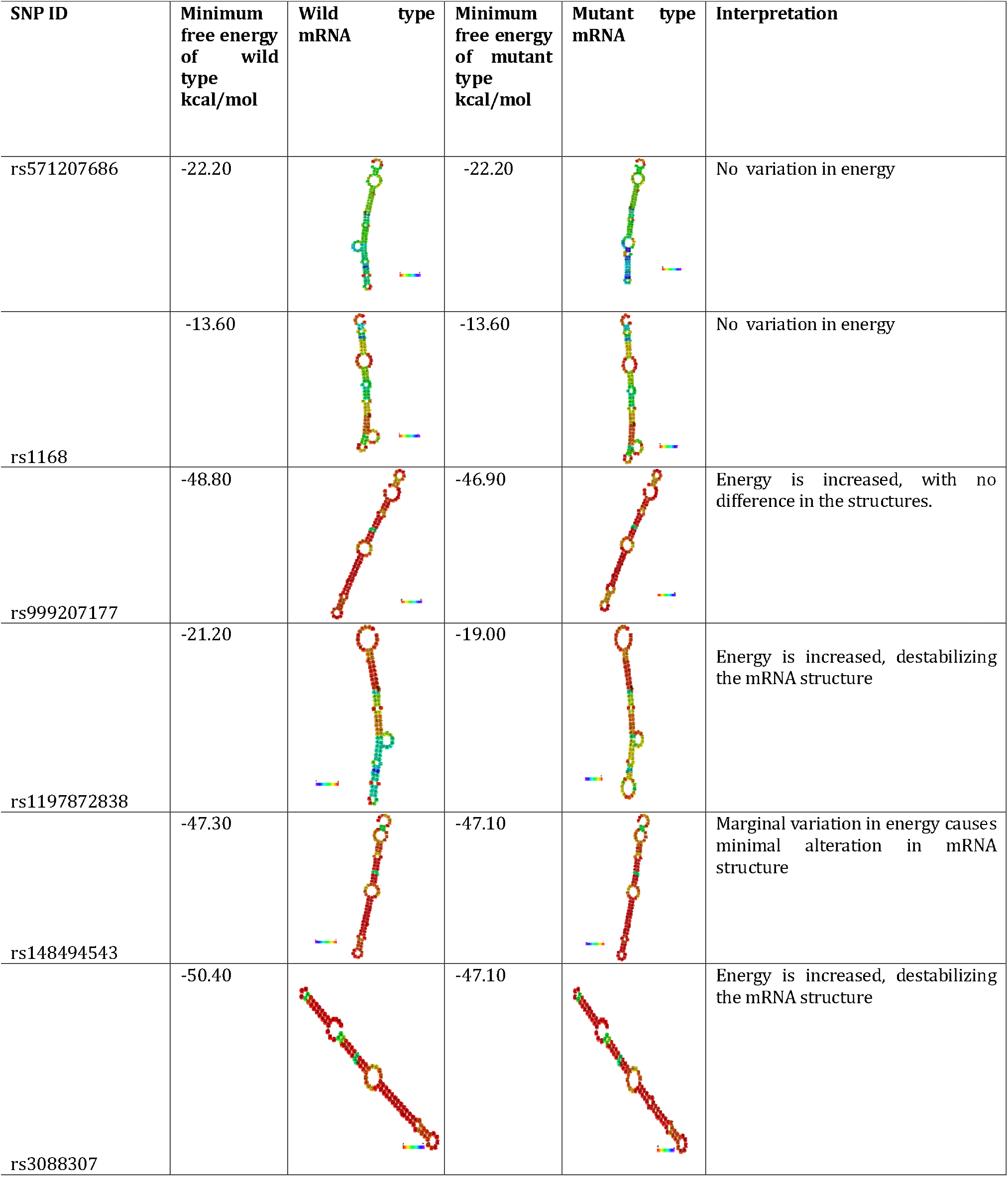
Comparison of structures of wild and mutated mRNA of 3⍰ UTR SNPs through RNAfold.

| SNP ID | Minimum free energy of wild type kcal/mol | Wild mRNA type | Minimum free energy of mutant type kcal/mol | Mutant mRNA type | Interpretation |
| --- | --- | --- | --- | --- | --- |
| rs571207686 | -22.20 |  | -22.20 |  | No variation in energy |
| rs1168 | -13.60 |  | -13.60 |  | No variation in energy |
| rs999207177 | -48.80 |  | -46.90 |  | Energy is increased, with no difference in the structures. |
| rs1197872838 | -21.20 |  | -19.00 |  | Energy is increased, destabilizing the mRNA structure |
| rs148494543 | -47.30 |  | -47.10 |  | Marginal variation in energy causes minimal alteration in mRNA structure |
| rs3088307 | -50.40 |  | -47.10 |  | Energy is increased, destabilizing the mRNA structure |

**Table 4.**
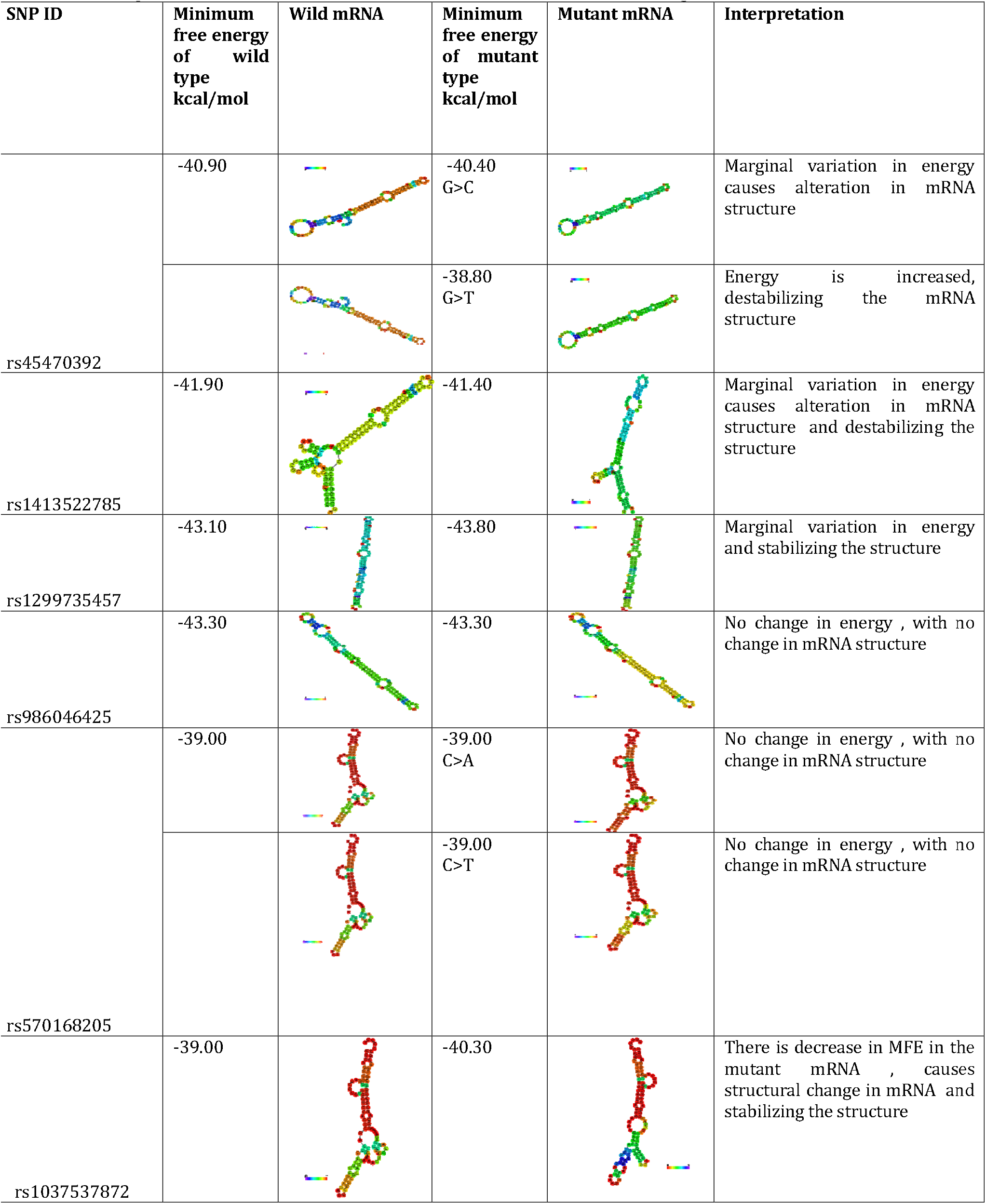
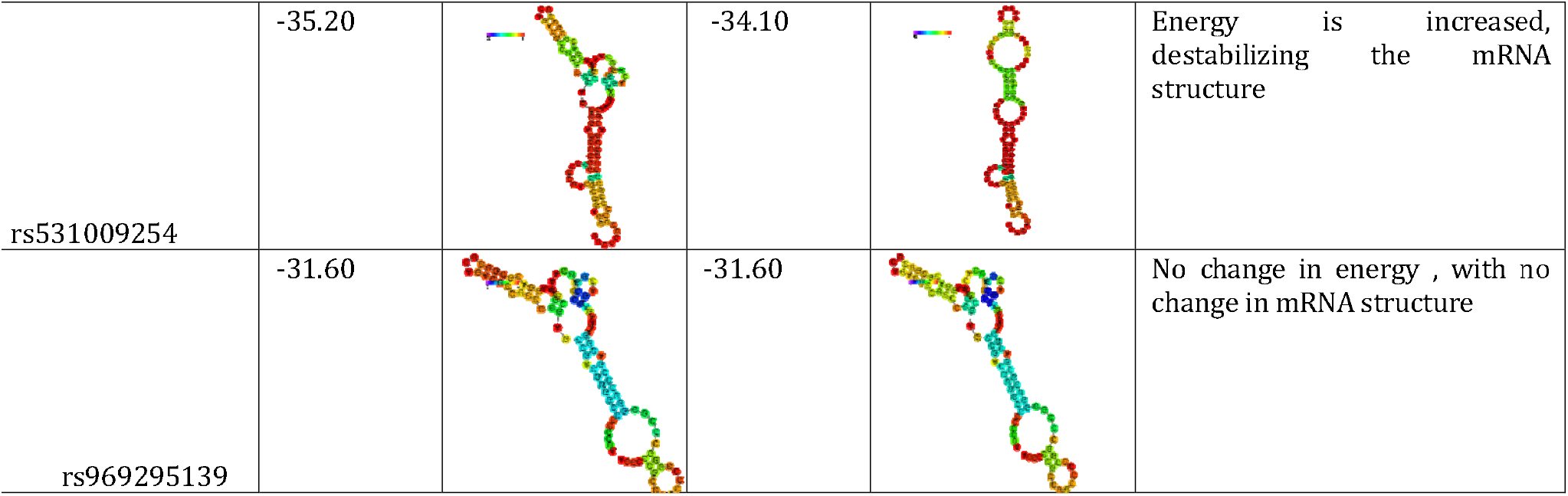
Comparison of structures of wild and mutated mRNA of 5⍰ UTR SNPs through RNAfold.

| SNP ID | Minimum free energy of wild type kcal/mol | Wild mRNA | Minimum free energy of mutant type kcal/mol | Mutant mRNA | Interpretation |
| --- | --- | --- | --- | --- | --- |
| rs45470392 | -40.90 |  | -40.40<br>G>C |  | Marginal variation in energy causes alteration in mRNA structure |
|  |  |  | -38.80<br>G>T |  | Energy is increased, destabilizing the mRNA structure |
| rs1413522785 | -41.90 |  | -41.40 |  | Marginal variation in energy causes alteration in mRNA structure and destabilizing the structure |
| rs1299735457 | -43.10 |  | -43.80 |  | Marginal variation in energy and stabilizing the structure |
| rs986046425 | -43.30 |  | -43.30 |  | No change in energy , with no change in mRNA structure |
| rs570168205 | -39.00 |  | -39.00<br>C>A |  | No change in energy , with no change in mRNA structure |
|  |  |  | -39.00<br>C>T |  | No change in energy , with no change in mRNA structure |
| rs1037537872 | -39.00 |  | -40.30 |  | There is decrease in MFE in the mutant mRNA , causes structural change in mRNA and stabilizing the structure |
| rs531009254 | -35.20 |  | -34.10 |  | Energy is increased, destabilizing the mRNA structure |
| rs969295139 | -31.60 |  | -31.60 |  | No change in energy , with no change in mRNA structure |

**Table 5.**
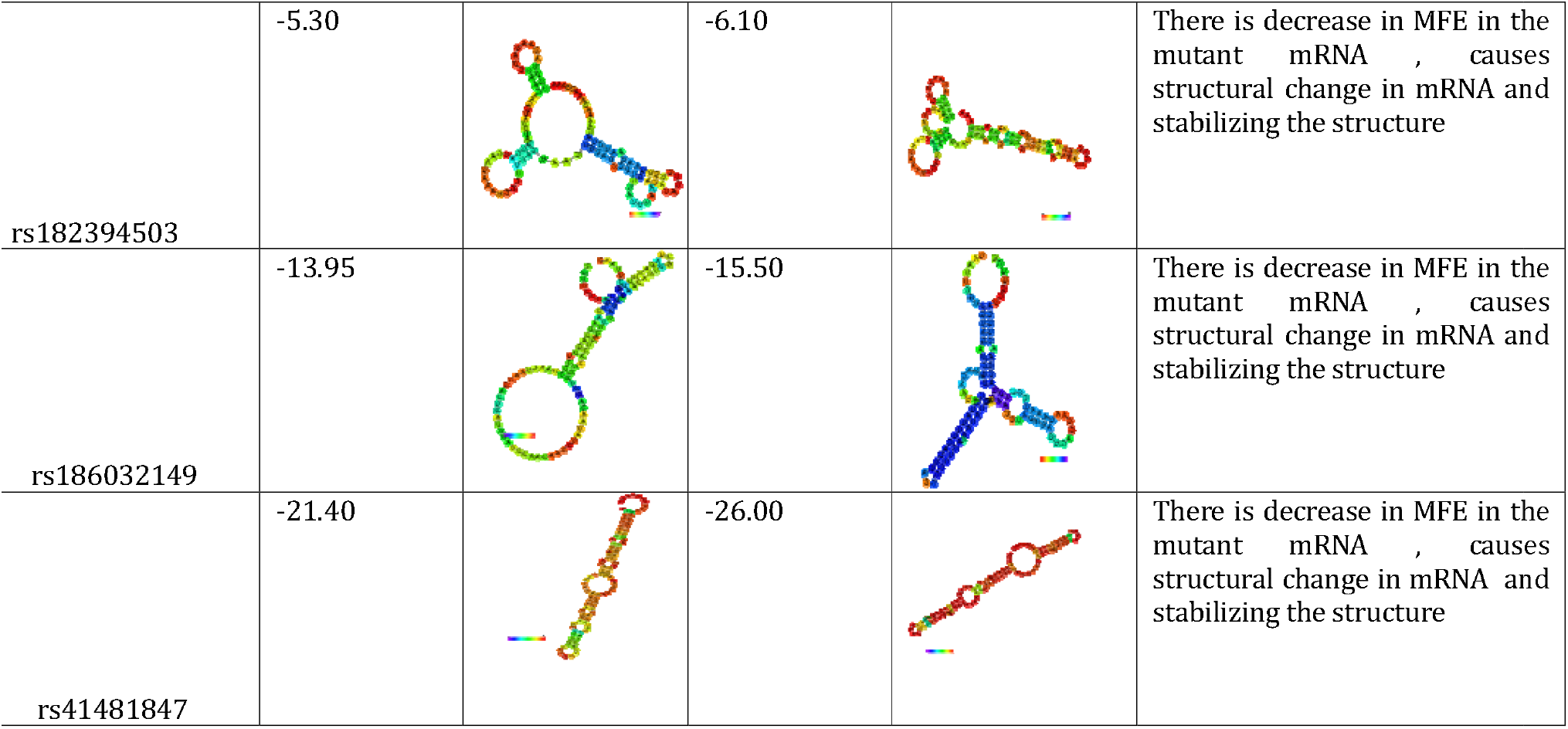
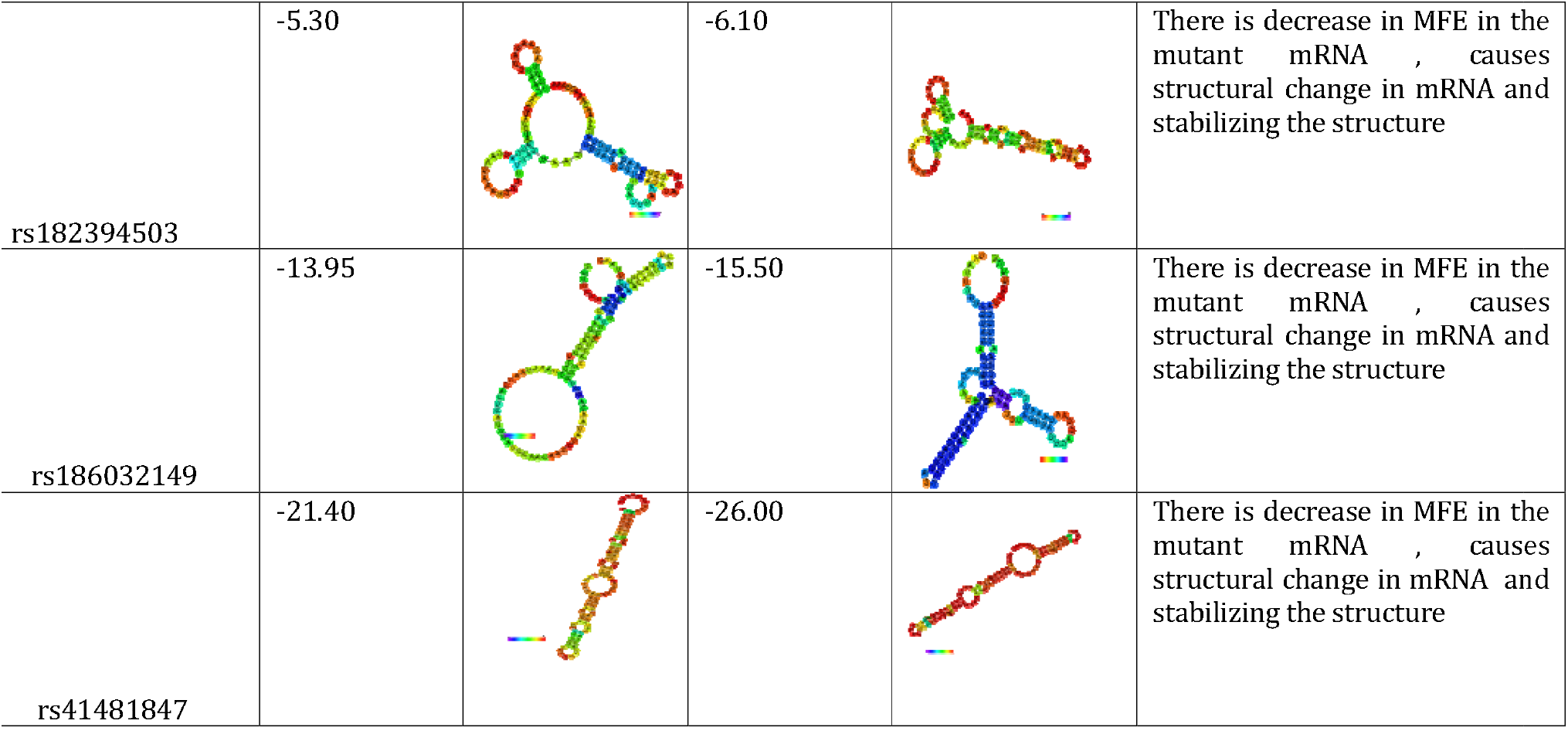
Comparison of structures of wild and mutated mRNA of functionally significant 3 UTR SNPs commonly predicted by PolymiRTS, MiRNASNP, and MicroSNIper.

| SNP ID | Minimum free energy of wild type kcal/mol | Wild mRNA | Minimum free energy of mutant type kcal/mol | Mutant mRNA | Interpretation |
| --- | --- | --- | --- | --- | --- |
| rs114360225 | -16.70 |  | -17.20 |  | There is decrease in MFE in the mutant mRNA , causes structural change in mRNA and stabilizing the structure |
| rs139958571 | -10.40 |  | -10.40 |  | No change in energy , with no change in mRNA structure |
| rs41363648 | -10.30 |  | -10.30 |  | No change in energy , with no change in mRNA structure |
| rs188557905 | -13.90 |  | -13.90 |  | No change in energy , with change in mRNA structure |
| rs190508584 | -17.00 |  | -16.70 |  | Energy is increased, destabilizing the mRNA structure |
| rs182394503 | -5.30 |  | -6.10 |  | There is decrease in MFE in the mutant mRNA , causes structural change in mRNA and stabilizing the structure |
| rs186032149 | -13.95 |  | -15.50 |  | There is decrease in MFE in the mutant mRNA , causes structural change in mRNA and stabilizing the structure |
| rs41481847 | -21.40 |  | -26.00 |  | There is decrease in MFE in the mutant mRNA , causes structural change in mRNA and stabilizing the structure |

Among the 8 SNPs related with miRNA, only one SNP (rs190508584) showed an increase in minimal free energy from –17.00 kcal/mol to –16.70 kcal/mol, resulting in destabilization of the mRNA structure. The remaining SNPs either result in reduced or no alteration in MFE.

### 3.5. Effect of 5′ UTR SNPs on Transcription Binding Factors

A total of eight 51l UTR SNPs annotated as ≤2a were selected for transcription factor binding analysis. Only one SNP, rs45470392, was predicted to alter transcription binding sites

### 3.6. Identification of cancer-associated SNPs

A total of 22 SNPs in the UTR (6 in 31l UTR and 8 in 51l UTR regions) were annotated as ≤2a (2a and 1f) by RegulomeDB, as well as the 8 functionally significant SNPs identified by the 3 databases (PolymiRTS, miRNASNP, and MicroSNIper) were further screened for their oncogenic potential using Cscape. Five polymorphisms were predicted as oncogenic with a score > 0.5. rs188557905, rs1413522785, rs1220766131, rs1168, and rs999207177 (Table 6).

**Table 6.** The oncogenic SNPs identified by C scape.

| SNP ID | Chromosomal location | Cscape score |
| --- | --- | --- |
| rs188557905 | 2,190969848,C,T | 0.567440 |
| rs1220766131 | 2, 191029618,T,A | 0.690487 |
| rs1413522785 | 2,191014053,G,A | 0.580640 |
| rs1168 | 2,190965051,C,T | 0.628264 |
| rs999207177 | 2,190964663,A,T | 0.676795 |

## Discussion

In the present study, we utilized the web tools PolymiRTS, miRNASNP, and MicroSNIper to ascertain the correlation of SNPs with miRNA. PolymiRTS was utilized to identify alterations in miRNA binding sites of *STAT1* due to 3 UTR SNPs. The integrated findings from these methods indicated that eight SNPs (rs41363648, rs188557905, rs190508584, rs114360225, rs139958571, rs182394503, rs41481847, and rs186032149) have the potential to modify microRNA targets within the 3’ UTR region.

The PolymiRTS tool found the same mutation at the same locus with SNP ID rs41363648 (A>A/G) in seven miRNAs: hsa-miR-2113, hsa-miR-4474-3p, hsa-miR-5010-3p, hsa-miR-7108-5p, hsa-miR-134-3p, hsa-miR-4778-5p, and hsa-miR-7114-5p. SNP ID rs41363648 was found to disrupt the conserved region of the 3’ UTR for hsa-miR-2113, hsa-miR-4474-3p, hsa-miR-5010-3p, and hsa-miR-7108-5p, with context scores of 0.003, –0.137, 0.01, and –0.143, respectively. Additionally, this SNP can generate new miRNA sites, hsa-miR-134-3p, hsa-miR-4778-5p, and hsa-miR-7114-5p, with scores of –0.113, 0.011, and –0.104, respectively. A higher context score correlated with increased structural change and disruption in miRNA. Table 1 enumerates the additional SNPs together with their impacts on miRNA binding sites and the corresponding miRNAs, accompanied by the context score.

A prior study revealed that gene interactions and FOXC2 expression were significantly influenced by SNPs located at the 5⍰ and 3⍰ regions. The three miRNAs, hsa-miR-6886-5p, hsa-miR-6886-5p, and hsa-miR-6720-3p, were significantly influenced by rs201118690, rs6413505, and rs201914560, respectively [27].

Additionally, we identified 17 miRNAs associated with the 8 SNPs that were consistently predicted by PolymiRTS, miRNASNP, and MicroSNIper (table 2). Consequently, SNPs in the STAT1 gene may modify the binding sites of miRNA, substantially influencing gene expression and contributing to cancer and other diseases [12]. Consistent with our findings, the identified 17 miRNAs were linked to multiple cancer types. Hsa-mir-466 is recognized as a tumor suppressor and a prognostic indicator in colorectal cancer [27], and it also inhibits tumor cell proliferation in osteosarcoma (28) and prostate cancer [29]. Hsa-mir-150-5p functions as a tumor suppressor, inhibiting cancer cell proliferation and migration across several cancer types [30–35], whereas hsa-mir-532-3p has been identified as a promoter of hepatocellular carcinoma [36].

Hsa-mir-337-3p suppresses gastric cancer [37] and neuroplastoma via matrix metalloproteinase 14 [38]. It inhibits the growth and invasion of hepatocellular carcinoma cells by targeting JAK2 [39]. Hsa-mir-603 functions as an oncogene and was upregulated in both glioma [40] and osteosarcoma [41], whereas it inhibited hexokinase-2 in ovarian cancer cells [42]. The expression of hsa-miR-5010-3p, hsa-miR-4778-5p, hsa-miR-188-3p, and hsa-miR-4699-3p was shown to be down-regulated in stage II colon cancer according to miRNA expression analysis, whereas hsa-miR-4713-5p and hsa-miR-4668-5p were discovered to be up-regulated [43]. Prior studies identified a correlation between multiple cancer types and hsa-miR-5010-3p (44), hsa-miR-4778-5p (45), hsa-miR-4643(46– 49), hsa-miR-629-3p (46, 50, 51), hsa-miR-598-3p (44, 46, 47), hsa-miR-4714-3p[44–47], and hsa-miR-1303[48– 51].

The present investigation a sum of 838 SNPs from the 3⍰ UTR and 161 SNPs from the 5⍰ UTR from NCBI. Upon performing a RegulomeDB analysis, it was determined that 14 SNPs in the UTR (six in the 3’ UTR and eight in the 5’ UTR sections) were classified as ≤2a, while the remainder SNPs were categorized from 3b to 6.

A prior study examining the functional relevance of UTR SNPs in the regulation of the PRKCI gene identified 6 SNPs categorized as 2a and 25 as 2b, while the rest SNPs were classified into different categories ranging from 3a to 6 [52]. Another investigation discovered SNPs within non-coding variations associated with the development of coronary artery disorders. A total of 1200 SNPs were examined, of which 858 had scores ranging from 1 to 6. Additionally, 97 of 858 individuals obtained scores < 3 [53].

The influence of functionally significant SNPs from RegulomeDB and miRNA associations on mRNA structure was assessed using RNAfold. Two SNPs from the 3⍰ UTR (rs1197872838 and rs3088307) and five from the 5⍰ UTR (rs45470392, rs1413522785, and rs531009254) destabilized the overall mRNA structure, resulting in a significant increase in the minimum free energy (MFE). A reduction in the MFE of the mutant mRNA was noted in the 5⍰ UTR SNPs, rs1299735457 and rs1037537872, accompanied by structural stability. Among the 8 SNPs linked to miRNA, rs190508584 exhibits a destabilized structure with elevated MFE.

Research indicates that the modulation and control of mRNA structural stability are crucial for optimal protein synthesis [54]. Prior research linked SNPs that destabilize mRNA with several illnesses. A case-control research identified a 3’-UTR SNP rs929271 on leukaemia inhibitory factor (LIF) that destabilizes the mRNA structure, correlating with heightened susceptibility to schizophrenia [55].

A separate research study identified a regulatory 3’-UTR SNP (rs2229611) in the G6PC1 gene that influences mRNA stability, as determined by RNAfold, and is correlated with a more aggressive disease pattern in GSD-Ia patients. [56]

The 5⍰UTR SNPs designated as 2a and 1f were examined using SNPinfo to assess their influence on the formation or disruption of DNA binding sites for transcription factors. The SNP rs45470392 was found to modify transcription factor binding sites; it possesses a Regulome DB class of 2a and a score of 1. It also contributes to the destabilization of the mRNA structure, indicating its potential correlation with diseases.

A published research report identified rs2728127 at the osteopontin promoter on transcription binding sites. This SNP correlates with a reduction in the quantity of transcription factors [57]

To investigate the potential oncogenic effects of the target SNPs, we utilized the Cscape tool and identified that rs188557905, rs1413522785, rs1220766131, rs1168, and rs999207177 possess oncogenic scores over 0.5. Only Rs188557905 was documented in ClinVar (variation ID: 333255) as of unknown importance, linked to Mendelian susceptibility to mycobacterial illnesses resulting from partial STAT1 impairment; additional SNPs were not recorded in Clinvar. Although the bioinformatics methods employed are reliable, the absence of experimental validation restricts the ability to draw definitive conclusions regarding the clinical significance of the identified UTR mutations. Experimental validation, encompassing both in vitro and in vivo studies, is necessary to corroborate these in silico predictions. Recognizing polymorphisms in the UTR of the STAT1 gene as possible modulators of gene expression paves the way for novel therapeutic techniques aimed at these regions in cancer and immune-related hereditary disorders. Future experimental investigations are required to corroborate these bioinformatics findings and evaluate their viability as biomarkers for precision medicine.

## Acknowledgment

We sincerely thank all participants for their important cooperation and scientific contributions. We are also grateful to Prof. Mohamedahmed Salih Hassan for his experience and guidance in bioinformatics, as well as his insightful consultation and overall support throughout the development of this article.

