## supplementary tables for "Bioinformatics Analysis of Non-Coding UTR Variants in *STAT1* Reveals Disruption of miRNA Binding, mRNA Stability, and Oncogenic Potential"

Table S1: list of SNPs in the 5 prime UTR of STAT1 gene from NCBI.

| SNP ID | Location |
| --- | --- |
| rs1695029891 | 1.91E+08 |
| rs1467622177 | 1.91E+08 |
| rs1559026214 | 1.91E+08 |
| rs960205570 | 1.91E+08 |
| rs567434849 | 1.91E+08 |
| rs531511204 | 1.91E+08 |
| rs963769760 | 1.91E+08 |
| rs1191503043 | 1.91E+08 |
| rs1405565781 | 1.91E+08 |
| rs1372817347 | 1.91E+08 |
| rs1695031835 | 1.91E+08 |
| rs765693257 | 1.91E+08 |
| rs1695032215 | 1.91E+08 |
| rs1695032397 | 1.91E+08 |
| rs1695032581 | 1.91E+08 |
| rs1695032760 | 1.91E+08 |
| rs1459026520 | 1.91E+08 |
| rs1695033151 | 1.91E+08 |
| rs549949329 | 1.91E+08 |
| rs1695033521 | 1.91E+08 |
| rs1405849643 | 1.91E+08 |
| rs1695033898 | 1.91E+08 |
| rs1695034076 | 1.91E+08 |
| rs571354699 | 1.91E+08 |
| rs1695034439 | 1.91E+08 |
| rs1695034818 | 1.91E+08 |
| rs1238278593 | 1.91E+08 |
| rs1180393338 | 1.91E+08 |
| rs1695035302 | 1.91E+08 |
| rs909295245 | 1.91E+08 |
| rs750862823 | 1.91E+08 |
| rs1695035837 | 1.91E+08 |
| rs772559222 | 1.91E+08 |
| rs1490002122 | 1.91E+08 |
| rs1695036568 | 1.91E+08 |
| rs1695302734 | 1.91E+08 |
| rs902278776 | 1.91E+08 |
| rs1695303105 | 1.91E+08 |
| rs1408491477 | 1.91E+08 |
| rs1695303465 | 1.91E+08 |
| rs1695303635 | 1.91E+08 |
| rs372299811 | 1.91E+08 |
| rs1695303957 | 1.91E+08 |
| rs1220675431 | 1.91E+08 |
| rs999306857 | 1.91E+08 |
| rs189188567 | 1.91E+08 |
| rs1279334374 | 1.91E+08 |
| rs1695304839 | 1.91E+08 |
| rs1490866036 | 1.91E+08 |
| rs1695305172 | 1.91E+08 |
| rs1240818117 | 1.91E+08 |
| rs1215740346 | 1.91E+08 |
| rs11549697 | 1.91E+08 |
| rs1286840473 | 1.91E+08 |
| rs1053528132 | 1.91E+08 |
| rs1201881287 | 1.91E+08 |
| rs938419162 | 1.91E+08 |
| rs1559028129 | 1.91E+08 |
| rs1695306854 | 1.91E+08 |
| rs1282245963 | 1.91E+08 |
| rs1223531891 | 1.91E+08 |
| rs1369144548 | 1.91E+08 |
| rs1695308019 | 1.91E+08 |
| rs1695308188 | 1.91E+08 |
| rs1695308336 | 1.91E+08 |
| rs545742697 | 1.91E+08 |
| rs1695308731 | 1.91E+08 |
| rs1429996774 | 1.91E+08 |
| rs1345553432 | 1.91E+08 |
| rs1320123706 | 1.91E+08 |
| rs1483073172 | 1.91E+08 |
| rs1695309659 | 1.91E+08 |
| rs1056003060 | 1.91E+08 |
| rs1481434204 | 1.91E+08 |
| rs1198263292 | 1.91E+08 |
| rs1695310373 | 1.91E+08 |
| rs1695310559 | 1.91E+08 |
| rs1695310732 | 1.91E+08 |
| rs118149197 | 1.91E+08 |
| rs144940940 | 1.91E+08 |
| rs947541007 | 1.91E+08 |
| rs1317848379 | 1.91E+08 |
| rs993374058 | 1.91E+08 |
| rs556645752 | 1.91E+08 |
| rs1574678292 | 1.91E+08 |
| rs994096848 | 1.91E+08 |
| rs1035535139 | 1.91E+08 |
| rs55724546 | 1.91E+08 |
| rs1574678306 | 1.91E+08 |
| rs1695351743 | 1.91E+08 |
| rs952804507 | 1.91E+08 |
| rs1574678321 | 1.91E+08 |
| rs1325660325 | 1.91E+08 |
| rs45470392 | 1.91E+08 |
| rs1695353681 | 1.91E+08 |
| rs1413522785 | 1.91E+08 |
| rs1299735457 | 1.91E+08 |
| rs986046425 | 1.91E+08 |
| rs1574678353 | 1.91E+08 |
| rs1490047123 | 1.91E+08 |
| rs1269450206 | 1.91E+08 |
| rs1695355339 | 1.91E+08 |
| rs1559028449 | 1.91E+08 |
| rs1211595473 | 1.91E+08 |
| rs1695355864 | 1.91E+08 |
| rs147950001 | 1.91E+08 |
| rs965520149 | 1.91E+08 |
| rs1240024048 | 1.91E+08 |
| rs1331407038 | 1.91E+08 |
| rs2125114909 | 1.91E+08 |
| rs564075337 | 1.91E+08 |
| rs1695357404 | 1.91E+08 |
| rs1263141636 | 1.91E+08 |
| rs968480409 | 1.91E+08 |
| rs978953371 | 1.91E+08 |
| rs1695358116 | 1.91E+08 |
| rs934985471 | 1.91E+08 |
| rs989136304 | 1.91E+08 |
| rs1695358766 | 1.91E+08 |
| rs914987046 | 1.91E+08 |
| rs1384678089 | 1.91E+08 |
| rs1695359379 | 1.91E+08 |
| rs528057545 | 1.91E+08 |
| rs1210087528 | 1.91E+08 |
| rs1574678448 | 1.91E+08 |
| rs947858016 | 1.91E+08 |
| rs1695360357 | 1.91E+08 |
| rs568025055 | 1.91E+08 |
| rs560084350 | 1.91E+08 |
| rs1695361247 | 1.91E+08 |
| rs371724653 | 1.91E+08 |
| rs1466106799 | 1.91E+08 |
| rs1445643161 | 1.91E+08 |
| rs1026959454 | 1.91E+08 |
| rs1276743117 | 1.91E+08 |
| rs977004227 | 1.91E+08 |
| rs2125115118 | 1.91E+08 |
| rs769850162 | 1.91E+08 |
| rs1181864140 | 1.91E+08 |
| rs1695363439 | 1.91E+08 |
| rs922842738 | 1.91E+08 |
| rs1007000950 | 1.91E+08 |
| rs1574678528 | 1.91E+08 |
| rs530356200 | 1.91E+08 |
| rs1695364453 | 1.91E+08 |
| rs548761843 | 1.91E+08 |
| rs1695364832 | 1.91E+08 |
| rs888550057 | 1.91E+08 |
| rs1460039243 | 1.91E+08 |
| rs1695365629 | 1.91E+08 |
| rs886055382 | 1.91E+08 |
| rs1695366340 | 1.91E+08 |
| rs570168205 | 1.91E+08 |
| rs1037537872 | 1.91E+08 |
| rs531009254 | 1.91E+08 |
| rs1695367190 | 1.91E+08 |
| rs1695367380 | 1.91E+08 |
| rs1695367559 | 1.91E+08 |
| rs1695367707 | 1.91E+08 |
| rs1695367898 | 1.91E+08 |
| rs969295139 | 1.91E+08 |

Table S2: list of SNPs in the 3 prime UTR of STAT1 gene from NCBI.

| SNP ID | Location |
| --- | --- |
| rs1691080429 | 2:190964648 |
| rs944203048 | 2:190964650 |
| rs1691080512 | 2:190964654 |
| rs578071388 | 2:190964655 |
| rs902756687 | 2:190964657 |
| rs1691080679 | 2:190964658 |
| rs1691080718 | 2:190964659 |
| rs999207177 | 2:190964663 |
| rs1691080867 | 2:190964670 |
| rs1691080916 | 2:190964671 |
| rs148494543 | 2:190964674 |
| rs1691082945 | 2:190964680 |
| rs1691083002 | 2:190964683 |
| rs1574628816 | 2:190964684 |
| rs1393979820 | 2:190964685 |
| rs3088307 | 2:190964686 |
| rs969015549 | 2:190964688 |
| rs1691083457 | 2:190964689 |
| rs978426115 | 2:190964693 |
| rs1691083645 | 2:190964703 |
| rs1272621503 | 2:190964706 |
| rs1175133175 | 2:190964709 |
| rs1691083792 | 2:190964710 |
| rs1691083972 | 2:190964716 |
| rs1574628849 | 2:190964717 |
| rs1691084170 | 2:190964726 |
| rs1691084227 | 2:190964728 |
| rs1691084309 | 2:190964729 |
| rs1691084386 | 2:190964730 |
| rs1245907638 | 2:190964732 |
| rs981897633 | 2:190964734 |
| rs142693004 | 2:190964737 |
| rs1384263900 | 2:190964740 |
| rs542532827 | 2:190964743 |
| rs1168861525 | 2:190964745 |
| rs146017620 | 2:190964748 |
| rs1691084957 | 2:190964749 |
| rs1691085014 | 2:190964751 |
| rs1691085055 | 2:190964752 |
| rs1445081648 | 2:190964753 |
| rs1691085179 | 2:190964754 |
| rs961912738 | 2:190964755 |
| rs1691085290 | 2:190964759 |
| rs972722568 | 2:190964761 |
| rs1691085460 | 2:190964762 |
| rs2124964534 | 2:190964766 |
| rs1691085557 | 2:190964767 |
| rs1691085609 | 2:190964768 |
| rs1691085654 | 2:190964771 |
| rs1398023780 | 2:190964774 |
| rs1325767181 | 2:190964777 |
| rs28930678 | 2:190964778 |
| rs1443115055 | 2:190964781 |
| rs1423926011 | 2:190964786 |
| rs1691086230 | 2:190964790 |
| rs922782733 | 2:190964794 |
| rs2124964602 | 2:190964795 |
| rs932821143 | 2:190964798 |
| rs750560218 | 2:190964800 |
| rs139926849 | 2:190964823 |
| rs1308984928 | 2:190964824 |
| rs1691086956 | 2:190964827 |
| rs571416970 | 2:190964829 |
| rs532137299 | 2:190964833 |
| rs547454699 | 2:190964842 |
| rs1691087221 | 2:190964845 |
| rs1691087325 | 2:190964849 |
| rs2124964686 | 2:190964853 |
| rs1691087428 | 2:190964854 |
| rs1373058535 | 2:190964855 |
| rs1278530373 | 2:190964858 |
| rs758675541 | 2:190964859 |
| rs1442463937 | 2:190964866 |
| rs1691087770 | 2:190964878 |
| rs1037029827 | 2:190964879 |
| rs897159627 | 2:190964881 |
| rs939462620 | 2:190964892 |
| rs1691088096 | 2:190964899 |
| rs1056539629 | 2:190964900 |
| rs1158994136 | 2:190964910 |
| rs1691088410 | 2:190964911 |
| rs1474902590 | 2:190964916 |
| rs1691088513 | 2:190964918 |
| rs1691088565 | 2:190964922 |
| rs1691088625 | 2:190964927 |
| rs1691088672 | 2:190964929 |
| rs868410365 | 2:190964934 |
| rs2124964787 | 2:190964937 |
| rs1165774038 | 2:190964938 |
| rs1422126533 | 2:190964939 |
| rs1267087229 | 2:190964940 |
| rs1691089042 | 2:190964941 |
| rs1198763312 | 2:190964942 |
| rs752964780 | 2:190964943 |
| rs1271634891 | 2:190964946 |
| rs902778052 | 2:190964948 |
| rs1285389439 | 2:190964953 |
| rs1691089427 | 2:190964957 |
| rs1164845107 | 2:190964958 |
| rs1392985321 | 2:190964959 |
| rs192670482 | 2:190964960 |
| rs1691089784 | 2:190964962 |
| rs1691089828 | 2:190964967 |
| rs1691089875 | 2:190964970 |
| rs200913982 | 2:190964978 |
| rs1222328384 | 2:190964981 |
| rs1691090359 | 2:190964988 |
| rs1044848643 | 2:190964989 |
| rs1691090458 | 2:190965004 |
| rs369150147 | 2:190965005 |
| rs1691090569 | 2:190965010 |
| rs1012769799 | 2:190965015 |
| rs1427402088 | 2:190965016 |
| rs536205722 | 2:190965017 |
| rs1480770453 | 2:190965020 |
| rs556104368 | 2:190965021 |
| rs1691091008 | 2:190965023 |
| rs1691091062 | 2:190965029 |
| rs1023179868 | 2:190965034 |
| rs1272104884 | 2:190965038 |
| rs143345038 | 2:190965040 |
| rs1000105374 | 2:190965041 |
| rs1691091463 | 2:190965045 |
| rs1691091512 | 2:190965048 |
| rs1272302640 | 2:190965050 |
| rs1168 | 2:190965051 |
| rs1691091759 | 2:190965052 |
| rs1323892661 | 2:190965053 |
| rs556506301 | 2:190965056 |
| rs1361164169 | 2:190965064 |
| rs1691092115 | 2:190965067 |
| rs1691092164 | 2:190965071 |
| rs955670740 | 2:190965075 |
| rs1691092369 | 2:190965076 |
| rs1691092417 | 2:190965077 |
| rs2124965007 | 2:190965078 |
| rs965812479 | 2:190965079 |
| rs1691092539 | 2:190965084 |
| rs976525545 | 2:190965086 |
| rs1691092712 | 2:190965088 |
| rs1691092851 | 2:190965098 |
| rs747337724 | 2:190965104 |
| rs1574629076 | 2:190965107 |
| rs146824155 | 2:190965111 |
| rs924065088 | 2:190965122 |
| rs545184536 | 2:190965123 |
| rs1574629087 | 2:190965125 |
| rs2124965079 | 2:190965132 |
| rs1243426905 | 2:190965133 |
| rs1691093261 | 2:190965135 |
| rs1379489171 | 2:190965136 |
| rs1242907769 | 2:190965138 |
| rs953889876 | 2:190965139 |
| rs1691093554 | 2:190965146 |
| rs1257447210 | 2:190965151 |
| rs868463796 | 2:190965165 |
| rs1201769718 | 2:190965168 |
| rs1306448620 | 2:190965170 |
| rs1691093807 | 2:190965172 |
| rs1691093849 | 2:190965174 |
| rs554209165 | 2:190965175 |
| rs915465134 | 2:190965176 |
| rs1691094000 | 2:190965181 |
| rs1380575153 | 2:190965185 |
| rs1691094096 | 2:190965187 |
| rs1691094132 | 2:190965188 |
| rs1416099500 | 2:190965189 |
| rs140488842 | 2:190965190 |
| rs1691094286 | 2:190965191 |
| rs941657708 | 2:190965193 |
| rs972941504 | 2:190965194 |
| rs918616783 | 2:190965197 |
| rs1381552086 | 2:190965198 |
| rs1691094516 | 2:190965202 |
| rs1691094590 | 2:190965209 |
| rs1691094633 | 2:190965214 |
| rs1296698533 | 2:190965216 |
| rs1691094721 | 2:190965218 |
| rs1691094816 | 2:190965228 |
| rs1398955907 | 2:190965229 |
| rs1691094988 | 2:190965245 |
| rs563090709 | 2:190965248 |
| rs1343050347 | 2:190965249 |
| rs1056845485 | 2:190965250 |
| rs1216429025 | 2:190965253 |
| rs895232493 | 2:190965254 |
| rs2124965216 | 2:190965261 |
| rs1691095309 | 2:190965262 |
| rs1446849892 | 2:190965263 |
| rs879641888 | 2:190965266 |
| rs1175127546 | 2:190965269 |
| rs1036156937 | 2:190965271 |
| rs769148496 | 2:190965272 |
| rs1691095594 | 2:190965278 |
| rs1691095636 | 2:190965281 |
| rs149993422 | 2:190965282 |
| rs867842021 | 2:190965283 |
| rs2124965259 | 2:190965285 |
| rs1691095790 | 2:190965294 |
| rs1691095822 | 2:190965296 |
| rs560939666 | 2:190965311 |
| rs1027603252 | 2:190965316 |
| rs1691096131 | 2:190965317 |
| rs1323288887 | 2:190965322 |
| rs1305062112 | 2:190965323 |
| rs1691096329 | 2:190965331 |
| rs575978803 | 2:190965332 |
| rs1691096499 | 2:190965333 |
| rs1691096659 | 2:190965334 |
| rs904352199 | 2:190965342 |
| rs1691096749 | 2:190965344 |
| rs1691096849 | 2:190965346 |
| rs1691096904 | 2:190965349 |
| rs1000137833 | 2:190965354 |
| rs1691097024 | 2:190965357 |
| rs1691097076 | 2:190965362 |
| rs1403234504 | 2:190965363 |
| rs1365452819 | 2:190965368 |
| rs1284671779 | 2:190965377 |
| rs543681256 | 2:190965379 |
| rs564968175 | 2:190965381 |
| rs1018656137 | 2:190965390 |
| rs1574629222 | 2:190965392 |
| rs1691097840 | 2:190965394 |
| rs1691097914 | 2:190965399 |
| rs1691097976 | 2:190965404 |
| rs2124965382 | 2:190965405 |
| rs1574629229 | 2:190965408 |
| rs1488434355 | 2:190965409 |
| rs1691098203 | 2:190965410 |
| rs1691098282 | 2:190965419 |
| rs1691098360 | 2:190965422 |
| rs1691098444 | 2:190965426 |
| rs779962342 | 2:190965433 |
| rs1691098741 | 2:190965436 |
| rs1183645904 | 2:190965441 |
| rs1255871629 | 2:190965446 |
| rs1691099296 | 2:190965454 |
| rs1029709527 | 2:190965459 |
| rs1489455527 | 2:190965460 |
| rs2124965439 | 2:190965467 |
| rs185595627 | 2:190965475 |
| rs190448557 | 2:190965476 |
| rs1691099826 | 2:190965477 |
| rs1691099886 | 2:190965480 |
| rs1017128964 | 2:190965482 |
| rs1238114169 | 2:190965485 |
| rs374737590 | 2:190965486 |
| rs1188858771 | 2:190965491 |
| rs1691100245 | 2:190965494 |
| rs962840379 | 2:190965499 |
| rs181808548 | 2:190965500 |
| rs2124965472 | 2:190965503 |
| rs918859474 | 2:190965507 |
| rs1380966191 | 2:190965508 |
| rs960904276 | 2:190965519 |
| rs990036357 | 2:190965521 |
| rs1691100818 | 2:190965525 |
| rs1383486621 | 2:190965526 |
| rs1691100941 | 2:190965528 |
| rs185221390 | 2:190965533 |
| rs1288629572 | 2:190965534 |
| rs992178675 | 2:190965535 |
| rs1434392052 | 2:190965537 |
| rs748591907 | 2:190965538 |
| rs1373433759 | 2:190965539 |
| rs1416328694 | 2:190965541 |
| rs1412193183 | 2:190965542 |
| rs116804530 | 2:190965548 |
| rs1199596477 | 2:190965549 |
| rs1691102148 | 2:190965552 |
| rs1691265517 | 2:190969154 |
| rs915074408 | 2:190969156 |
| rs1691265908 | 2:190969161 |
| rs946639566 | 2:190969165 |
| rs1029098266 | 2:190969168 |
| rs987641912 | 2:190969179 |
| rs1691267717 | 2:190969182 |
| rs2124972264 | 2:190969184 |
| rs1160249930 | 2:190969185 |
| rs1433022831 | 2:190969189 |
| rs1691268752 | 2:190969192 |
| rs1691269044 | 2:190969197 |
| rs1559001454 | 2:190969200 |
| rs1042683838 | 2:190969201 |
| rs1020840161 | 2:190969202 |
| rs1691270427 | 2:190969211 |
| rs902405364 | 2:190969212 |
| rs1259793627 | 2:190969217 |
| rs1186439062 | 2:190969219 |
| rs1691271835 | 2:190969221 |
| rs1486511037 | 2:190969228 |
| rs944374411 | 2:190969229 |
| rs755752702 | 2:190969230 |
| rs879696627 | 2:190969231 |
| rs1345243082 | 2:190969233 |
| rs1244494846 | 2:190969236 |
| rs1574631502 | 2:190969237 |
| rs1313406203 | 2:190969248 |
| rs2124972499 | 2:190969252 |
| rs979463051 | 2:190969253 |
| rs1040006583 | 2:190969255 |
| rs746484042 | 2:190969256 |
| rs1574631529 | 2:190969259 |
| rs900296058 | 2:190969262 |
| rs937037547 | 2:190969263 |
| rs1299766797 | 2:190969264 |
| rs2124972555 | 2:190969270 |
| rs991085331 | 2:190969271 |
| rs1374679950 | 2:190969281 |
| rs1691278354 | 2:190969282 |
| rs995881201 | 2:190969285 |
| rs542136528 | 2:190969286 |
| rs1691279227 | 2:190969291 |
| rs2124972625 | 2:190969292 |
| rs917014338 | 2:190969293 |
| rs563782303 | 2:190969298 |
| rs1691279882 | 2:190969300 |
| rs1691280155 | 2:190969302 |
| rs1257637616 | 2:190969303 |
| rs11305 | 2:190969304 |
| rs1691281279 | 2:190969306 |
| rs1046888151 | 2:190969313 |
| rs887935841 | 2:190969315 |
| rs886973377 | 2:190969317 |
| rs2124972735 | 2:190969320 |
| rs1691282874 | 2:190969322 |
| rs1478394175 | 2:190969327 |
| rs1691283251 | 2:190969335 |
| rs1691283435 | 2:190969337 |
| rs1266013997 | 2:190969340 |
| rs1186197063 | 2:190969341 |
| rs941197964 | 2:190969344 |
| rs1691285148 | 2:190969348 |
| rs1005025126 | 2:190969356 |
| rs1200665369 | 2:190969366 |
| rs1691285729 | 2:190969367 |
| rs1484038511 | 2:190969369 |
| rs1249044418 | 2:190969372 |
| rs1691286504 | 2:190969378 |
| rs1691286691 | 2:190969381 |
| rs1038714220 | 2:190969383 |
| rs368576923 | 2:190969386 |
| rs996199996 | 2:190969393 |
| rs1354581153 | 2:190969394 |
| rs1691287603 | 2:190969396 |
| rs1691287819 | 2:190969398 |
| rs189030575 | 2:190969402 |
| rs1691288268 | 2:190969405 |
| rs1691288463 | 2:190969406 |
| rs1397943465 | 2:190969407 |
| rs1691288824 | 2:190969411 |
| rs981856342 | 2:190969412 |
| rs1034344469 | 2:190969413 |
| rs958816905 | 2:190969414 |
| rs1431755920 | 2:190969417 |
| rs1691289771 | 2:190969419 |
| rs1466227575 | 2:190969421 |
| rs1425931456 | 2:190969423 |
| rs990213925 | 2:190969429 |
| rs915105846 | 2:190969432 |
| rs2124973059 | 2:190969433 |
| rs886055374 | 2:190969443 |
| rs557207722 | 2:190969447 |
| rs1691291721 | 2:190969456 |
| rs1425145245 | 2:190969462 |
| rs1248007093 | 2:190969469 |
| rs1691292307 | 2:190969472 |
| rs1691292504 | 2:190969473 |
| rs1178663231 | 2:190969481 |
| rs1691292882 | 2:190969482 |
| rs1691293061 | 2:190969483 |
| rs1691293234 | 2:190969485 |
| rs1197681996 | 2:190969487 |
| rs1691293624 | 2:190969488 |
| rs1691293809 | 2:190969497 |
| rs1457799429 | 2:190969500 |
| rs2124973222 | 2:190969512 |
| rs1465557782 | 2:190969515 |
| rs1691294583 | 2:190969516 |
| rs1020366127 | 2:190969519 |
| rs1691294945 | 2:190969521 |
| rs967975098 | 2:190969528 |
| rs1691295501 | 2:190969536 |
| rs1334084021 | 2:190969539 |
| rs1329457174 | 2:190969540 |
| rs1691296285 | 2:190969542 |
| rs1000357522 | 2:190969543 |
| rs946544691 | 2:190969544 |
| rs1691296648 | 2:190969545 |
| rs1402477351 | 2:190969548 |
| rs977912956 | 2:190969549 |
| rs1691297234 | 2:190969550 |
| rs1306532833 | 2:190969555 |
| rs1033692335 | 2:190969560 |
| rs1691297580 | 2:190969562 |
| rs1447801311 | 2:190969565 |
| rs879538247 | 2:190969567 |
| rs1461320344 | 2:190969570 |
| rs1691298427 | 2:190969572 |
| rs1392122056 | 2:190969573 |
| rs1471371676 | 2:190969579 |
| rs2124973510 | 2:190969583 |
| rs1274440385 | 2:190969584 |
| rs1691299203 | 2:190969587 |
| rs1691299382 | 2:190969590 |
| rs2124973562 | 2:190969596 |
| rs1217310251 | 2:190969601 |
| rs1691300234 | 2:190969602 |
| rs550489659 | 2:190969609 |
| rs1691300895 | 2:190969617 |
| rs1363631252 | 2:190969622 |
| rs1159444997 | 2:190969624 |
| rs923863441 | 2:190969628 |
| rs1458685708 | 2:190969629 |
| rs1691302485 | 2:190969630 |
| rs1691302849 | 2:190969631 |
| rs944632360 | 2:190969633 |
| rs1239405513 | 2:190969634 |
| rs1191208553 | 2:190969635 |
| rs916880590 | 2:190969636 |
| rs780655672 | 2:190969637 |
| rs900160511 | 2:190969642 |
| rs570854463 | 2:190969643 |
| rs971028087 | 2:190969649 |
| rs982975198 | 2:190969650 |
| rs1259996266 | 2:190969655 |
| rs1241131501 | 2:190969666 |
| rs528626040 | 2:190969667 |
| rs887503321 | 2:190969668 |
| rs1005462562 | 2:190969670 |
| rs547101504 | 2:190969672 |
| rs1036496798 | 2:190969675 |
| rs907480414 | 2:190969677 |
| rs1003123973 | 2:190969678 |
| rs941218552 | 2:190969683 |
| rs1691307365 | 2:190969686 |
| rs1425906541 | 2:190969687 |
| rs1691307752 | 2:190969691 |
| rs1331803615 | 2:190969693 |
| rs1038185021 | 2:190969696 |
| rs1691308278 | 2:190969698 |
| rs1034372654 | 2:190969699 |
| rs1384687037 | 2:190969700 |
| rs562081130 | 2:190969701 |
| rs1691309203 | 2:190969704 |
| rs1691309372 | 2:190969708 |
| rs769403390 | 2:190969711 |
| rs2124973968 | 2:190969714 |
| rs1381808050 | 2:190969715 |
| rs777682355 | 2:190969716 |
| rs932524197 | 2:190969717 |
| rs749141524 | 2:190969719 |
| rs1477395561 | 2:190969723 |
| rs1246824597 | 2:190969724 |
| rs1691311123 | 2:190969725 |
| rs1691311299 | 2:190969727 |
| rs1691311467 | 2:190969728 |
| rs1200907242 | 2:190969733 |
| rs1021720997 | 2:190969736 |
| rs1691312134 | 2:190969739 |
| rs967454724 | 2:190969744 |
| rs1691312560 | 2:190969746 |
| rs2124974116 | 2:190969748 |
| rs1691312734 | 2:190969749 |
| rs41324145 | 2:190969750 |
| rs184180073 | 2:190969751 |
| rs1356062539 | 2:190969752 |
| rs569019701 | 2:190969754 |
| rs1691313660 | 2:190969758 |
| rs976014229 | 2:190969759 |
| rs41363648 | 2:190969761 |
| rs551312847 | 2:190969763 |
| rs2124974220 | 2:190969767 |
| rs1345840617 | 2:190969769 |
| rs886055375 | 2:190969771 |
| rs1691315410 | 2:190969781 |
| rs886055376 | 2:190969782 |
| rs1691315828 | 2:190969786 |
| rs1691316019 | 2:190969790 |
| rs2124974295 | 2:190969792 |
| rs1326360631 | 2:190969801 |
| rs1691316366 | 2:190969803 |
| rs1691316519 | 2:190969808 |
| rs1691316682 | 2:190969811 |
| rs1691316866 | 2:190969819 |
| rs1691317046 | 2:190969821 |
| rs566313103 | 2:190969824 |
| rs1691317390 | 2:190969825 |
| rs1691317560 | 2:190969826 |
| rs1299140983 | 2:190969827 |
| rs1048669825 | 2:190969829 |
| rs1374073561 | 2:190969836 |
| rs1691318285 | 2:190969837 |
| rs908972162 | 2:190969839 |
| rs1171326918 | 2:190969840 |
| rs1691318851 | 2:190969841 |
| rs1413622801 | 2:190969845 |
| rs1425238981 | 2:190969847 |
| rs188557905 | 2:190969848 |
| rs1214688680 | 2:190969852 |
| rs1474363657 | 2:190969856 |
| rs1691320042 | 2:190969857 |
| rs1691320331 | 2:190969867 |
| rs1288525683 | 2:190969873 |
| rs1422860396 | 2:190969875 |
| rs1691321254 | 2:190969876 |
| rs1194364922 | 2:190969878 |
| rs1691321794 | 2:190969879 |
| rs1691322060 | 2:190969880 |
| rs903239098 | 2:190969882 |
| rs1036975160 | 2:190969884 |
| rs1691322897 | 2:190969885 |
| rs1691323143 | 2:190969887 |
| rs907349445 | 2:190969889 |
| rs1691323996 | 2:190969892 |
| rs1033141232 | 2:190969894 |
| rs958955361 | 2:190969896 |
| rs1013154846 | 2:190969898 |
| rs1024508098 | 2:190969899 |
| rs1455797633 | 2:190969900 |
| rs1691325700 | 2:190969904 |
| rs1270055315 | 2:190969906 |
| rs1691326262 | 2:190969907 |
| rs2124974685 | 2:190969909 |
| rs1003564535 | 2:190969916 |
| rs1221044628 | 2:190969922 |
| rs180904823 | 2:190969926 |
| rs1691327680 | 2:190969928 |
| rs2124974748 | 2:190969929 |
| rs982454382 | 2:190969931 |
| rs1266031406 | 2:190969934 |
| rs1478415376 | 2:190969937 |
| rs908300239 | 2:190969943 |
| rs894706086 | 2:190969946 |
| rs1398973007 | 2:190969947 |
| rs2124974826 | 2:190969954 |
| rs1407303390 | 2:190969961 |
| rs1691330307 | 2:190969972 |
| rs1165708718 | 2:190969981 |
| rs772206364 | 2:190969982 |
| rs1691330829 | 2:190969984 |
| rs566925477 | 2:190969985 |
| rs537402240 | 2:190969988 |
| rs921134748 | 2:190969990 |
| rs1691331572 | 2:190969992 |
| rs1691331728 | 2:190969993 |
| rs1691331873 | 2:190970001 |
| rs1239449774 | 2:190970003 |
| rs1365987295 | 2:190970010 |
| rs1011592854 | 2:190970011 |
| rs1051032536 | 2:190970017 |
| rs1258870044 | 2:190970018 |
| rs1691332962 | 2:190970026 |
| rs1201056639 | 2:190970028 |
| rs186033487 | 2:190970033 |
| rs575893899 | 2:190970038 |
| rs45449693 | 2:190970042 |
| rs1691333987 | 2:190970043 |
| rs1691334144 | 2:190970046 |
| rs1381858784 | 2:190970055 |
| rs1691334718 | 2:190970057 |
| rs1285562768 | 2:190970058 |
| rs1398907439 | 2:190970059 |
| rs1691335665 | 2:190970060 |
| rs1691335845 | 2:190970062 |
| rs1031282966 | 2:190970064 |
| rs1392245161 | 2:190970068 |
| rs965920313 | 2:190970070 |
| rs1691336809 | 2:190970071 |
| rs2124975138 | 2:190970073 |
| rs975866030 | 2:190970075 |
| rs1163743171 | 2:190970078 |
| rs1691337951 | 2:190970083 |
| rs1415740340 | 2:190970091 |
| rs921851886 | 2:190970098 |
| rs760945547 | 2:190970101 |
| rs1691339163 | 2:190970105 |
| rs79073086 | 2:190970106 |
| rs146036682 | 2:190970112 |
| rs1180591844 | 2:190970114 |
| rs984455097 | 2:190970115 |
| rs1691340250 | 2:190970125 |
| rs2124975297 | 2:190970127 |
| rs190508584 | 2:190970129 |
| rs114360225 | 2:190970130 |
| rs1691341061 | 2:190970131 |
| rs2124975346 | 2:190970132 |
| rs1691341438 | 2:190970138 |
| rs1691341719 | 2:190970141 |
| rs1323876080 | 2:190970144 |
| rs907433798 | 2:190970146 |
| rs2124975396 | 2:190970147 |
| rs1691342654 | 2:190970149 |
| rs1237438235 | 2:190970150 |
| rs1691343112 | 2:190970152 |
| rs1574632305 | 2:190970153 |
| rs1004290548 | 2:190970155 |
| rs41476445 | 2:190970168 |
| rs1448379540 | 2:190970170 |
| rs1036098684 | 2:190970171 |
| rs1292838083 | 2:190970172 |
| rs1574632336 | 2:190970178 |
| rs1437185939 | 2:190970179 |
| rs1351391769 | 2:190970182 |
| rs1156626637 | 2:190970184 |
| rs1691346688 | 2:190970185 |
| rs1448097711 | 2:190970188 |
| rs1381185396 | 2:190970193 |
| rs1160732426 | 2:190970198 |
| rs754155168 | 2:190970200 |
| rs1452625062 | 2:190970201 |
| rs1187379026 | 2:190970205 |
| rs1691348782 | 2:190970210 |
| rs938880825 | 2:190970221 |
| rs1691349404 | 2:190970224 |
| rs1192270071 | 2:190970234 |
| rs1691350280 | 2:190970237 |
| rs200344731 | 2:190970242 |
| rs1691351235 | 2:190970243 |
| rs1691351529 | 2:190970244 |
| rs1691351822 | 2:190970246 |
| rs1240659511 | 2:190970250 |
| rs1691352451 | 2:190970255 |
| rs1055938745 | 2:190970260 |
| rs1473792625 | 2:190970261 |
| rs1691353518 | 2:190970262 |
| rs529316850 | 2:190970263 |
| rs1352469364 | 2:190970266 |
| rs1286254077 | 2:190970274 |
| rs139958571 | 2:190970275 |
| rs562899714 | 2:190970276 |
| rs1691354815 | 2:190970284 |
| rs1309927096 | 2:190970286 |
| rs1300061520 | 2:190970288 |
| rs1406977266 | 2:190970289 |
| rs1156677557 | 2:190970290 |
| rs182394503 | 2:190970297 |
| rs1296073393 | 2:190970298 |
| rs1691356161 | 2:190970303 |
| rs2124975782 | 2:190970305 |
| rs1401221651 | 2:190970307 |
| rs1691356580 | 2:190970314 |
| rs1691357316 | 2:190970327 |
| rs1691357473 | 2:190970328 |
| rs1691357627 | 2:190970333 |
| rs1300275650 | 2:190970337 |
| rs186032149 | 2:190970339 |
| rs1691358240 | 2:190970340 |
| rs886055377 | 2:190970341 |
| rs903172029 | 2:190970342 |
| rs1691358827 | 2:190970343 |
| rs1691359008 | 2:190970345 |
| rs566349352 | 2:190970346 |
| rs977832294 | 2:190970348 |
| rs1193632643 | 2:190970364 |
| rs777014385 | 2:190970367 |
| rs1691360023 | 2:190970368 |
| rs1691360160 | 2:190970374 |
| rs1691360323 | 2:190970384 |
| rs1691360490 | 2:190970386 |
| rs999235618 | 2:190970390 |
| rs935996678 | 2:190970392 |
| rs1691361095 | 2:190970395 |
| rs1315095292 | 2:190970397 |
| rs190542524 | 2:190970399 |
| rs1233377807 | 2:190970405 |
| rs1691361941 | 2:190970416 |
| rs1574632511 | 2:190970418 |
| rs1691362488 | 2:190970419 |
| rs1313832882 | 2:190970420 |
| rs1311927895 | 2:190970421 |
| rs762197567 | 2:190970422 |
| rs1200412744 | 2:190970423 |
| rs997310104 | 2:190970427 |
| rs1574632532 | 2:190970429 |
| rs1691364575 | 2:190970432 |
| rs1028773305 | 2:190970433 |
| rs1691365092 | 2:190970434 |
| rs765667420 | 2:190970435 |
| rs548677582 | 2:190970436 |
| rs1395511430 | 2:190970437 |
| rs182725919 | 2:190970440 |
| rs1401808260 | 2:190970442 |
| rs1303456711 | 2:190970446 |
| rs1691367171 | 2:190970450 |
| rs1691367688 | 2:190970452 |
| rs984653102 | 2:190970457 |
| rs1691368530 | 2:190970465 |
| rs1204464083 | 2:190970466 |
| rs1398246936 | 2:190970467 |
| rs1691369417 | 2:190970479 |
| rs1237627664 | 2:190970488 |
| rs1016371601 | 2:190970489 |
| rs907458830 | 2:190970491 |
| rs961749006 | 2:190970492 |
| rs1574632596 | 2:190970503 |
| rs1691371155 | 2:190970510 |
| rs1004504808 | 2:190970511 |
| rs1691371971 | 2:190970518 |
| rs537942049 | 2:190970522 |
| rs750941607 | 2:190970523 |
| rs1365351649 | 2:190970529 |
| rs1470359868 | 2:190970531 |
| rs972115850 | 2:190970538 |
| rs1691374598 | 2:190970539 |
| rs1421648128 | 2:190970541 |
| rs2124976473 | 2:190970547 |
| rs41481847 | 2:190970549 |
| rs1200715284 | 2:190970560 |
| rs768945585 | 2:190970562 |
| rs1691376160 | 2:190970563 |
| rs571207686 | 2:190970566 |
| rs1197872838 | 2:190970573 |
| rs1691377027 | 2:190970578 |
| rs1240953570 | 2:190970581 |
| rs938403650 | 2:190970582 |
| rs1259708984 | 2:190970584 |
| rs1691378212 | 2:190970593 |
| rs991696488 | 2:190970601 |
| rs1691378920 | 2:190970603 |
| rs1313664856 | 2:190970610 |
| rs1691379305 | 2:190970612 |
| rs1028090653 | 2:190970618 |
| rs1330776493 | 2:190970623 |
| rs1691380171 | 2:190970626 |
| rs1288692855 | 2:190970630 |
| rs1444165759 | 2:190970632 |
| rs1669547443 | 2:190970635 |
| rs1691381102 | 2:190970638 |
| rs1403123349 | 2:190970646 |
| rs1691381679 | 2:190970647 |
| rs1691382007 | 2:190970648 |
| rs916064299 | 2:190970650 |
| rs1691382612 | 2:190970651 |
| rs548086115 | 2:190970652 |
| rs749380631 | 2:190970655 |
| rs1161774047 | 2:190970659 |
| rs953752983 | 2:190970661 |
| rs768700585 | 2:190970664 |
| rs1218764714 | 2:190970669 |
| rs1369475742 | 2:190970670 |
| rs774654212 | 2:190970674 |
| rs762042700 | 2:190970675 |
| rs1255205369 | 2:190970679 |
| rs1691386310 | 2:190970683 |
| rs1177503647 | 2:190970684 |
| rs947648115 | 2:190970686 |
| rs767892470 | 2:190970694 |
| rs773624561 | 2:190970696 |
| rs983566520 | 2:190975544 |
| rs1574637199 | 2:190975546 |
| rs1412164181 | 2:190975561 |
| rs1691840652 | 2:190975568 |
| rs1691841178 | 2:190975571 |
| rs1335503966 | 2:190975573 |
| rs1412318644 | 2:190975574 |
| rs1389517862 | 2:190975578 |
| rs1440676684 | 2:190975580 |
| rs2124995604 | 2:190975589 |
| rs1163018930 | 2:190975595 |
| rs2124995622 | 2:190975597 |
| rs978510232 | 2:190975599 |
| rs1418118567 | 2:190975601 |
| rs1691843144 | 2:190975602 |
| rs1691843405 | 2:190975608 |
| rs925679024 | 2:190975610 |
| rs1481752857 | 2:190975615 |
| rs1574637239 | 2:190975617 |
| rs1574637245 | 2:190975618 |
| rs1691844903 | 2:190975622 |
| rs1205082591 | 2:190975623 |
| rs1574637259 | 2:190975624 |
| rs1574637266 | 2:190975625 |
| rs1287699892 | 2:190975626 |
| rs937196233 | 2:190975632 |
| rs1691847541 | 2:190975635 |
| rs879583102 | 2:190975641 |
| rs898814962 | 2:190975642 |
| rs77910835 | 2:190975643 |
| rs1280181707 | 2:190975645 |
| rs1574637316 | 2:190975646 |
| rs112071828 | 2:190975648 |
| rs1574637322 | 2:190975649 |
| rs1047419091 | 2:190975655 |
| rs1691850708 | 2:190975657 |
| rs1691850953 | 2:190975659 |
| rs1691851227 | 2:190975660 |
| rs1691851475 | 2:190975668 |
| rs566136866 | 2:190975672 |
| rs940656467 | 2:190975673 |
| rs1691852388 | 2:190975674 |
| rs1691852636 | 2:190975675 |
| rs116554639 | 2:190975678 |
| rs1691853275 | 2:190975682 |
| rs1471423016 | 2:190975683 |
| rs554936429 | 2:190975684 |
| rs1691854401 | 2:190975692 |
| rs1691854828 | 2:190975696 |
| rs2124996125 | 2:190975697 |
| rs1410817426 | 2:190975698 |
| rs1691855410 | 2:190975700 |
| rs899224935 | 2:190975702 |
| rs575057281 | 2:190975703 |
| rs961200796 | 2:190975705 |
| rs45623541 | 2:190975708 |
| rs1029197409 | 2:190975715 |
| rs1691857143 | 2:190975718 |
| rs1274851520 | 2:190975723 |
| rs927902093 | 2:190975729 |
| rs1691858237 | 2:190975730 |
| rs1691858500 | 2:190975732 |
| rs1179362530 | 2:190975734 |
| rs1691859032 | 2:190975735 |
| rs1691859260 | 2:190975737 |
| rs1691859484 | 2:190975739 |
| rs890617468 | 2:190975745 |
| rs1691860238 | 2:190975747 |
| rs1328729536 | 2:190975752 |
| rs2124996379 | 2:190975754 |
| rs1691860946 | 2:190975755 |
| rs201391478 | 2:190975760 |
| rs753796178 | 2:190975764 |
| rs11549698 | 2:190975766 |
| rs1691862717 | 2:190975769 |
| rs779046006 | 2:190975770 |
| rs2124996458 | 2:190975771 |
| rs369953295 | 2:190975772 |
| rs1284802601 | 2:190975773 |
| rs748181005 | 2:190975776 |
| rs1211217813 | 2:190975786 |
| rs1691864605 | 2:190975791 |
| rs372657991 | 2:190975792 |
| rs1691865158 | 2:190975793 |
| rs777991780 | 2:190975796 |
| rs1442310621 | 2:190975799 |
| rs2124996589 | 2:190975805 |
| rs1691866152 | 2:190975807 |

Table S3. Regulome DB score for the 5 prime UTR

| SNP ID |  | rank |
| --- | --- | --- |
| rs1467622177 | 0.60906 | 4 |
| rs1559026214 | 0.60906 | 4 |
| rs960205570 | 0.60906 | 4 |
| rs567434849 | 0.60906 | 4 |
| rs531511204 | 0.58955 | 5 |
| rs963769760 | 0.58955 | 5 |
| rs1191503043 | 0.58955 | 5 |
| rs1405565781 | 0.58955 | 5 |
| rs1372817347 | 0.58955 | 5 |
| rs765693257 | 0.8575 | 5 |
| rs1459026520 | 0.58955 | 5 |
| rs549949329 | 0.58955 | 5 |
| rs1405849643 | 0.58955 | 5 |
| rs571354699 | 0.58955 | 5 |
| rs1238278593 | 0.58955 | 5 |
| rs1180393338 | 0.58955 | 5 |
| rs909295245 | 0.58955 | 5 |
| rs750862823 | 0.58955 | 5 |
| rs772559222 | 0.58955 | 5 |
| rs1490002122 | 0.58955 | 5 |
| rs902278776 | 0.70497 | 4 |
| rs1408491477 | 0.70497 | 4 |
| rs372299811 | 0.60906 | 4 |
| rs1220675431 | 0.70497 | 4 |
| rs999306857 | 0.70497 | 4 |
| rs189188567 | 0.70497 | 4 |
| rs1279334374 | 0.70497 | 4 |
| rs1490866036 | 0.60906 | 4 |
| rs1240818117 | 0.60906 | 4 |
| rs1215740346 | 0.60906 | 4 |
| rs11549697 | 0.60906 | 4 |
| rs1286840473 | 0.60906 | 4 |
| rs1053528132 | 0.60906 | 4 |
| rs1201881287 | 0.6749 | 2b |
| rs938419162 | 0.7614 | 2b |
| rs1559028129 | 0.60906 | 4 |
| rs1282245963 | 0.60906 | 4 |
| rs1223531891 | 0.60906 | 4 |
| rs1369144548 | 0.60906 | 4 |
| rs545742697 | 0.60906 | 4 |
| rs1429996774 | 0.60906 | 4 |
| rs1345553432 | 0.945 | 2b |
| rs1320123706 | 0.61749 | 2b |
| rs1483073172 | 0.63331 | 2b |
| rs1056003060 | 0.70497 | 4 |
| rs1481434204 | 0.70497 | 4 |
| rs1198263292 | 0.70497 | 4 |
| rs118149197 | 0.60906 | 4 |
| rs144940940 | 0.70497 | 4 |
| rs947541007 | 0.70497 | 4 |
| rs1317848379 | 0.60906 | 4 |
| rs993374058 | 0.60906 | 4 |
| rs556645752 | 0.60906 | 4 |
| rs994096848 | 0.60906 | 4 |
| rs1035535139 | 0.60906 | 4 |
| rs55724546 | 0.60906 | 4 |
| rs952804507 | 0.60906 | 4 |
| rs1325660325 | 0.60906 | 4 |
| rs45470392 | 1 | 2a |
| rs1413522785 | 1 | 2a |
| rs1299735457 | 0.71481 | 2a |
| rs986046425 | 0.70787 | 2a |
| rs1490047123 | 0.70497 | 4 |
| rs1269450206 | 0.60906 | 4 |
| rs1559028449 | 0.60906 | 4 |
| rs1211595473 | 0.60906 | 4 |
| rs147950001 | 0.60906 | 4 |
| rs965520149 | 0.60906 | 4 |
| rs1240024048 | 0.60906 | 4 |
| rs1331407038 | 0.60906 | 4 |
| rs564075337 | 0.60906 | 4 |
| rs1263141636 | 0.60906 | 4 |
| rs968480409 | 0.60906 | 4 |
| rs978953371 | 0.60906 | 4 |
| rs934985471 | 0.60906 | 4 |
| rs989136304 | 0.60906 | 4 |
| rs914987046 | 0.60906 | 4 |
| rs1384678089 | 0.60906 | 4 |
| rs528057545 | 0.60906 | 4 |
| rs1210087528 | 0.60906 | 4 |
| rs947858016 | 0.60906 | 4 |
| rs568025055 | 0.60906 | 4 |
| rs560084350 | 0.60906 | 4 |
| rs371724653 | 0.60906 | 4 |
| rs1466106799 | 0.60906 | 4 |
| rs1445643161 | 0.60906 | 4 |
| rs1026959454 | 0.60906 | 4 |
| rs1276743117 | 0.60906 | 4 |
| rs977004227 | 0.70497 | 4 |
| rs769850162 | 0.70497 | 4 |
| rs1181864140 | 0.70497 | 4 |
| rs922842738 | 0.74401 | 4 |
| rs1007000950 | 0.74401 | 4 |
| rs530356200 | 0.74401 | 4 |
| rs548761843 | 0.60906 | 4 |
| rs888550057 | 0.60906 | 4 |
| rs1460039243 | 0.60906 | 4 |
| rs886055382 | 0.74401 | 4 |
| rs570168205 | 0.84 | 2a |
| rs1037537872 | 1 | 2a |
| rs531009254 | 0.9 | 2a |
| rs969295139 | 0.3145 | 2a |

Table S4. Regulome DB score for the 3 prime UTR

| SNP ID |  | Rank |
| --- | --- | --- |
| rs537942049 | 0.60906 | 4 |
| rs750941607 | 0.60906 | 4 |
| rs1365351649 | 0.60906 | 4 |
| rs1470359868 | 0.60906 | 4 |
| rs972115850 | 0.60906 | 4 |
| rs1421648128 | 0.60906 | 4 |
| rs41481847 | 0.60906 | 4 |
| rs1200715284 | 0.60906 | 4 |
| rs768945585 | 0.60906 | 4 |
| rs571207686 | 0.71481 | 2a |
| rs1197872838 | 0.17875 | 2a |
| rs1240953570 | 0.60906 | 4 |
| rs938403650 | 0.60906 | 4 |
| rs1259708984 | 0.60906 | 4 |
| rs991696488 | 0.65734 | 2b |
| rs1313664856 | 0.59638 | 2b |
| rs1028090653 | 1 | 2b |
| rs1330776493 | 0.66759 | 2b |
| rs1288692855 | 0.42482 | 2b |
| rs1444165759 | 0.60906 | 4 |
| rs1403123349 | 0.60906 | 4 |
| rs916064299 | 0.60906 | 4 |
| rs548086115 | 0.60906 | 4 |
| rs749380631 | 0.60906 | 4 |
| rs1161774047 | 0.60906 | 4 |
| rs953752983 | 0.60906 | 4 |
| rs768700585 | 0.85505 | 3a |
| rs1218764714 | 0.60567 | 3a |
| rs1369475742 | 0.60906 | 4 |
| rs774654212 | 0.60906 | 4 |
| rs762042700 | 0.60906 | 4 |
| rs1255205369 | 0.60906 | 4 |
| rs1177503647 | 0.60906 | 4 |
| rs947648115 | 0.60906 | 4 |
| rs767892470 | 0.60906 | 4 |
| rs773624561 | 0.60906 | 4 |
| rs983566520 | 0.58955 | 5 |
| rs1412164181 | 0.62525 | 5 |
| rs1335503966 | 0.39333 | 5 |
| rs1412318644 | 0.74 | 5 |
| rs1389517862 | 0.51855 | 5 |
| rs1440676684 | 0.7075 | 5 |
| rs1163018930 | 0.58955 | 5 |
| rs978510232 | 0.58955 | 5 |
| rs1418118567 | 0.58955 | 5 |
| rs925679024 | 0.58955 | 5 |
| rs1481752857 | 0.58955 | 5 |
| rs1205082591 | 0.08 | 5 |
| rs1287699892 | 0.54974 | 5 |
| rs937196233 | 0.85633 | 5 |
| rs879583102 | 0.88556 | 5 |
| rs898814962 | 0.48781 | 5 |
| rs77910835 | 0.48781 | 5 |
| rs1280181707 | 1 | 5 |
| rs112071828 | 0.64776 | 2b |
| rs1047419091 | 0.89367 | 2b |
| rs566136866 | 0.60906 | 4 |
| rs940656467 | 0.60906 | 4 |
| rs116554639 | 0.60906 | 4 |
| rs1471423016 | 0.60906 | 4 |
| rs554936429 | 0.60906 | 4 |
| rs1410817426 | 0.60906 | 4 |
| rs899224935 | 0.60906 | 4 |
| rs575057281 | 0.60906 | 4 |
| rs961200796 | 0.60906 | 4 |
| rs45623541 | 0.406 | 2b |
| rs1029197409 | 0.73655 | 2b |
| rs1274851520 | 0.70377 | 2b |
| rs927902093 | 0.60906 | 4 |
| rs1179362530 | 0.60906 | 4 |
| rs890617468 | 1 | 2b |
| rs1328729536 | 0.56896 | 2b |
| rs201391478 | 0.60906 | 4 |
| rs753796178 | 0.60906 | 4 |
| rs11549698 | 0.60906 | 4 |
| rs779046006 | 0.76041 | 2b |
| rs369953295 | 0.71939 | 2b |
| rs1284802601 | 0.73843 | 2b |
| rs748181005 | 0.80226 | 2b |
| rs1211217813 | 0.90505 | 3a |
| rs372657991 | 0.60906 | 4 |
| rs777991780 | 0.60906 | 4 |
| rs1442310621 | 0.60906 | 4 |
| rs188557905 | 0.60906 | 4 |
| rs1214688680 | 0.60906 | 4 |
| rs1474363657 | 0.60906 | 4 |
| rs1288525683 | 0.60906 | 4 |
| rs1422860396 | 0.60906 | 4 |
| rs1194364922 | 0.60906 | 4 |
| rs903239098 | 0.60906 | 4 |
| rs1036975160 | 0.60906 | 4 |
| rs907349445 | 0.60906 | 4 |
| rs1033141232 | 0.60906 | 4 |
| rs958955361 | 0.60906 | 4 |
| rs1013154846 | 0.60906 | 4 |
| rs1024508098 | 0.8288 | 2b |
| rs1455797633 | 0.7516 | 2b |
| rs1270055315 | 0.57802 | 2b |
| rs1003564535 | 0.60906 | 4 |
| rs1221044628 | 0.60906 | 4 |
| rs180904823 | 0.60906 | 4 |
| rs982454382 | 0.60906 | 4 |
| rs1266031406 | 0.60906 | 4 |
| rs1478415376 | 0.77783 | 2b |
| rs908300239 | 0.81166 | 2b |
| rs894706086 | 0.51348 | 2b |
| rs1398973007 | 0.73843 | 2b |
| rs1407303390 | 0.60906 | 4 |
| rs1165708718 | 0.60906 | 4 |
| rs772206364 | 0.60906 | 4 |
| rs566925477 | 0.60906 | 4 |
| rs537402240 | 0.60906 | 4 |
| rs921134748 | 0.60906 | 4 |
| rs1239449774 | 0.60906 | 4 |
| rs1365987295 | 0.60906 | 4 |
| rs1011592854 | 0.60906 | 4 |
| rs1051032536 | 0.60906 | 4 |
| rs1258870044 | 0.60906 | 4 |
| rs1201056639 | 0.73502 | 3a |
| rs186033487 | 0.51977 | 3a |
| rs575893899 | 0.60906 | 4 |
| rs45449693 | 0.60906 | 4 |
| rs1381858784 | 0.60906 | 4 |
| rs1285562768 | 0.60906 | 4 |
| rs1398907439 | 0.60906 | 4 |
| rs1031282966 | 0.84289 | 2b |
| rs1392245161 | 0.82541 | 2b |
| rs965920313 | 0.41199 | 2b |
| rs975866030 | 0.50526 | 2b |
| rs1163743171 | 0.6561 | 3a |
| rs1415740340 | 0.89367 | 2b |
| rs921851886 | 0.7614 | 2b |
| rs760945547 | 0.40272 | 2b |
| rs79073086 | 0.60906 | 4 |
| rs146036682 | 0.60906 | 4 |
| rs1180591844 | 0.49614 | 2b |
| rs984455097 | 0.58489 | 2b |
| rs190508584 | 0.60906 | 4 |
| rs114360225 | 0.70377 | 2b |
| rs1323876080 | 0.79615 | 2b |
| rs907433798 | 0.34744 | 2b |
| rs1237438235 | 0.71276 | 2b |
| rs1004290548 | 0.64249 | 2b |
| rs41476445 | 0.60906 | 4 |
| rs1448379540 | 0.60906 | 4 |
| rs1036098684 | 0.60906 | 4 |
| rs1292838083 | 0.60906 | 4 |
| rs1437185939 | 0.60906 | 4 |
| rs1351391769 | 0.60906 | 4 |
| rs1156626637 | 0.60906 | 4 |
| rs1448097711 | 0.60906 | 4 |
| rs1381185396 | 0.60906 | 4 |
| rs1160732426 | 0.64591 | 2b |
| rs754155168 | 0.79882 | 2b |
| rs1452625062 | 0.7614 | 2b |
| rs1187379026 | 0.2664 | 2b |
| rs938880825 | 0.60906 | 4 |
| rs1192270071 | 0.60906 | 4 |
| rs200344731 | 0.60906 | 4 |
| rs1240659511 | 0.60906 | 4 |
| rs1055938745 | 0.60906 | 4 |
| rs1473792625 | 0.60906 | 4 |
| rs529316850 | 0.60906 | 4 |
| rs1352469364 | 0.60906 | 4 |
| rs1286254077 | 0.61652 | 2b |
| rs139958571 | 0.65734 | 2b |
| rs562899714 | 0.86817 | 2b |
| rs1309927096 | 0.60906 | 4 |
| rs1300061520 | 0.60906 | 4 |
| rs1406977266 | 0.60906 | 4 |
| rs1156677557 | 0.60906 | 4 |
| rs182394503 | 0.60906 | 4 |
| rs1296073393 | 0.60906 | 4 |
| rs1401221651 | 0.60906 | 4 |
| rs1300275650 | 0.60906 | 4 |
| rs186032149 | 0.60906 | 4 |
| rs886055377 | 0.60906 | 4 |
| rs903172029 | 0.60906 | 4 |
| rs566349352 | 0.70497 | 4 |
| rs977832294 | 0.70497 | 4 |
| rs1193632643 | 0.60906 | 4 |
| rs777014385 | 0.60906 | 4 |
| rs999235618 | 0.60906 | 4 |
| rs935996678 | 0.60906 | 4 |
| rs1315095292 | 0.60906 | 4 |
| rs190542524 | 0.60906 | 4 |
| rs1233377807 | 0.70497 | 4 |
| rs1313832882 | 0.60906 | 4 |
| rs1311927895 | 0.60906 | 4 |
| rs762197567 | 0.60906 | 4 |
| rs1200412744 | 0.60906 | 4 |
| rs997310104 | 0.60906 | 4 |
| rs1028773305 | 0.60906 | 4 |
| rs765667420 | 0.60906 | 4 |
| rs548677582 | 0.60906 | 4 |
| rs1395511430 | 0.60906 | 4 |
| rs182725919 | 0.60906 | 4 |
| rs1401808260 | 0.60906 | 4 |
| rs1303456711 | 0.60906 | 4 |
| rs984653102 | 0.60906 | 4 |
| rs1204464083 | 0.60906 | 4 |
| rs1398246936 | 0.60906 | 4 |
| rs1237627664 | 0.70497 | 4 |
| rs1016371601 | 0.70497 | 4 |
| rs907458830 | 0.70497 | 4 |
| rs961749006 | 0.70497 | 4 |
| rs1004504808 | 0.60906 | 4 |
| rs979463051 | 0.60906 | 4 |
| rs1040006583 | 0.60906 | 4 |
| rs746484042 | 0.60906 | 4 |
| rs900296058 | 0.60906 | 4 |
| rs937037547 | 0.60906 | 4 |
| rs1299766797 | 0.60906 | 4 |
| rs991085331 | 0.83617 | 2b |
| rs1374679950 | 0.63331 | 2b |
| rs995881201 | 0.68277 | 2b |
| rs542136528 | 0.67002 | 2b |
| rs917014338 | 0.60906 | 4 |
| rs563782303 | 0.60906 | 4 |
| rs1257637616 | 0.60906 | 4 |
| rs11305 | 0.60906 | 4 |
| rs1046888151 | 0.13454 | 5 |
| rs887935841 | 0.13454 | 5 |
| rs886973377 | 0.13454 | 5 |
| rs1478394175 | 0.13454 | 5 |
| rs1266013997 | 0.13454 | 5 |
| rs1186197063 | 0.13454 | 5 |
| rs941197964 | 0.13454 | 5 |
| rs1005025126 | 0.13454 | 5 |
| rs1200665369 | 0.60906 | 4 |
| rs1484038511 | 0.60906 | 4 |
| rs1249044418 | 0.60906 | 4 |
| rs1038714220 | 0.60906 | 4 |
| rs368576923 | 0.34744 | 2b |
| rs996199996 | 0.80975 | 2b |
| rs1354581153 | 0.82852 | 2b |
| rs189030575 | 0.60906 | 4 |
| rs1397943465 | 0.60906 | 4 |
| rs981856342 | 0.60906 | 4 |
| rs1034344469 | 0.60906 | 4 |
| rs958816905 | 0.60906 | 4 |
| rs1431755920 | 0.60906 | 4 |
| rs1466227575 | 0.85505 | 3a |
| rs1425931456 | 0.85505 | 3a |
| rs990213925 | 0.60906 | 4 |
| rs915105846 | 0.60906 | 4 |
| rs886055374 | 0.60906 | 4 |
| rs557207722 | 0.60906 | 4 |
| rs1425145245 | 0.60906 | 4 |
| rs1248007093 | 0.60906 | 4 |
| rs1178663231 | 0.60906 | 4 |
| rs1197681996 | 0.60906 | 4 |
| rs1457799429 | 0.60906 | 4 |
| rs1465557782 | 0.091 | 2b |
| rs1020366127 | 0.85505 | 3a |
| rs967975098 | 0.60906 | 4 |
| rs1334084021 | 0.60906 | 4 |
| rs1329457174 | 0.60906 | 4 |
| rs1000357522 | 0.60906 | 4 |
| rs946544691 | 0.60906 | 4 |
| rs1402477351 | 0.60906 | 4 |
| rs977912956 | 0.60906 | 4 |
| rs1306532833 | 0.60906 | 4 |
| rs1033692335 | 0.67017 | 2b |
| rs1447801311 | 0.71276 | 2b |
| rs879538247 | 0.8288 | 2b |
| rs1461320344 | 0.60906 | 4 |
| rs1392122056 | 0.86056 | 3a |
| rs1471371676 | 0.85505 | 3a |
| rs1274440385 | 0.60906 | 4 |
| rs1217310251 | 0.60906 | 4 |
| rs550489659 | 0.60906 | 4 |
| rs1363631252 | 0.60906 | 4 |
| rs1159444997 | 0.60906 | 4 |
| rs923863441 | 0.60906 | 4 |
| rs1458685708 | 0.60906 | 4 |
| rs944632360 | 0.60906 | 4 |
| rs1239405513 | 0.60906 | 4 |
| rs1191208553 | 0.60906 | 4 |
| rs916880590 | 0.60906 | 4 |
| rs780655672 | 0.60906 | 4 |
| rs900160511 | 0.86056 | 3a |
| rs570854463 | 0.38672 | 3a |
| rs971028087 | 0.60906 | 4 |
| rs982975198 | 0.60906 | 4 |
| rs1259996266 | 0.60906 | 4 |
| rs1241131501 | 0.60906 | 4 |
| rs528626040 | 0.60906 | 4 |
| rs887503321 | 0.60906 | 4 |
| rs1005462562 | 0.60906 | 4 |
| rs547101504 | 0.60906 | 4 |
| rs1036496798 | 0.60906 | 4 |
| rs907480414 | 0.60906 | 4 |
| rs1003123973 | 0.60906 | 4 |
| rs941218552 | 0.60906 | 4 |
| rs1425906541 | 0.60906 | 4 |
| rs1331803615 | 0.60906 | 4 |
| rs1038185021 | 0.60906 | 4 |
| rs1034372654 | 0.60906 | 4 |
| rs1384687037 | 0.60906 | 4 |
| rs562081130 | 0.60906 | 4 |
| rs769403390 | 0.60906 | 4 |
| rs1381808050 | 0.60906 | 4 |
| rs777682355 | 0.60906 | 4 |
| rs932524197 | 0.60906 | 4 |
| rs749141524 | 0.60906 | 4 |
| rs1477395561 | 0.60906 | 4 |
| rs1246824597 | 0.60906 | 4 |
| rs1200907242 | 0.60906 | 4 |
| rs1021720997 | 0.60906 | 4 |
| rs967454724 | 0.60906 | 4 |
| rs41324145 | 0.60906 | 4 |
| rs184180073 | 0.60906 | 4 |
| rs1356062539 | 0.60906 | 4 |
| rs569019701 | 0.60906 | 4 |
| rs976014229 | 0.60906 | 4 |
| rs41363648 | 0.60906 | 4 |
| rs551312847 | 0.60906 | 4 |
| rs1345840617 | 0.60906 | 4 |
| rs886055375 | 0.60906 | 4 |
| rs886055376 | 0.60906 | 4 |
| rs1326360631 | 0.60906 | 4 |
| rs566313103 | 0.60906 | 4 |
| rs1299140983 | 0.60906 | 4 |
| rs1048669825 | 0.69102 | 3a |
| rs1374073561 | 0.60906 | 4 |
| rs908972162 | 0.60906 | 4 |
| rs1171326918 | 0.60906 | 4 |
| rs1413622801 | 0.60906 | 4 |
| rs1425238981 | 0.60906 | 4 |
| rs1392985321 | 0.58955 | 5 |
| rs192670482 | 0.58955 | 5 |
| rs200913982 | 0.04578 | 5 |
| rs1222328384 | 0.04578 | 5 |
| rs1044848643 | 0.75333 | 5 |
| rs369150147 | 0.58955 | 5 |
| rs1012769799 | 0.60906 | 4 |
| rs1427402088 | 0.60906 | 4 |
| rs536205722 | 0.60906 | 4 |
| rs1480770453 | 0.8288 | 2b |
| rs556104368 | 0.71571 | 2b |
| rs1023179868 | 0.60906 | 4 |
| rs1272104884 | 0.60906 | 4 |
| rs143345038 | 0.60906 | 4 |
| rs1000105374 | 0.60906 | 4 |
| rs1272302640 | 0.60906 | 4 |
| rs1168 | 0.55436 | 1f |
| rs1323892661 | 0.60906 | 4 |
| rs556506301 | 0.99917 | 3a |
| rs1361164169 | 0.60906 | 4 |
| rs955670740 | 0.60906 | 4 |
| rs965812479 | 0.82541 | 2b |
| rs976525545 | 0.57802 | 2b |
| rs747337724 | 0.60906 | 4 |
| rs146824155 | 0.60906 | 4 |
| rs924065088 | 0.60906 | 4 |
| rs545184536 | 0.60906 | 4 |
| rs1243426905 | 0.58955 | 5 |
| rs1379489171 | 0.58955 | 5 |
| rs1242907769 | 0.58955 | 5 |
| rs953889876 | 0.58955 | 5 |
| rs1257447210 | 0.58955 | 5 |
| rs868463796 | 0.58955 | 5 |
| rs1201769718 | 0.58955 | 5 |
| rs1306448620 | 0.58955 | 5 |
| rs554209165 | 0.04578 | 5 |
| rs915465134 | 0.04578 | 5 |
| rs1380575153 | 0.04578 | 5 |
| rs1416099500 | 0.58955 | 5 |
| rs140488842 | 0.58955 | 5 |
| rs941657708 | 0.58955 | 5 |
| rs972941504 | 0.58955 | 5 |
| rs918616783 | 0.58955 | 5 |
| rs1381552086 | 0.58955 | 5 |
| rs1296698533 | 0.992 | 5 |
| rs1398955907 | 0.22625 | 5 |
| rs563090709 | 0.58955 | 5 |
| rs1343050347 | 0.58955 | 5 |
| rs1056845485 | 0.58955 | 5 |
| rs1216429025 | 0.58955 | 5 |
| rs895232493 | 0.58955 | 5 |
| rs1446849892 | 0.58955 | 5 |
| rs879641888 | 0.58955 | 5 |
| rs1175127546 | 0.58955 | 5 |
| rs1036156937 | 0.58955 | 5 |
| rs769148496 | 0.58955 | 5 |
| rs149993422 | 0.60906 | 4 |
| rs867842021 | 0.60906 | 4 |
| rs560939666 | 0.7614 | 2b |
| rs1027603252 | 0.84289 | 2b |
| rs1323288887 | 0.33881 | 3a |
| rs1305062112 | 0.60906 | 4 |
| rs575978803 | 0.60906 | 4 |
| rs904352199 | 0.60906 | 4 |
| rs1000137833 | 0.40272 | 2b |
| rs1403234504 | 0.99513 | 2b |
| rs1365452819 | 0.51305 | 2b |
| rs1284671779 | 0.60906 | 4 |
| rs543681256 | 0.60906 | 4 |
| rs564968175 | 0.60906 | 4 |
| rs1018656137 | 0.76502 | 2b |
| rs1488434355 | 0.60906 | 4 |
| rs779962342 | 0.74722 | 5 |
| rs1183645904 | 0.54036 | 5 |
| rs1255871629 | 0.99471 | 5 |
| rs1029709527 | 0.58955 | 5 |
| rs1489455527 | 0.58955 | 5 |
| rs185595627 | 0.58955 | 5 |
| rs190448557 | 0.58955 | 5 |
| rs1017128964 | 0.58955 | 5 |
| rs1238114169 | 0.58955 | 5 |
| rs374737590 | 0.58955 | 5 |
| rs1188858771 | 0.58955 | 5 |
| rs962840379 | 0.7075 | 5 |
| rs181808548 | 0.54406 | 5 |
| rs918859474 | 0.58955 | 5 |
| rs1380966191 | 0.58955 | 5 |
| rs960904276 | 0.57381 | 5 |
| rs990036357 | 0.56658 | 5 |
| rs1383486621 | 0.59359 | 5 |
| rs185221390 | 0.52742 | 3a |
| rs1288629572 | 0.85988 | 3a |
| rs992178675 | 0.63796 | 2b |
| rs1434392052 | 0.32745 | 2b |
| rs748591907 | 0.82852 | 2b |
| rs1373433759 | 0.82852 | 2b |
| rs1416328694 | 0.70377 | 2b |
| rs1412193183 | 0.82852 | 2b |
| rs116804530 | 0.82852 | 2b |
| rs1199596477 | 0.82852 | 2b |
| rs915074408 | 0.69529 | 3a |
| rs946639566 | 0.79703 | 3a |
| rs1029098266 | 0.80555 | 3a |
| rs987641912 | 0.55835 | 2b |
| rs1160249930 | 0.83617 | 2b |
| rs1433022831 | 0.60906 | 4 |
| rs1559001454 | 0.60906 | 4 |
| rs1042683838 | 0.60906 | 4 |
| rs1020840161 | 0.60906 | 4 |
| rs902405364 | 0.60906 | 4 |
| rs1259793627 | 0.60906 | 4 |
| rs1186439062 | 0.60906 | 4 |
| rs1486511037 | 0.60906 | 4 |
| rs944374411 | 0.60906 | 4 |
| rs755752702 | 0.60906 | 4 |
| rs879696627 | 0.60906 | 4 |
| rs1345243082 | 0.60906 | 4 |
| rs1244494846 | 0.60906 | 4 |
| rs1313406203 | 0.60906 | 4 |
| rs944203048 | 0.74401 | 4 |
| rs578071388 | 0.82206 | 2b |
| rs902756687 | 0.79283 | 2b |
| rs999207177 | 0.26 | 2a |
| rs148494543 | 0.85 | 2a |
| rs1393979820 | 0.60906 | 4 |
| rs3088307 | 0.55436 | 1f |
| rs969015549 | 0.60906 | 4 |
| rs978426115 | 0.66864 | 3a |
| rs1272621503 | 0.60906 | 4 |
| rs1175133175 | 0.60906 | 4 |
| rs1245907638 | 0.60906 | 4 |
| rs981897633 | 0.60906 | 4 |
| rs142693004 | 0.60906 | 4 |
| rs1384263900 | 0.60906 | 4 |
| rs542532827 | 0.60906 | 4 |
| rs1168861525 | 0.60906 | 4 |
| rs146017620 | 0.60906 | 4 |
| rs1445081648 | 0.60906 | 4 |
| rs961912738 | 0.60906 | 4 |
| rs972722568 | 0.60906 | 4 |
| rs1398023780 | 0.60906 | 4 |
| rs1325767181 | 0.60906 | 4 |
| rs28930678 | 0.60906 | 4 |
| rs1443115055 | 0.60906 | 4 |
| rs1423926011 | 0.70497 | 4 |
| rs922782733 | 0.82852 | 2b |
| rs932821143 | 0.38606 | 2b |
| rs750560218 | 0.70497 | 4 |
| rs139926849 | 0.60906 | 4 |
| rs1308984928 | 0.58955 | 5 |
| rs571416970 | 0.58955 | 5 |
| rs532137299 | 0.58955 | 5 |
| rs547454699 | 0.234 | 5 |
| rs1373058535 | 0.225 | 5 |
| rs1278530373 | 0.2769 | 5 |
| rs758675541 | 0.58955 | 5 |
| rs1442463937 | 0.58955 | 5 |
| rs1037029827 | 0.58955 | 5 |
| rs897159627 | 0.58955 | 5 |
| rs939462620 | 0.58955 | 5 |
| rs1056539629 | 0.58955 | 5 |
| rs1158994136 | 0.58955 | 5 |
| rs1474902590 | 0.58955 | 5 |
| rs868410365 | 0.9 | 5 |
| rs1165774038 | 0.85643 | 5 |
| rs1422126533 | 0.70087 | 5 |
| rs1267087229 | 0.27474 | 5 |
| rs1198763312 | 0.58955 | 5 |
| rs752964780 | 0.58955 | 5 |
| rs1271634891 | 0.58955 | 5 |
| rs902778052 | 0.58955 | 5 |
| rs1285389439 | 0.58955 | 5 |
| rs1164845107 | 0.58955 | 5 |

Table 5. Functionally significant 3 prime UTR SNP analysis by miRNASNP

|  | Position | Ref/Alt | Ref freq./  Alt freq. | Gain | Loss |
| --- | --- | --- | --- | --- | --- |
| [rs1272699435](https://guolab.wchscu.cn/miRNASNP/#!/snp?snp_id=rs1272699435&location=UTR3&one=1) | chr2:190969045 | A/G | 1/- | [5](https://guolab.wchscu.cn/miRNASNP/#!/snp?snp_id=rs1272699435&location=UTR3&four='1') | 0 |
| [rs1177647156](https://guolab.wchscu.cn/miRNASNP/#!/snp?snp_id=rs1177647156&location=UTR3&one=1) | chr2:190969048 | T/G | -/- | [4](https://guolab.wchscu.cn/miRNASNP/#!/snp?snp_id=rs1177647156&location=UTR3&four='1') | 0 |
| [rs1234425699](https://guolab.wchscu.cn/miRNASNP/#!/snp?snp_id=rs1234425699&location=UTR3&one=1) | chr2:190969054 | ACTT/A | -/- | 0 | [2](https://guolab.wchscu.cn/miRNASNP/#!/snp?snp_id=rs1234425699&location=UTR3&five='1') |
| [rs1014957291](https://guolab.wchscu.cn/miRNASNP/#!/snp?snp_id=rs1014957291&location=UTR3&one=1) | chr2:190969055 | C/A | 1/- | [1](https://guolab.wchscu.cn/miRNASNP/#!/snp?snp_id=rs1014957291&location=UTR3&four='1') | [8](https://guolab.wchscu.cn/miRNASNP/#!/snp?snp_id=rs1014957291&location=UTR3&five='1') |
| [rs1014957291](https://guolab.wchscu.cn/miRNASNP/#!/snp?snp_id=rs1014957291&location=UTR3&one=1) | chr2:190969055 | C/G | 1/- | [7](https://guolab.wchscu.cn/miRNASNP/#!/snp?snp_id=rs1014957291&location=UTR3&four='1') | [8](https://guolab.wchscu.cn/miRNASNP/#!/snp?snp_id=rs1014957291&location=UTR3&five='1') |
| [rs1378143535](https://guolab.wchscu.cn/miRNASNP/#!/snp?snp_id=rs1378143535&location=UTR3&one=1) | chr2:190969059 | TTAAAAG/T | -/0.0004 | [2](https://guolab.wchscu.cn/miRNASNP/#!/snp?snp_id=rs1378143535&location=UTR3&four='1') | [29](https://guolab.wchscu.cn/miRNASNP/#!/snp?snp_id=rs1378143535&location=UTR3&five='1') |
| [rs937020532](https://guolab.wchscu.cn/miRNASNP/#!/snp?snp_id=rs937020532&location=UTR3&one=1) | chr2:190969060 | TAAAAG/T | 1.0/- | [4](https://guolab.wchscu.cn/miRNASNP/#!/snp?snp_id=rs937020532&location=UTR3&four='1') | [31](https://guolab.wchscu.cn/miRNASNP/#!/snp?snp_id=rs937020532&location=UTR3&five='1') |
| [rs1159694777](https://guolab.wchscu.cn/miRNASNP/#!/snp?snp_id=rs1159694777&location=UTR3&one=1) | chr2:190969063 | A/AAT | -/- | [1](https://guolab.wchscu.cn/miRNASNP/#!/snp?snp_id=rs1159694777&location=UTR3&four='1') | [30](https://guolab.wchscu.cn/miRNASNP/#!/snp?snp_id=rs1159694777&location=UTR3&five='1') |
| [rs1055492520](https://guolab.wchscu.cn/miRNASNP/#!/snp?snp_id=rs1055492520&location=UTR3&one=1) | chr2:190969065 | G/T | 1/- | [1](https://guolab.wchscu.cn/miRNASNP/#!/snp?snp_id=rs1055492520&location=UTR3&four='1') | [30](https://guolab.wchscu.cn/miRNASNP/#!/snp?snp_id=rs1055492520&location=UTR3&five='1') |
| [rs573263994](https://guolab.wchscu.cn/miRNASNP/#!/snp?snp_id=rs573263994&location=UTR3&one=1) | chr2:190969066 | T/C | 1/- | [8](https://guolab.wchscu.cn/miRNASNP/#!/snp?snp_id=rs573263994&location=UTR3&four='1') | [30](https://guolab.wchscu.cn/miRNASNP/#!/snp?snp_id=rs573263994&location=UTR3&five='1') |
| [rs1391643601](https://guolab.wchscu.cn/miRNASNP/#!/snp?snp_id=rs1391643601&location=UTR3&one=1) | chr2:190969067 | A/C | 1/- | [5](https://guolab.wchscu.cn/miRNASNP/#!/snp?snp_id=rs1391643601&location=UTR3&four='1') | [27](https://guolab.wchscu.cn/miRNASNP/#!/snp?snp_id=rs1391643601&location=UTR3&five='1') |
| [rs1464989664](https://guolab.wchscu.cn/miRNASNP/#!/snp?snp_id=rs1464989664&location=UTR3&one=1) | chr2:190969069 | AGT/A | -/- | [6](https://guolab.wchscu.cn/miRNASNP/#!/snp?snp_id=rs1464989664&location=UTR3&four='1') | [6](https://guolab.wchscu.cn/miRNASNP/#!/snp?snp_id=rs1464989664&location=UTR3&five='1') |
| [rs971816169](https://guolab.wchscu.cn/miRNASNP/#!/snp?snp_id=rs971816169&location=UTR3&one=1) | chr2:190969074 | C/G | 1/- | [1](https://guolab.wchscu.cn/miRNASNP/#!/snp?snp_id=rs971816169&location=UTR3&four='1') | [3](https://guolab.wchscu.cn/miRNASNP/#!/snp?snp_id=rs971816169&location=UTR3&five='1') |
| [rs971816169](https://guolab.wchscu.cn/miRNASNP/#!/snp?snp_id=rs971816169&location=UTR3&one=1) | chr2:190969074 | C/T | 1/- | [1](https://guolab.wchscu.cn/miRNASNP/#!/snp?snp_id=rs971816169&location=UTR3&four='1') | [3](https://guolab.wchscu.cn/miRNASNP/#!/snp?snp_id=rs971816169&location=UTR3&five='1') |
| [rs886055373](https://guolab.wchscu.cn/miRNASNP/#!/snp?snp_id=rs886055373&location=UTR3&one=1) | chr2:190969075 | T/C | 1/- | [12](https://guolab.wchscu.cn/miRNASNP/#!/snp?snp_id=rs886055373&location=UTR3&four='1') | [3](https://guolab.wchscu.cn/miRNASNP/#!/snp?snp_id=rs886055373&location=UTR3&five='1') |
| [rs981440509](https://guolab.wchscu.cn/miRNASNP/#!/snp?snp_id=rs981440509&location=UTR3&one=1) | chr2:190969076 | T/C | 0.9999/0.0001 | [5](https://guolab.wchscu.cn/miRNASNP/#!/snp?snp_id=rs981440509&location=UTR3&four='1') | [1](https://guolab.wchscu.cn/miRNASNP/#!/snp?snp_id=rs981440509&location=UTR3&five='1') |
| [rs1408499050](https://guolab.wchscu.cn/miRNASNP/#!/snp?snp_id=rs1408499050&location=UTR3&one=1) | chr2:190969079 | C/T | 1/- | [4](https://guolab.wchscu.cn/miRNASNP/#!/snp?snp_id=rs1408499050&location=UTR3&four='1') | [3](https://guolab.wchscu.cn/miRNASNP/#!/snp?snp_id=rs1408499050&location=UTR3&five='1') |
| [rs887032302](https://guolab.wchscu.cn/miRNASNP/#!/snp?snp_id=rs887032302&location=UTR3&one=1) | chr2:190969080 | T/C | 1/- | [11](https://guolab.wchscu.cn/miRNASNP/#!/snp?snp_id=rs887032302&location=UTR3&four='1') | [5](https://guolab.wchscu.cn/miRNASNP/#!/snp?snp_id=rs887032302&location=UTR3&five='1') |
| [rs1005845582](https://guolab.wchscu.cn/miRNASNP/#!/snp?snp_id=rs1005845582&location=UTR3&one=1) | chr2:190969082 | G/C | 1/- | [10](https://guolab.wchscu.cn/miRNASNP/#!/snp?snp_id=rs1005845582&location=UTR3&four='1') | [14](https://guolab.wchscu.cn/miRNASNP/#!/snp?snp_id=rs1005845582&location=UTR3&five='1') |
| [rs1016841716](https://guolab.wchscu.cn/miRNASNP/#!/snp?snp_id=rs1016841716&location=UTR3&one=1) | chr2:190969085 | C/A | 1/- | [6](https://guolab.wchscu.cn/miRNASNP/#!/snp?snp_id=rs1016841716&location=UTR3&four='1') | [12](https://guolab.wchscu.cn/miRNASNP/#!/snp?snp_id=rs1016841716&location=UTR3&five='1') |
| [rs1214408280](https://guolab.wchscu.cn/miRNASNP/#!/snp?snp_id=rs1214408280&location=UTR3&one=1) | chr2:190969088 | T/C | 1/- | [14](https://guolab.wchscu.cn/miRNASNP/#!/snp?snp_id=rs1214408280&location=UTR3&four='1') | [1](https://guolab.wchscu.cn/miRNASNP/#!/snp?snp_id=rs1214408280&location=UTR3&five='1') |
| [rs899198125](https://guolab.wchscu.cn/miRNASNP/#!/snp?snp_id=rs899198125&location=UTR3&one=1) | chr2:190969098 | C/T | -/- | [4](https://guolab.wchscu.cn/miRNASNP/#!/snp?snp_id=rs899198125&location=UTR3&four='1') | [3](https://guolab.wchscu.cn/miRNASNP/#!/snp?snp_id=rs899198125&location=UTR3&five='1') |
| [rs757741528](https://guolab.wchscu.cn/miRNASNP/#!/snp?snp_id=rs757741528&location=UTR3&one=1) | chr2:190969100 | A/G | -/- | [2](https://guolab.wchscu.cn/miRNASNP/#!/snp?snp_id=rs757741528&location=UTR3&four='1') | [3](https://guolab.wchscu.cn/miRNASNP/#!/snp?snp_id=rs757741528&location=UTR3&five='1') |
| [rs537683771](https://guolab.wchscu.cn/miRNASNP/#!/snp?snp_id=rs537683771&location=UTR3&one=1) | chr2:190969101 | T/C | -/- | [1](https://guolab.wchscu.cn/miRNASNP/#!/snp?snp_id=rs537683771&location=UTR3&four='1') | [2](https://guolab.wchscu.cn/miRNASNP/#!/snp?snp_id=rs537683771&location=UTR3&five='1') |
| [rs779421003](https://guolab.wchscu.cn/miRNASNP/#!/snp?snp_id=rs779421003&location=UTR3&one=1) | chr2:190969104 | C/T | 1/- | [8](https://guolab.wchscu.cn/miRNASNP/#!/snp?snp_id=rs779421003&location=UTR3&four='1') | 0 |
| [rs1259509774](https://guolab.wchscu.cn/miRNASNP/#!/snp?snp_id=rs1259509774&location=UTR3&one=1) | chr2:190969105 | G/A | 1/- | [2](https://guolab.wchscu.cn/miRNASNP/#!/snp?snp_id=rs1259509774&location=UTR3&four='1') | [1](https://guolab.wchscu.cn/miRNASNP/#!/snp?snp_id=rs1259509774&location=UTR3&five='1') |
| [rs937373338](https://guolab.wchscu.cn/miRNASNP/#!/snp?snp_id=rs937373338&location=UTR3&one=1) | chr2:190969107 | A/G | 1/- | [1](https://guolab.wchscu.cn/miRNASNP/#!/snp?snp_id=rs937373338&location=UTR3&four='1') | [2](https://guolab.wchscu.cn/miRNASNP/#!/snp?snp_id=rs937373338&location=UTR3&five='1') |
| [rs990685172](https://guolab.wchscu.cn/miRNASNP/#!/snp?snp_id=rs990685172&location=UTR3&one=1) | chr2:190969122 | A/C | -/- | [2](https://guolab.wchscu.cn/miRNASNP/#!/snp?snp_id=rs990685172&location=UTR3&four='1') | [10](https://guolab.wchscu.cn/miRNASNP/#!/snp?snp_id=rs990685172&location=UTR3&five='1') |
| [rs1317862332](https://guolab.wchscu.cn/miRNASNP/#!/snp?snp_id=rs1317862332&location=UTR3&one=1) | chr2:190969125 | G/A | 1/- | [21](https://guolab.wchscu.cn/miRNASNP/#!/snp?snp_id=rs1317862332&location=UTR3&four='1') | [5](https://guolab.wchscu.cn/miRNASNP/#!/snp?snp_id=rs1317862332&location=UTR3&five='1') |
| [rs1317862332](https://guolab.wchscu.cn/miRNASNP/#!/snp?snp_id=rs1317862332&location=UTR3&one=1) | chr2:190969125 | G/T | 1/- | [2](https://guolab.wchscu.cn/miRNASNP/#!/snp?snp_id=rs1317862332&location=UTR3&four='1') | [5](https://guolab.wchscu.cn/miRNASNP/#!/snp?snp_id=rs1317862332&location=UTR3&five='1') |
| [rs1241847098](https://guolab.wchscu.cn/miRNASNP/#!/snp?snp_id=rs1241847098&location=UTR3&one=1) | chr2:190969129 | T/C | 1/- | [2](https://guolab.wchscu.cn/miRNASNP/#!/snp?snp_id=rs1241847098&location=UTR3&four='1') | [5](https://guolab.wchscu.cn/miRNASNP/#!/snp?snp_id=rs1241847098&location=UTR3&five='1') |
| [rs752398249](https://guolab.wchscu.cn/miRNASNP/#!/snp?snp_id=rs752398249&location=UTR3&one=1) | chr2:190969137 | A/G | -/- | [3](https://guolab.wchscu.cn/miRNASNP/#!/snp?snp_id=rs752398249&location=UTR3&four='1') | [1](https://guolab.wchscu.cn/miRNASNP/#!/snp?snp_id=rs752398249&location=UTR3&five='1') |
| [rs915074408](https://guolab.wchscu.cn/miRNASNP/#!/snp?snp_id=rs915074408&location=UTR3&one=1) | chr2:190969156 | C/T | 1/- | [6](https://guolab.wchscu.cn/miRNASNP/#!/snp?snp_id=rs915074408&location=UTR3&four='1') | [4](https://guolab.wchscu.cn/miRNASNP/#!/snp?snp_id=rs915074408&location=UTR3&five='1') |
| [rs946639566](https://guolab.wchscu.cn/miRNASNP/#!/snp?snp_id=rs946639566&location=UTR3&one=1) | chr2:190969165 | T/G | -/- | [4](https://guolab.wchscu.cn/miRNASNP/#!/snp?snp_id=rs946639566&location=UTR3&four='1') | 0 |
| [rs1029098266](https://guolab.wchscu.cn/miRNASNP/#!/snp?snp_id=rs1029098266&location=UTR3&one=1) | chr2:190969168 | A/C | 1/- | [2](https://guolab.wchscu.cn/miRNASNP/#!/snp?snp_id=rs1029098266&location=UTR3&four='1') | [2](https://guolab.wchscu.cn/miRNASNP/#!/snp?snp_id=rs1029098266&location=UTR3&five='1') |
| [rs148970951](https://guolab.wchscu.cn/miRNASNP/#!/snp?snp_id=rs148970951&location=UTR3&one=1) | chr2:190969176 | CTTCT/C | -/0.0057 | [1](https://guolab.wchscu.cn/miRNASNP/#!/snp?snp_id=rs148970951&location=UTR3&four='1') | [3](https://guolab.wchscu.cn/miRNASNP/#!/snp?snp_id=rs148970951&location=UTR3&five='1') |
| [rs1360916619](https://guolab.wchscu.cn/miRNASNP/#!/snp?snp_id=rs1360916619&location=UTR3&one=1) | chr2:190969179 | CTTTG/C | -/0.0001 | 0 | [1](https://guolab.wchscu.cn/miRNASNP/#!/snp?snp_id=rs1360916619&location=UTR3&five='1') |
| [rs987641912](https://guolab.wchscu.cn/miRNASNP/#!/snp?snp_id=rs987641912&location=UTR3&one=1) | chr2:190969179 | C/A | 0.9998/- | [2](https://guolab.wchscu.cn/miRNASNP/#!/snp?snp_id=rs987641912&location=UTR3&four='1') | [3](https://guolab.wchscu.cn/miRNASNP/#!/snp?snp_id=rs987641912&location=UTR3&five='1') |
| [rs987641912](https://guolab.wchscu.cn/miRNASNP/#!/snp?snp_id=rs987641912&location=UTR3&one=1) | chr2:190969179 | C/G | 0.9998/0.0001 | [2](https://guolab.wchscu.cn/miRNASNP/#!/snp?snp_id=rs987641912&location=UTR3&four='1') | [3](https://guolab.wchscu.cn/miRNASNP/#!/snp?snp_id=rs987641912&location=UTR3&five='1') |
| [rs987641912](https://guolab.wchscu.cn/miRNASNP/#!/snp?snp_id=rs987641912&location=UTR3&one=1) | chr2:190969179 | C/T | 0.9998/- | [3](https://guolab.wchscu.cn/miRNASNP/#!/snp?snp_id=rs987641912&location=UTR3&four='1') | [3](https://guolab.wchscu.cn/miRNASNP/#!/snp?snp_id=rs987641912&location=UTR3&five='1') |
| [rs1160249930](https://guolab.wchscu.cn/miRNASNP/#!/snp?snp_id=rs1160249930&location=UTR3&one=1) | chr2:190969185 | T/G | 1/- | [3](https://guolab.wchscu.cn/miRNASNP/#!/snp?snp_id=rs1160249930&location=UTR3&four='1') | 0 |
| [rs1433022831](https://guolab.wchscu.cn/miRNASNP/#!/snp?snp_id=rs1433022831&location=UTR3&one=1) | chr2:190969189 | T/C | 1/- | [5](https://guolab.wchscu.cn/miRNASNP/#!/snp?snp_id=rs1433022831&location=UTR3&four='1') | 0 |
| [rs1042683838](https://guolab.wchscu.cn/miRNASNP/#!/snp?snp_id=rs1042683838&location=UTR3&one=1) | chr2:190969201 | T/C | 1/- | [1](https://guolab.wchscu.cn/miRNASNP/#!/snp?snp_id=rs1042683838&location=UTR3&four='1') | 0 |
| [rs1020840161](https://guolab.wchscu.cn/miRNASNP/#!/snp?snp_id=rs1020840161&location=UTR3&one=1) | chr2:190969202 | T/C | 1/- | [1](https://guolab.wchscu.cn/miRNASNP/#!/snp?snp_id=rs1020840161&location=UTR3&four='1') | 0 |
| [rs902405364](https://guolab.wchscu.cn/miRNASNP/#!/snp?snp_id=rs902405364&location=UTR3&one=1) | chr2:190969212 | A/G | 1/- | [3](https://guolab.wchscu.cn/miRNASNP/#!/snp?snp_id=rs902405364&location=UTR3&four='1') | [1](https://guolab.wchscu.cn/miRNASNP/#!/snp?snp_id=rs902405364&location=UTR3&five='1') |
| [rs1259793627](https://guolab.wchscu.cn/miRNASNP/#!/snp?snp_id=rs1259793627&location=UTR3&one=1) | chr2:190969217 | A/T | 1/- | 0 | [3](https://guolab.wchscu.cn/miRNASNP/#!/snp?snp_id=rs1259793627&location=UTR3&five='1') |
| [rs1186439062](https://guolab.wchscu.cn/miRNASNP/#!/snp?snp_id=rs1186439062&location=UTR3&one=1) | chr2:190969219 | C/T | 1/- | [5](https://guolab.wchscu.cn/miRNASNP/#!/snp?snp_id=rs1186439062&location=UTR3&four='1') | [2](https://guolab.wchscu.cn/miRNASNP/#!/snp?snp_id=rs1186439062&location=UTR3&five='1') |
| [rs1241131631](https://guolab.wchscu.cn/miRNASNP/#!/snp?snp_id=rs1241131631&location=UTR3&one=1) | chr2:190969224 | GT/G | -/- | 0 | [4](https://guolab.wchscu.cn/miRNASNP/#!/snp?snp_id=rs1241131631&location=UTR3&five='1') |
| [rs1486511037](https://guolab.wchscu.cn/miRNASNP/#!/snp?snp_id=rs1486511037&location=UTR3&one=1) | chr2:190969228 | T/C | 1/- | [3](https://guolab.wchscu.cn/miRNASNP/#!/snp?snp_id=rs1486511037&location=UTR3&four='1') | [5](https://guolab.wchscu.cn/miRNASNP/#!/snp?snp_id=rs1486511037&location=UTR3&five='1') |
| [rs944374411](https://guolab.wchscu.cn/miRNASNP/#!/snp?snp_id=rs944374411&location=UTR3&one=1) | chr2:190969229 | C/T | 0.9998/0.0002 | [7](https://guolab.wchscu.cn/miRNASNP/#!/snp?snp_id=rs944374411&location=UTR3&four='1') | [2](https://guolab.wchscu.cn/miRNASNP/#!/snp?snp_id=rs944374411&location=UTR3&five='1') |
| [rs755752702](https://guolab.wchscu.cn/miRNASNP/#!/snp?snp_id=rs755752702&location=UTR3&one=1) | chr2:190969230 | G/A | 1/- | [7](https://guolab.wchscu.cn/miRNASNP/#!/snp?snp_id=rs755752702&location=UTR3&four='1') | [1](https://guolab.wchscu.cn/miRNASNP/#!/snp?snp_id=rs755752702&location=UTR3&five='1') |
| [rs879696627](https://guolab.wchscu.cn/miRNASNP/#!/snp?snp_id=rs879696627&location=UTR3&one=1) | chr2:190969231 | C/G | -/- | [4](https://guolab.wchscu.cn/miRNASNP/#!/snp?snp_id=rs879696627&location=UTR3&four='1') | [1](https://guolab.wchscu.cn/miRNASNP/#!/snp?snp_id=rs879696627&location=UTR3&five='1') |
| [rs1345243082](https://guolab.wchscu.cn/miRNASNP/#!/snp?snp_id=rs1345243082&location=UTR3&one=1) | chr2:190969233 | A/T | 1/- | [8](https://guolab.wchscu.cn/miRNASNP/#!/snp?snp_id=rs1345243082&location=UTR3&four='1') | [1](https://guolab.wchscu.cn/miRNASNP/#!/snp?snp_id=rs1345243082&location=UTR3&five='1') |
| [rs1244494846](https://guolab.wchscu.cn/miRNASNP/#!/snp?snp_id=rs1244494846&location=UTR3&one=1) | chr2:190969236 | T/C | -/- | [1](https://guolab.wchscu.cn/miRNASNP/#!/snp?snp_id=rs1244494846&location=UTR3&four='1') | [1](https://guolab.wchscu.cn/miRNASNP/#!/snp?snp_id=rs1244494846&location=UTR3&five='1') |
| [rs1279188138](https://guolab.wchscu.cn/miRNASNP/#!/snp?snp_id=rs1279188138&location=UTR3&one=1) | chr2:190969240 | TA/T | -/- | 0 | [12](https://guolab.wchscu.cn/miRNASNP/#!/snp?snp_id=rs1279188138&location=UTR3&five='1') |
| [rs1311321881](https://guolab.wchscu.cn/miRNASNP/#!/snp?snp_id=rs1311321881&location=UTR3&one=1) | chr2:190969246 | CTG/C | 1.0/- | [5](https://guolab.wchscu.cn/miRNASNP/#!/snp?snp_id=rs1311321881&location=UTR3&four='1') | [12](https://guolab.wchscu.cn/miRNASNP/#!/snp?snp_id=rs1311321881&location=UTR3&five='1') |
| [rs1313406203](https://guolab.wchscu.cn/miRNASNP/#!/snp?snp_id=rs1313406203&location=UTR3&one=1) | chr2:190969248 | G/A | 1/- | [3](https://guolab.wchscu.cn/miRNASNP/#!/snp?snp_id=rs1313406203&location=UTR3&four='1') | [5](https://guolab.wchscu.cn/miRNASNP/#!/snp?snp_id=rs1313406203&location=UTR3&five='1') |
| [rs979463051](https://guolab.wchscu.cn/miRNASNP/#!/snp?snp_id=rs979463051&location=UTR3&one=1) | chr2:190969253 | A/G | 1/- | [17](https://guolab.wchscu.cn/miRNASNP/#!/snp?snp_id=rs979463051&location=UTR3&four='1') | [9](https://guolab.wchscu.cn/miRNASNP/#!/snp?snp_id=rs979463051&location=UTR3&five='1') |
| [rs1040006583](https://guolab.wchscu.cn/miRNASNP/#!/snp?snp_id=rs1040006583&location=UTR3&one=1) | chr2:190969255 | G/A | -/- | [8](https://guolab.wchscu.cn/miRNASNP/#!/snp?snp_id=rs1040006583&location=UTR3&four='1') | [11](https://guolab.wchscu.cn/miRNASNP/#!/snp?snp_id=rs1040006583&location=UTR3&five='1') |
| [rs746484042](https://guolab.wchscu.cn/miRNASNP/#!/snp?snp_id=rs746484042&location=UTR3&one=1) | chr2:190969256 | A/C | -/- | [6](https://guolab.wchscu.cn/miRNASNP/#!/snp?snp_id=rs746484042&location=UTR3&four='1') | [13](https://guolab.wchscu.cn/miRNASNP/#!/snp?snp_id=rs746484042&location=UTR3&five='1') |
| [rs900296058](https://guolab.wchscu.cn/miRNASNP/#!/snp?snp_id=rs900296058&location=UTR3&one=1) | chr2:190969262 | C/G | 0.9998/0.0002 | [4](https://guolab.wchscu.cn/miRNASNP/#!/snp?snp_id=rs900296058&location=UTR3&four='1') | [5](https://guolab.wchscu.cn/miRNASNP/#!/snp?snp_id=rs900296058&location=UTR3&five='1') |
| [rs937037547](https://guolab.wchscu.cn/miRNASNP/#!/snp?snp_id=rs937037547&location=UTR3&one=1) | chr2:190969263 | A/G | 1/- | [1](https://guolab.wchscu.cn/miRNASNP/#!/snp?snp_id=rs937037547&location=UTR3&four='1') | [3](https://guolab.wchscu.cn/miRNASNP/#!/snp?snp_id=rs937037547&location=UTR3&five='1') |
| [rs1299766797](https://guolab.wchscu.cn/miRNASNP/#!/snp?snp_id=rs1299766797&location=UTR3&one=1) | chr2:190969264 | G/A | 1/- | [5](https://guolab.wchscu.cn/miRNASNP/#!/snp?snp_id=rs1299766797&location=UTR3&four='1') | [8](https://guolab.wchscu.cn/miRNASNP/#!/snp?snp_id=rs1299766797&location=UTR3&five='1') |
| [rs991085331](https://guolab.wchscu.cn/miRNASNP/#!/snp?snp_id=rs991085331&location=UTR3&one=1) | chr2:190969271 | T/C | 1/- | 0 | [3](https://guolab.wchscu.cn/miRNASNP/#!/snp?snp_id=rs991085331&location=UTR3&five='1') |
| [rs1374679950](https://guolab.wchscu.cn/miRNASNP/#!/snp?snp_id=rs1374679950&location=UTR3&one=1) | chr2:190969281 | C/T | 1/- | [6](https://guolab.wchscu.cn/miRNASNP/#!/snp?snp_id=rs1374679950&location=UTR3&four='1') | [3](https://guolab.wchscu.cn/miRNASNP/#!/snp?snp_id=rs1374679950&location=UTR3&five='1') |
| [rs995881201](https://guolab.wchscu.cn/miRNASNP/#!/snp?snp_id=rs995881201&location=UTR3&one=1) | chr2:190969285 | A/G | -/- | [8](https://guolab.wchscu.cn/miRNASNP/#!/snp?snp_id=rs995881201&location=UTR3&four='1') | [3](https://guolab.wchscu.cn/miRNASNP/#!/snp?snp_id=rs995881201&location=UTR3&five='1') |
| [rs542136528](https://guolab.wchscu.cn/miRNASNP/#!/snp?snp_id=rs542136528&location=UTR3&one=1) | chr2:190969286 | A/G | -/- | [11](https://guolab.wchscu.cn/miRNASNP/#!/snp?snp_id=rs542136528&location=UTR3&four='1') | [3](https://guolab.wchscu.cn/miRNASNP/#!/snp?snp_id=rs542136528&location=UTR3&five='1') |
| [rs917014338](https://guolab.wchscu.cn/miRNASNP/#!/snp?snp_id=rs917014338&location=UTR3&one=1) | chr2:190969293 | T/G | 1/- | [6](https://guolab.wchscu.cn/miRNASNP/#!/snp?snp_id=rs917014338&location=UTR3&four='1') | [6](https://guolab.wchscu.cn/miRNASNP/#!/snp?snp_id=rs917014338&location=UTR3&five='1') |
| [rs563782303](https://guolab.wchscu.cn/miRNASNP/#!/snp?snp_id=rs563782303&location=UTR3&one=1) | chr2:190969298 | A/G | -/- | [2](https://guolab.wchscu.cn/miRNASNP/#!/snp?snp_id=rs563782303&location=UTR3&four='1') | [4](https://guolab.wchscu.cn/miRNASNP/#!/snp?snp_id=rs563782303&location=UTR3&five='1') |
| [rs1257637616](https://guolab.wchscu.cn/miRNASNP/#!/snp?snp_id=rs1257637616&location=UTR3&one=1) | chr2:190969303 | A/G | -/- | [3](https://guolab.wchscu.cn/miRNASNP/#!/snp?snp_id=rs1257637616&location=UTR3&four='1') | [3](https://guolab.wchscu.cn/miRNASNP/#!/snp?snp_id=rs1257637616&location=UTR3&five='1') |
| [rs11305](https://guolab.wchscu.cn/miRNASNP/#!/snp?snp_id=rs11305&location=UTR3&one=1) | chr2:190969304 | T/C | 0.9593/0.0407 | [4](https://guolab.wchscu.cn/miRNASNP/#!/snp?snp_id=rs11305&location=UTR3&four='1') | [2](https://guolab.wchscu.cn/miRNASNP/#!/snp?snp_id=rs11305&location=UTR3&five='1') |
| [rs1046888151](https://guolab.wchscu.cn/miRNASNP/#!/snp?snp_id=rs1046888151&location=UTR3&one=1) | chr2:190969313 | C/A | -/- | [1](https://guolab.wchscu.cn/miRNASNP/#!/snp?snp_id=rs1046888151&location=UTR3&four='1') | [3](https://guolab.wchscu.cn/miRNASNP/#!/snp?snp_id=rs1046888151&location=UTR3&five='1') |
| [rs887935841](https://guolab.wchscu.cn/miRNASNP/#!/snp?snp_id=rs887935841&location=UTR3&one=1) | chr2:190969315 | A/G | -/- | 0 | [1](https://guolab.wchscu.cn/miRNASNP/#!/snp?snp_id=rs887935841&location=UTR3&five='1') |
| [rs1235818401](https://guolab.wchscu.cn/miRNASNP/#!/snp?snp_id=rs1235818401&location=UTR3&one=1) | chr2:190969316 | TAC/T | -/- | [2](https://guolab.wchscu.cn/miRNASNP/#!/snp?snp_id=rs1235818401&location=UTR3&four='1') | [1](https://guolab.wchscu.cn/miRNASNP/#!/snp?snp_id=rs1235818401&location=UTR3&five='1') |
| [rs886973377](https://guolab.wchscu.cn/miRNASNP/#!/snp?snp_id=rs886973377&location=UTR3&one=1) | chr2:190969317 | A/G | 1/- | [2](https://guolab.wchscu.cn/miRNASNP/#!/snp?snp_id=rs886973377&location=UTR3&four='1') | [1](https://guolab.wchscu.cn/miRNASNP/#!/snp?snp_id=rs886973377&location=UTR3&five='1') |
| [rs1478394175](https://guolab.wchscu.cn/miRNASNP/#!/snp?snp_id=rs1478394175&location=UTR3&one=1) | chr2:190969327 | T/A | 1/- | [2](https://guolab.wchscu.cn/miRNASNP/#!/snp?snp_id=rs1478394175&location=UTR3&four='1') | [1](https://guolab.wchscu.cn/miRNASNP/#!/snp?snp_id=rs1478394175&location=UTR3&five='1') |
| [rs1266013997](https://guolab.wchscu.cn/miRNASNP/#!/snp?snp_id=rs1266013997&location=UTR3&one=1) | chr2:190969340 | A/G | 1/- | [2](https://guolab.wchscu.cn/miRNASNP/#!/snp?snp_id=rs1266013997&location=UTR3&four='1') | 0 |
| [rs1186197063](https://guolab.wchscu.cn/miRNASNP/#!/snp?snp_id=rs1186197063&location=UTR3&one=1) | chr2:190969341 | A/T | 1/- | 0 | 0 |
| [rs941197964](https://guolab.wchscu.cn/miRNASNP/#!/snp?snp_id=rs941197964&location=UTR3&one=1) | chr2:190969344 | C/A | 1/- | [12](https://guolab.wchscu.cn/miRNASNP/#!/snp?snp_id=rs941197964&location=UTR3&four='1') | [6](https://guolab.wchscu.cn/miRNASNP/#!/snp?snp_id=rs941197964&location=UTR3&five='1') |
| [rs941197964](https://guolab.wchscu.cn/miRNASNP/#!/snp?snp_id=rs941197964&location=UTR3&one=1) | chr2:190969344 | C/G | 1/- | [2](https://guolab.wchscu.cn/miRNASNP/#!/snp?snp_id=rs941197964&location=UTR3&four='1') | [6](https://guolab.wchscu.cn/miRNASNP/#!/snp?snp_id=rs941197964&location=UTR3&five='1') |
| [rs1005025126](https://guolab.wchscu.cn/miRNASNP/#!/snp?snp_id=rs1005025126&location=UTR3&one=1) | chr2:190969356 | A/T | -/- | [2](https://guolab.wchscu.cn/miRNASNP/#!/snp?snp_id=rs1005025126&location=UTR3&four='1') | [1](https://guolab.wchscu.cn/miRNASNP/#!/snp?snp_id=rs1005025126&location=UTR3&five='1') |
| [rs1200665369](https://guolab.wchscu.cn/miRNASNP/#!/snp?snp_id=rs1200665369&location=UTR3&one=1) | chr2:190969366 | T/G | 1/- | [1](https://guolab.wchscu.cn/miRNASNP/#!/snp?snp_id=rs1200665369&location=UTR3&four='1') | [2](https://guolab.wchscu.cn/miRNASNP/#!/snp?snp_id=rs1200665369&location=UTR3&five='1') |
| [rs1484038511](https://guolab.wchscu.cn/miRNASNP/#!/snp?snp_id=rs1484038511&location=UTR3&one=1) | chr2:190969369 | A/G | 1/- | [1](https://guolab.wchscu.cn/miRNASNP/#!/snp?snp_id=rs1484038511&location=UTR3&four='1') | [2](https://guolab.wchscu.cn/miRNASNP/#!/snp?snp_id=rs1484038511&location=UTR3&five='1') |
| [rs1249044418](https://guolab.wchscu.cn/miRNASNP/#!/snp?snp_id=rs1249044418&location=UTR3&one=1) | chr2:190969372 | A/T | 1/- | 0 | [2](https://guolab.wchscu.cn/miRNASNP/#!/snp?snp_id=rs1249044418&location=UTR3&five='1') |
| [rs1038714220](https://guolab.wchscu.cn/miRNASNP/#!/snp?snp_id=rs1038714220&location=UTR3&one=1) | chr2:190969383 | C/A | 1/- | [5](https://guolab.wchscu.cn/miRNASNP/#!/snp?snp_id=rs1038714220&location=UTR3&four='1') | [3](https://guolab.wchscu.cn/miRNASNP/#!/snp?snp_id=rs1038714220&location=UTR3&five='1') |
| [rs1038714220](https://guolab.wchscu.cn/miRNASNP/#!/snp?snp_id=rs1038714220&location=UTR3&one=1) | chr2:190969383 | C/T | 1/- | [1](https://guolab.wchscu.cn/miRNASNP/#!/snp?snp_id=rs1038714220&location=UTR3&four='1') | [3](https://guolab.wchscu.cn/miRNASNP/#!/snp?snp_id=rs1038714220&location=UTR3&five='1') |
| [rs368576923](https://guolab.wchscu.cn/miRNASNP/#!/snp?snp_id=rs368576923&location=UTR3&one=1) | chr2:190969386 | C/G | 0.9999/- | [6](https://guolab.wchscu.cn/miRNASNP/#!/snp?snp_id=rs368576923&location=UTR3&four='1') | [4](https://guolab.wchscu.cn/miRNASNP/#!/snp?snp_id=rs368576923&location=UTR3&five='1') |
| [rs368576923](https://guolab.wchscu.cn/miRNASNP/#!/snp?snp_id=rs368576923&location=UTR3&one=1) | chr2:190969386 | C/T | 0.9999/0.0001 | [4](https://guolab.wchscu.cn/miRNASNP/#!/snp?snp_id=rs368576923&location=UTR3&four='1') | [4](https://guolab.wchscu.cn/miRNASNP/#!/snp?snp_id=rs368576923&location=UTR3&five='1') |
| [rs996199996](https://guolab.wchscu.cn/miRNASNP/#!/snp?snp_id=rs996199996&location=UTR3&one=1) | chr2:190969393 | A/T | -/- | [1](https://guolab.wchscu.cn/miRNASNP/#!/snp?snp_id=rs996199996&location=UTR3&four='1') | [8](https://guolab.wchscu.cn/miRNASNP/#!/snp?snp_id=rs996199996&location=UTR3&five='1') |
| [rs1354581153](https://guolab.wchscu.cn/miRNASNP/#!/snp?snp_id=rs1354581153&location=UTR3&one=1) | chr2:190969394 | T/C | 1/- | 0 | [5](https://guolab.wchscu.cn/miRNASNP/#!/snp?snp_id=rs1354581153&location=UTR3&five='1') |
| [rs189030575](https://guolab.wchscu.cn/miRNASNP/#!/snp?snp_id=rs189030575&location=UTR3&one=1) | chr2:190969402 | T/G | 1/- | [4](https://guolab.wchscu.cn/miRNASNP/#!/snp?snp_id=rs189030575&location=UTR3&four='1') | [2](https://guolab.wchscu.cn/miRNASNP/#!/snp?snp_id=rs189030575&location=UTR3&five='1') |
| [rs1397943465](https://guolab.wchscu.cn/miRNASNP/#!/snp?snp_id=rs1397943465&location=UTR3&one=1) | chr2:190969407 | G/T | 1/- | [4](https://guolab.wchscu.cn/miRNASNP/#!/snp?snp_id=rs1397943465&location=UTR3&four='1') | [1](https://guolab.wchscu.cn/miRNASNP/#!/snp?snp_id=rs1397943465&location=UTR3&five='1') |
| [rs981856342](https://guolab.wchscu.cn/miRNASNP/#!/snp?snp_id=rs981856342&location=UTR3&one=1) | chr2:190969412 | A/G | 1/- | [7](https://guolab.wchscu.cn/miRNASNP/#!/snp?snp_id=rs981856342&location=UTR3&four='1') | [5](https://guolab.wchscu.cn/miRNASNP/#!/snp?snp_id=rs981856342&location=UTR3&five='1') |
| [rs1034344469](https://guolab.wchscu.cn/miRNASNP/#!/snp?snp_id=rs1034344469&location=UTR3&one=1) | chr2:190969413 | G/A | -/- | [5](https://guolab.wchscu.cn/miRNASNP/#!/snp?snp_id=rs1034344469&location=UTR3&four='1') | [4](https://guolab.wchscu.cn/miRNASNP/#!/snp?snp_id=rs1034344469&location=UTR3&five='1') |
| [rs958816905](https://guolab.wchscu.cn/miRNASNP/#!/snp?snp_id=rs958816905&location=UTR3&one=1) | chr2:190969414 | C/A | -/- | [8](https://guolab.wchscu.cn/miRNASNP/#!/snp?snp_id=rs958816905&location=UTR3&four='1') | [8](https://guolab.wchscu.cn/miRNASNP/#!/snp?snp_id=rs958816905&location=UTR3&five='1') |
| [rs1431755920](https://guolab.wchscu.cn/miRNASNP/#!/snp?snp_id=rs1431755920&location=UTR3&one=1) | chr2:190969417 | T/C | 1/- | [16](https://guolab.wchscu.cn/miRNASNP/#!/snp?snp_id=rs1431755920&location=UTR3&four='1') | [9](https://guolab.wchscu.cn/miRNASNP/#!/snp?snp_id=rs1431755920&location=UTR3&five='1') |
| [rs1466227575](https://guolab.wchscu.cn/miRNASNP/#!/snp?snp_id=rs1466227575&location=UTR3&one=1) | chr2:190969421 | G/A | 1/- | [1](https://guolab.wchscu.cn/miRNASNP/#!/snp?snp_id=rs1466227575&location=UTR3&four='1') | [11](https://guolab.wchscu.cn/miRNASNP/#!/snp?snp_id=rs1466227575&location=UTR3&five='1') |
| [rs1425931456](https://guolab.wchscu.cn/miRNASNP/#!/snp?snp_id=rs1425931456&location=UTR3&one=1) | chr2:190969423 | A/G | 1/- | [11](https://guolab.wchscu.cn/miRNASNP/#!/snp?snp_id=rs1425931456&location=UTR3&four='1') | [17](https://guolab.wchscu.cn/miRNASNP/#!/snp?snp_id=rs1425931456&location=UTR3&five='1') |
| [rs990213925](https://guolab.wchscu.cn/miRNASNP/#!/snp?snp_id=rs990213925&location=UTR3&one=1) | chr2:190969429 | A/G | -/- | [2](https://guolab.wchscu.cn/miRNASNP/#!/snp?snp_id=rs990213925&location=UTR3&four='1') | [1](https://guolab.wchscu.cn/miRNASNP/#!/snp?snp_id=rs990213925&location=UTR3&five='1') |
| [rs915105846](https://guolab.wchscu.cn/miRNASNP/#!/snp?snp_id=rs915105846&location=UTR3&one=1) | chr2:190969432 | A/G | -/- | [6](https://guolab.wchscu.cn/miRNASNP/#!/snp?snp_id=rs915105846&location=UTR3&four='1') | [1](https://guolab.wchscu.cn/miRNASNP/#!/snp?snp_id=rs915105846&location=UTR3&five='1') |
| [rs890554652](https://guolab.wchscu.cn/miRNASNP/#!/snp?snp_id=rs890554652&location=UTR3&one=1) | chr2:190969438 | CCA/C | -/- | [1](https://guolab.wchscu.cn/miRNASNP/#!/snp?snp_id=rs890554652&location=UTR3&four='1') | [2](https://guolab.wchscu.cn/miRNASNP/#!/snp?snp_id=rs890554652&location=UTR3&five='1') |
| [rs1190058198](https://guolab.wchscu.cn/miRNASNP/#!/snp?snp_id=rs1190058198&location=UTR3&one=1) | chr2:190969443 | TAGAC/T | -/- | [4](https://guolab.wchscu.cn/miRNASNP/#!/snp?snp_id=rs1190058198&location=UTR3&four='1') | [3](https://guolab.wchscu.cn/miRNASNP/#!/snp?snp_id=rs1190058198&location=UTR3&five='1') |
| [rs1369356450](https://guolab.wchscu.cn/miRNASNP/#!/snp?snp_id=rs1369356450&location=UTR3&one=1) | chr2:190969443 | T/TA | -/- | [5](https://guolab.wchscu.cn/miRNASNP/#!/snp?snp_id=rs1369356450&location=UTR3&four='1') | [1](https://guolab.wchscu.cn/miRNASNP/#!/snp?snp_id=rs1369356450&location=UTR3&five='1') |
| [rs886055374](https://guolab.wchscu.cn/miRNASNP/#!/snp?snp_id=rs886055374&location=UTR3&one=1) | chr2:190969443 | T/C | 0.9999/0.0001 | [2](https://guolab.wchscu.cn/miRNASNP/#!/snp?snp_id=rs886055374&location=UTR3&four='1') | [1](https://guolab.wchscu.cn/miRNASNP/#!/snp?snp_id=rs886055374&location=UTR3&five='1') |
| [rs557207722](https://guolab.wchscu.cn/miRNASNP/#!/snp?snp_id=rs557207722&location=UTR3&one=1) | chr2:190969447 | C/T | -/- | [3](https://guolab.wchscu.cn/miRNASNP/#!/snp?snp_id=rs557207722&location=UTR3&four='1') | [7](https://guolab.wchscu.cn/miRNASNP/#!/snp?snp_id=rs557207722&location=UTR3&five='1') |
| [rs977912956](https://guolab.wchscu.cn/miRNASNP/#!/snp?snp_id=rs977912956&location=UTR3&one=1) | chr2:190969549 | G/A | -/- | [3](https://guolab.wchscu.cn/miRNASNP/#!/snp?snp_id=rs977912956&location=UTR3&four='1') | [1](https://guolab.wchscu.cn/miRNASNP/#!/snp?snp_id=rs977912956&location=UTR3&five='1') |
| [rs1306532833](https://guolab.wchscu.cn/miRNASNP/#!/snp?snp_id=rs1306532833&location=UTR3&one=1) | chr2:190969555 | T/C | 1/- | [3](https://guolab.wchscu.cn/miRNASNP/#!/snp?snp_id=rs1306532833&location=UTR3&four='1') | [3](https://guolab.wchscu.cn/miRNASNP/#!/snp?snp_id=rs1306532833&location=UTR3&five='1') |
| [rs1033692335](https://guolab.wchscu.cn/miRNASNP/#!/snp?snp_id=rs1033692335&location=UTR3&one=1) | chr2:190969560 | C/T | 1/- | [1](https://guolab.wchscu.cn/miRNASNP/#!/snp?snp_id=rs1033692335&location=UTR3&four='1') | [4](https://guolab.wchscu.cn/miRNASNP/#!/snp?snp_id=rs1033692335&location=UTR3&five='1') |
| [rs1447801311](https://guolab.wchscu.cn/miRNASNP/#!/snp?snp_id=rs1447801311&location=UTR3&one=1) | chr2:190969565 | A/G | 1/- | 0 | [7](https://guolab.wchscu.cn/miRNASNP/#!/snp?snp_id=rs1447801311&location=UTR3&five='1') |
| [rs879538247](https://guolab.wchscu.cn/miRNASNP/#!/snp?snp_id=rs879538247&location=UTR3&one=1) | chr2:190969567 | A/G | 1/- | [1](https://guolab.wchscu.cn/miRNASNP/#!/snp?snp_id=rs879538247&location=UTR3&four='1') | [10](https://guolab.wchscu.cn/miRNASNP/#!/snp?snp_id=rs879538247&location=UTR3&five='1') |
| [rs1461320344](https://guolab.wchscu.cn/miRNASNP/#!/snp?snp_id=rs1461320344&location=UTR3&one=1) | chr2:190969570 | A/T | 1/- | [5](https://guolab.wchscu.cn/miRNASNP/#!/snp?snp_id=rs1461320344&location=UTR3&four='1') | [16](https://guolab.wchscu.cn/miRNASNP/#!/snp?snp_id=rs1461320344&location=UTR3&five='1') |
| [rs1392122056](https://guolab.wchscu.cn/miRNASNP/#!/snp?snp_id=rs1392122056&location=UTR3&one=1) | chr2:190969573 | G/A | 1/- | [28](https://guolab.wchscu.cn/miRNASNP/#!/snp?snp_id=rs1392122056&location=UTR3&four='1') | [16](https://guolab.wchscu.cn/miRNASNP/#!/snp?snp_id=rs1392122056&location=UTR3&five='1') |
| [rs1471371676](https://guolab.wchscu.cn/miRNASNP/#!/snp?snp_id=rs1471371676&location=UTR3&one=1) | chr2:190969579 | A/G | -/- | [10](https://guolab.wchscu.cn/miRNASNP/#!/snp?snp_id=rs1471371676&location=UTR3&four='1') | [7](https://guolab.wchscu.cn/miRNASNP/#!/snp?snp_id=rs1471371676&location=UTR3&five='1') |
| [rs1274440385](https://guolab.wchscu.cn/miRNASNP/#!/snp?snp_id=rs1274440385&location=UTR3&one=1) | chr2:190969584 | G/A | -/- | [1](https://guolab.wchscu.cn/miRNASNP/#!/snp?snp_id=rs1274440385&location=UTR3&four='1') | [27](https://guolab.wchscu.cn/miRNASNP/#!/snp?snp_id=rs1274440385&location=UTR3&five='1') |
| [rs1366670744](https://guolab.wchscu.cn/miRNASNP/#!/snp?snp_id=rs1366670744&location=UTR3&one=1) | chr2:190969600 | G/GTTTT | -/- | [7](https://guolab.wchscu.cn/miRNASNP/#!/snp?snp_id=rs1366670744&location=UTR3&four='1') | [3](https://guolab.wchscu.cn/miRNASNP/#!/snp?snp_id=rs1366670744&location=UTR3&five='1') |
| [rs1217310251](https://guolab.wchscu.cn/miRNASNP/#!/snp?snp_id=rs1217310251&location=UTR3&one=1) | chr2:190969601 | G/C | -/- | [1](https://guolab.wchscu.cn/miRNASNP/#!/snp?snp_id=rs1217310251&location=UTR3&four='1') | [2](https://guolab.wchscu.cn/miRNASNP/#!/snp?snp_id=rs1217310251&location=UTR3&five='1') |
| [rs1303132092](https://guolab.wchscu.cn/miRNASNP/#!/snp?snp_id=rs1303132092&location=UTR3&one=1) | chr2:190969602 | AAC/A | -/- | [6](https://guolab.wchscu.cn/miRNASNP/#!/snp?snp_id=rs1303132092&location=UTR3&four='1') | [2](https://guolab.wchscu.cn/miRNASNP/#!/snp?snp_id=rs1303132092&location=UTR3&five='1') |
| [rs550489659](https://guolab.wchscu.cn/miRNASNP/#!/snp?snp_id=rs550489659&location=UTR3&one=1) | chr2:190969609 | C/T | -/- | [6](https://guolab.wchscu.cn/miRNASNP/#!/snp?snp_id=rs550489659&location=UTR3&four='1') | 0 |
| [rs1197874166](https://guolab.wchscu.cn/miRNASNP/#!/snp?snp_id=rs1197874166&location=UTR3&one=1) | chr2:190969622 | AAC/A | -/- | [1](https://guolab.wchscu.cn/miRNASNP/#!/snp?snp_id=rs1197874166&location=UTR3&four='1') | [2](https://guolab.wchscu.cn/miRNASNP/#!/snp?snp_id=rs1197874166&location=UTR3&five='1') |
| [rs1363631252](https://guolab.wchscu.cn/miRNASNP/#!/snp?snp_id=rs1363631252&location=UTR3&one=1) | chr2:190969622 | A/G | 1/- | [6](https://guolab.wchscu.cn/miRNASNP/#!/snp?snp_id=rs1363631252&location=UTR3&four='1') | [12](https://guolab.wchscu.cn/miRNASNP/#!/snp?snp_id=rs1363631252&location=UTR3&five='1') |
| [rs1425145245](https://guolab.wchscu.cn/miRNASNP/#!/snp?snp_id=rs1425145245&location=UTR3&one=1) | chr2:190969462 | G/A | 1/- | [5](https://guolab.wchscu.cn/miRNASNP/#!/snp?snp_id=rs1425145245&location=UTR3&four='1') | [8](https://guolab.wchscu.cn/miRNASNP/#!/snp?snp_id=rs1425145245&location=UTR3&five='1') |
| [rs1248007093](https://guolab.wchscu.cn/miRNASNP/#!/snp?snp_id=rs1248007093&location=UTR3&one=1) | chr2:190969469 | A/G | 1/- | [4](https://guolab.wchscu.cn/miRNASNP/#!/snp?snp_id=rs1248007093&location=UTR3&four='1') | [6](https://guolab.wchscu.cn/miRNASNP/#!/snp?snp_id=rs1248007093&location=UTR3&five='1') |
| [rs1178663231](https://guolab.wchscu.cn/miRNASNP/#!/snp?snp_id=rs1178663231&location=UTR3&one=1) | chr2:190969481 | C/T | -/- | [3](https://guolab.wchscu.cn/miRNASNP/#!/snp?snp_id=rs1178663231&location=UTR3&four='1') | [3](https://guolab.wchscu.cn/miRNASNP/#!/snp?snp_id=rs1178663231&location=UTR3&five='1') |
| [rs1197681996](https://guolab.wchscu.cn/miRNASNP/#!/snp?snp_id=rs1197681996&location=UTR3&one=1) | chr2:190969487 | C/G | 1/- | [2](https://guolab.wchscu.cn/miRNASNP/#!/snp?snp_id=rs1197681996&location=UTR3&four='1') | [2](https://guolab.wchscu.cn/miRNASNP/#!/snp?snp_id=rs1197681996&location=UTR3&five='1') |
| [rs1457799429](https://guolab.wchscu.cn/miRNASNP/#!/snp?snp_id=rs1457799429&location=UTR3&one=1) | chr2:190969500 | T/A | 1/- | [1](https://guolab.wchscu.cn/miRNASNP/#!/snp?snp_id=rs1457799429&location=UTR3&four='1') | [6](https://guolab.wchscu.cn/miRNASNP/#!/snp?snp_id=rs1457799429&location=UTR3&five='1') |
| [rs1360292452](https://guolab.wchscu.cn/miRNASNP/#!/snp?snp_id=rs1360292452&location=UTR3&one=1) | chr2:190969512 | TAAGG/T | -/- | [5](https://guolab.wchscu.cn/miRNASNP/#!/snp?snp_id=rs1360292452&location=UTR3&four='1') | [4](https://guolab.wchscu.cn/miRNASNP/#!/snp?snp_id=rs1360292452&location=UTR3&five='1') |
| [rs1465557782](https://guolab.wchscu.cn/miRNASNP/#!/snp?snp_id=rs1465557782&location=UTR3&one=1) | chr2:190969515 | G/T | -/- | [2](https://guolab.wchscu.cn/miRNASNP/#!/snp?snp_id=rs1465557782&location=UTR3&four='1') | [4](https://guolab.wchscu.cn/miRNASNP/#!/snp?snp_id=rs1465557782&location=UTR3&five='1') |
| [rs1020366127](https://guolab.wchscu.cn/miRNASNP/#!/snp?snp_id=rs1020366127&location=UTR3&one=1) | chr2:190969519 | G/C | 0.9999/0.0001 | [1](https://guolab.wchscu.cn/miRNASNP/#!/snp?snp_id=rs1020366127&location=UTR3&four='1') | [9](https://guolab.wchscu.cn/miRNASNP/#!/snp?snp_id=rs1020366127&location=UTR3&five='1') |
| [rs967975098](https://guolab.wchscu.cn/miRNASNP/#!/snp?snp_id=rs967975098&location=UTR3&one=1) | chr2:190969528 | T/A | 1/- | 0 | [6](https://guolab.wchscu.cn/miRNASNP/#!/snp?snp_id=rs967975098&location=UTR3&five='1') |
| [rs869300110](https://guolab.wchscu.cn/miRNASNP/#!/snp?snp_id=rs869300110&location=UTR3&one=1) | chr2:190969534 | T/TA | -/- | [3](https://guolab.wchscu.cn/miRNASNP/#!/snp?snp_id=rs869300110&location=UTR3&four='1') | [1](https://guolab.wchscu.cn/miRNASNP/#!/snp?snp_id=rs869300110&location=UTR3&five='1') |
| [rs1334084021](https://guolab.wchscu.cn/miRNASNP/#!/snp?snp_id=rs1334084021&location=UTR3&one=1) | chr2:190969539 | T/C | 1/- | [5](https://guolab.wchscu.cn/miRNASNP/#!/snp?snp_id=rs1334084021&location=UTR3&four='1') | [4](https://guolab.wchscu.cn/miRNASNP/#!/snp?snp_id=rs1334084021&location=UTR3&five='1') |
| [rs1329457174](https://guolab.wchscu.cn/miRNASNP/#!/snp?snp_id=rs1329457174&location=UTR3&one=1) | chr2:190969540 | A/G | 1/- | [4](https://guolab.wchscu.cn/miRNASNP/#!/snp?snp_id=rs1329457174&location=UTR3&four='1') | [3](https://guolab.wchscu.cn/miRNASNP/#!/snp?snp_id=rs1329457174&location=UTR3&five='1') |
| [rs1000357522](https://guolab.wchscu.cn/miRNASNP/#!/snp?snp_id=rs1000357522&location=UTR3&one=1) | chr2:190969543 | C/T | 1/- | 0 | [4](https://guolab.wchscu.cn/miRNASNP/#!/snp?snp_id=rs1000357522&location=UTR3&five='1') |
| [rs946544691](https://guolab.wchscu.cn/miRNASNP/#!/snp?snp_id=rs946544691&location=UTR3&one=1) | chr2:190969544 | T/G | -/- | [3](https://guolab.wchscu.cn/miRNASNP/#!/snp?snp_id=rs946544691&location=UTR3&four='1') | [4](https://guolab.wchscu.cn/miRNASNP/#!/snp?snp_id=rs946544691&location=UTR3&five='1') |
| [rs1402477351](https://guolab.wchscu.cn/miRNASNP/#!/snp?snp_id=rs1402477351&location=UTR3&one=1) | chr2:190969548 | C/T | 1/- | [3](https://guolab.wchscu.cn/miRNASNP/#!/snp?snp_id=rs1402477351&location=UTR3&four='1') | [3](https://guolab.wchscu.cn/miRNASNP/#!/snp?snp_id=rs1402477351&location=UTR3&five='1') |
| [rs1159444997](https://guolab.wchscu.cn/miRNASNP/#!/snp?snp_id=rs1159444997&location=UTR3&one=1) | chr2:190969624 | C/A | -/- | [1](https://guolab.wchscu.cn/miRNASNP/#!/snp?snp_id=rs1159444997&location=UTR3&four='1') | [2](https://guolab.wchscu.cn/miRNASNP/#!/snp?snp_id=rs1159444997&location=UTR3&five='1') |
| [rs923863441](https://guolab.wchscu.cn/miRNASNP/#!/snp?snp_id=rs923863441&location=UTR3&one=1) | chr2:190969628 | T/G | 0.9998/0.0002 | [3](https://guolab.wchscu.cn/miRNASNP/#!/snp?snp_id=rs923863441&location=UTR3&four='1') | [4](https://guolab.wchscu.cn/miRNASNP/#!/snp?snp_id=rs923863441&location=UTR3&five='1') |
| [rs1458685708](https://guolab.wchscu.cn/miRNASNP/#!/snp?snp_id=rs1458685708&location=UTR3&one=1) | chr2:190969629 | A/G | 1/- | [1](https://guolab.wchscu.cn/miRNASNP/#!/snp?snp_id=rs1458685708&location=UTR3&four='1') | [3](https://guolab.wchscu.cn/miRNASNP/#!/snp?snp_id=rs1458685708&location=UTR3&five='1') |
| [rs944632360](https://guolab.wchscu.cn/miRNASNP/#!/snp?snp_id=rs944632360&location=UTR3&one=1) | chr2:190969633 | C/T | -/- | [1](https://guolab.wchscu.cn/miRNASNP/#!/snp?snp_id=rs944632360&location=UTR3&four='1') | [3](https://guolab.wchscu.cn/miRNASNP/#!/snp?snp_id=rs944632360&location=UTR3&five='1') |
| [rs1239405513](https://guolab.wchscu.cn/miRNASNP/#!/snp?snp_id=rs1239405513&location=UTR3&one=1) | chr2:190969634 | A/G | 1/- | [2](https://guolab.wchscu.cn/miRNASNP/#!/snp?snp_id=rs1239405513&location=UTR3&four='1') | [3](https://guolab.wchscu.cn/miRNASNP/#!/snp?snp_id=rs1239405513&location=UTR3&five='1') |
| [rs1191208553](https://guolab.wchscu.cn/miRNASNP/#!/snp?snp_id=rs1191208553&location=UTR3&one=1) | chr2:190969635 | G/A | 1/- | [1](https://guolab.wchscu.cn/miRNASNP/#!/snp?snp_id=rs1191208553&location=UTR3&four='1') | [2](https://guolab.wchscu.cn/miRNASNP/#!/snp?snp_id=rs1191208553&location=UTR3&five='1') |
| [rs916880590](https://guolab.wchscu.cn/miRNASNP/#!/snp?snp_id=rs916880590&location=UTR3&one=1) | chr2:190969636 | C/T | 1/- | [4](https://guolab.wchscu.cn/miRNASNP/#!/snp?snp_id=rs916880590&location=UTR3&four='1') | [1](https://guolab.wchscu.cn/miRNASNP/#!/snp?snp_id=rs916880590&location=UTR3&five='1') |
| [rs780655672](https://guolab.wchscu.cn/miRNASNP/#!/snp?snp_id=rs780655672&location=UTR3&one=1) | chr2:190969637 | G/A | 1/- | [9](https://guolab.wchscu.cn/miRNASNP/#!/snp?snp_id=rs780655672&location=UTR3&four='1') | [1](https://guolab.wchscu.cn/miRNASNP/#!/snp?snp_id=rs780655672&location=UTR3&five='1') |
| [rs1487284348](https://guolab.wchscu.cn/miRNASNP/#!/snp?snp_id=rs1487284348&location=UTR3&one=1) | chr2:190969641 | CAT/C | -/- | [1](https://guolab.wchscu.cn/miRNASNP/#!/snp?snp_id=rs1487284348&location=UTR3&four='1') | [1](https://guolab.wchscu.cn/miRNASNP/#!/snp?snp_id=rs1487284348&location=UTR3&five='1') |
| [rs900160511](https://guolab.wchscu.cn/miRNASNP/#!/snp?snp_id=rs900160511&location=UTR3&one=1) | chr2:190969642 | A/G | -/- | 0 | [2](https://guolab.wchscu.cn/miRNASNP/#!/snp?snp_id=rs900160511&location=UTR3&five='1') |
| [rs570854463](https://guolab.wchscu.cn/miRNASNP/#!/snp?snp_id=rs570854463&location=UTR3&one=1) | chr2:190969643 | T/C | -/- | [1](https://guolab.wchscu.cn/miRNASNP/#!/snp?snp_id=rs570854463&location=UTR3&four='1') | [1](https://guolab.wchscu.cn/miRNASNP/#!/snp?snp_id=rs570854463&location=UTR3&five='1') |
| [rs570854463](https://guolab.wchscu.cn/miRNASNP/#!/snp?snp_id=rs570854463&location=UTR3&one=1) | chr2:190969643 | T/G | -/- | [2](https://guolab.wchscu.cn/miRNASNP/#!/snp?snp_id=rs570854463&location=UTR3&four='1') | [1](https://guolab.wchscu.cn/miRNASNP/#!/snp?snp_id=rs570854463&location=UTR3&five='1') |
| [rs971028087](https://guolab.wchscu.cn/miRNASNP/#!/snp?snp_id=rs971028087&location=UTR3&one=1) | chr2:190969649 | G/A | 1/- | [2](https://guolab.wchscu.cn/miRNASNP/#!/snp?snp_id=rs971028087&location=UTR3&four='1') | [7](https://guolab.wchscu.cn/miRNASNP/#!/snp?snp_id=rs971028087&location=UTR3&five='1') |
| [rs982975198](https://guolab.wchscu.cn/miRNASNP/#!/snp?snp_id=rs982975198&location=UTR3&one=1) | chr2:190969650 | T/C | 1/- | [6](https://guolab.wchscu.cn/miRNASNP/#!/snp?snp_id=rs982975198&location=UTR3&four='1') | [7](https://guolab.wchscu.cn/miRNASNP/#!/snp?snp_id=rs982975198&location=UTR3&five='1') |
| [rs1259996266](https://guolab.wchscu.cn/miRNASNP/#!/snp?snp_id=rs1259996266&location=UTR3&one=1) | chr2:190969655 | A/G | 1/- | [1](https://guolab.wchscu.cn/miRNASNP/#!/snp?snp_id=rs1259996266&location=UTR3&four='1') | [3](https://guolab.wchscu.cn/miRNASNP/#!/snp?snp_id=rs1259996266&location=UTR3&five='1') |
| [rs1241131501](https://guolab.wchscu.cn/miRNASNP/#!/snp?snp_id=rs1241131501&location=UTR3&one=1) | chr2:190969666 | A/G | 1/- | [2](https://guolab.wchscu.cn/miRNASNP/#!/snp?snp_id=rs1241131501&location=UTR3&four='1') | [1](https://guolab.wchscu.cn/miRNASNP/#!/snp?snp_id=rs1241131501&location=UTR3&five='1') |
| [rs528626040](https://guolab.wchscu.cn/miRNASNP/#!/snp?snp_id=rs528626040&location=UTR3&one=1) | chr2:190969667 | T/C | 0.9997/0.0003 | [6](https://guolab.wchscu.cn/miRNASNP/#!/snp?snp_id=rs528626040&location=UTR3&four='1') | [1](https://guolab.wchscu.cn/miRNASNP/#!/snp?snp_id=rs528626040&location=UTR3&five='1') |
| [rs887503321](https://guolab.wchscu.cn/miRNASNP/#!/snp?snp_id=rs887503321&location=UTR3&one=1) | chr2:190969668 | A/G | -/- | [8](https://guolab.wchscu.cn/miRNASNP/#!/snp?snp_id=rs887503321&location=UTR3&four='1') | [3](https://guolab.wchscu.cn/miRNASNP/#!/snp?snp_id=rs887503321&location=UTR3&five='1') |
| [rs1005462562](https://guolab.wchscu.cn/miRNASNP/#!/snp?snp_id=rs1005462562&location=UTR3&one=1) | chr2:190969670 | G/C | 1/- | [5](https://guolab.wchscu.cn/miRNASNP/#!/snp?snp_id=rs1005462562&location=UTR3&four='1') | [3](https://guolab.wchscu.cn/miRNASNP/#!/snp?snp_id=rs1005462562&location=UTR3&five='1') |
| [rs547101504](https://guolab.wchscu.cn/miRNASNP/#!/snp?snp_id=rs547101504&location=UTR3&one=1) | chr2:190969672 | A/C | -/- | [2](https://guolab.wchscu.cn/miRNASNP/#!/snp?snp_id=rs547101504&location=UTR3&four='1') | [5](https://guolab.wchscu.cn/miRNASNP/#!/snp?snp_id=rs547101504&location=UTR3&five='1') |
| [rs1036496798](https://guolab.wchscu.cn/miRNASNP/#!/snp?snp_id=rs1036496798&location=UTR3&one=1) | chr2:190969675 | G/T | -/- | 0 | [2](https://guolab.wchscu.cn/miRNASNP/#!/snp?snp_id=rs1036496798&location=UTR3&five='1') |
| [rs907480414](https://guolab.wchscu.cn/miRNASNP/#!/snp?snp_id=rs907480414&location=UTR3&one=1) | chr2:190969677 | G/T | -/- | [3](https://guolab.wchscu.cn/miRNASNP/#!/snp?snp_id=rs907480414&location=UTR3&four='1') | [4](https://guolab.wchscu.cn/miRNASNP/#!/snp?snp_id=rs907480414&location=UTR3&five='1') |
| [rs1003123973](https://guolab.wchscu.cn/miRNASNP/#!/snp?snp_id=rs1003123973&location=UTR3&one=1) | chr2:190969678 | T/A | -/- | [2](https://guolab.wchscu.cn/miRNASNP/#!/snp?snp_id=rs1003123973&location=UTR3&four='1') | [4](https://guolab.wchscu.cn/miRNASNP/#!/snp?snp_id=rs1003123973&location=UTR3&five='1') |
| [rs941218552](https://guolab.wchscu.cn/miRNASNP/#!/snp?snp_id=rs941218552&location=UTR3&one=1) | chr2:190969683 | A/G | 1/- | [5](https://guolab.wchscu.cn/miRNASNP/#!/snp?snp_id=rs941218552&location=UTR3&four='1') | [2](https://guolab.wchscu.cn/miRNASNP/#!/snp?snp_id=rs941218552&location=UTR3&five='1') |
| [rs1425906541](https://guolab.wchscu.cn/miRNASNP/#!/snp?snp_id=rs1425906541&location=UTR3&one=1) | chr2:190969687 | G/A | -/- | [1](https://guolab.wchscu.cn/miRNASNP/#!/snp?snp_id=rs1425906541&location=UTR3&four='1') | [1](https://guolab.wchscu.cn/miRNASNP/#!/snp?snp_id=rs1425906541&location=UTR3&five='1') |
| [rs1331803615](https://guolab.wchscu.cn/miRNASNP/#!/snp?snp_id=rs1331803615&location=UTR3&one=1) | chr2:190969693 | A/G | -/- | [8](https://guolab.wchscu.cn/miRNASNP/#!/snp?snp_id=rs1331803615&location=UTR3&four='1') | [4](https://guolab.wchscu.cn/miRNASNP/#!/snp?snp_id=rs1331803615&location=UTR3&five='1') |
| [rs1038185021](https://guolab.wchscu.cn/miRNASNP/#!/snp?snp_id=rs1038185021&location=UTR3&one=1) | chr2:190969696 | C/T | 1/- | [1](https://guolab.wchscu.cn/miRNASNP/#!/snp?snp_id=rs1038185021&location=UTR3&four='1') | 0 |
| [rs1034372654](https://guolab.wchscu.cn/miRNASNP/#!/snp?snp_id=rs1034372654&location=UTR3&one=1) | chr2:190969699 | A/G | 1/- | 0 | [3](https://guolab.wchscu.cn/miRNASNP/#!/snp?snp_id=rs1034372654&location=UTR3&five='1') |
| [rs1384687037](https://guolab.wchscu.cn/miRNASNP/#!/snp?snp_id=rs1384687037&location=UTR3&one=1) | chr2:190969700 | T/C | 1/- | [7](https://guolab.wchscu.cn/miRNASNP/#!/snp?snp_id=rs1384687037&location=UTR3&four='1') | [4](https://guolab.wchscu.cn/miRNASNP/#!/snp?snp_id=rs1384687037&location=UTR3&five='1') |
| [rs562081130](https://guolab.wchscu.cn/miRNASNP/#!/snp?snp_id=rs562081130&location=UTR3&one=1) | chr2:190969701 | A/G | 0.9999/0.0001 | [1](https://guolab.wchscu.cn/miRNASNP/#!/snp?snp_id=rs562081130&location=UTR3&four='1') | [4](https://guolab.wchscu.cn/miRNASNP/#!/snp?snp_id=rs562081130&location=UTR3&five='1') |
| [rs769403390](https://guolab.wchscu.cn/miRNASNP/#!/snp?snp_id=rs769403390&location=UTR3&one=1) | chr2:190969711 | G/A | 0.9999/0.0001 | [7](https://guolab.wchscu.cn/miRNASNP/#!/snp?snp_id=rs769403390&location=UTR3&four='1') | [9](https://guolab.wchscu.cn/miRNASNP/#!/snp?snp_id=rs769403390&location=UTR3&five='1') |
| [rs1381808050](https://guolab.wchscu.cn/miRNASNP/#!/snp?snp_id=rs1381808050&location=UTR3&one=1) | chr2:190969715 | A/G | 1/- | [3](https://guolab.wchscu.cn/miRNASNP/#!/snp?snp_id=rs1381808050&location=UTR3&four='1') | [3](https://guolab.wchscu.cn/miRNASNP/#!/snp?snp_id=rs1381808050&location=UTR3&five='1') |
| [rs777682355](https://guolab.wchscu.cn/miRNASNP/#!/snp?snp_id=rs777682355&location=UTR3&one=1) | chr2:190969716 | T/C | -/- | [4](https://guolab.wchscu.cn/miRNASNP/#!/snp?snp_id=rs777682355&location=UTR3&four='1') | [3](https://guolab.wchscu.cn/miRNASNP/#!/snp?snp_id=rs777682355&location=UTR3&five='1') |
| [rs932524197](https://guolab.wchscu.cn/miRNASNP/#!/snp?snp_id=rs932524197&location=UTR3&one=1) | chr2:190969717 | T/A | 1/- | [4](https://guolab.wchscu.cn/miRNASNP/#!/snp?snp_id=rs932524197&location=UTR3&four='1') | [3](https://guolab.wchscu.cn/miRNASNP/#!/snp?snp_id=rs932524197&location=UTR3&five='1') |
| [rs749141524](https://guolab.wchscu.cn/miRNASNP/#!/snp?snp_id=rs749141524&location=UTR3&one=1) | chr2:190969719 | T/A | -/- | [1](https://guolab.wchscu.cn/miRNASNP/#!/snp?snp_id=rs749141524&location=UTR3&four='1') | [4](https://guolab.wchscu.cn/miRNASNP/#!/snp?snp_id=rs749141524&location=UTR3&five='1') |
| [rs1477395561](https://guolab.wchscu.cn/miRNASNP/#!/snp?snp_id=rs1477395561&location=UTR3&one=1) | chr2:190969723 | A/G | 1/- | [6](https://guolab.wchscu.cn/miRNASNP/#!/snp?snp_id=rs1477395561&location=UTR3&four='1') | [8](https://guolab.wchscu.cn/miRNASNP/#!/snp?snp_id=rs1477395561&location=UTR3&five='1') |
| [rs1246824597](https://guolab.wchscu.cn/miRNASNP/#!/snp?snp_id=rs1246824597&location=UTR3&one=1) | chr2:190969724 | G/A | 1/- | [1](https://guolab.wchscu.cn/miRNASNP/#!/snp?snp_id=rs1246824597&location=UTR3&four='1') | [8](https://guolab.wchscu.cn/miRNASNP/#!/snp?snp_id=rs1246824597&location=UTR3&five='1') |
| [rs1200907242](https://guolab.wchscu.cn/miRNASNP/#!/snp?snp_id=rs1200907242&location=UTR3&one=1) | chr2:190969733 | T/C | 1/- | [7](https://guolab.wchscu.cn/miRNASNP/#!/snp?snp_id=rs1200907242&location=UTR3&four='1') | [1](https://guolab.wchscu.cn/miRNASNP/#!/snp?snp_id=rs1200907242&location=UTR3&five='1') |
| [rs1021720997](https://guolab.wchscu.cn/miRNASNP/#!/snp?snp_id=rs1021720997&location=UTR3&one=1) | chr2:190969736 | T/C | -/- | [1](https://guolab.wchscu.cn/miRNASNP/#!/snp?snp_id=rs1021720997&location=UTR3&four='1') | 0 |
| [rs967454724](https://guolab.wchscu.cn/miRNASNP/#!/snp?snp_id=rs967454724&location=UTR3&one=1) | chr2:190969744 | C/T | 0.9999/0.0001 | [1](https://guolab.wchscu.cn/miRNASNP/#!/snp?snp_id=rs967454724&location=UTR3&four='1') | 0 |
| [rs41324145](https://guolab.wchscu.cn/miRNASNP/#!/snp?snp_id=rs41324145&location=UTR3&one=1) | chr2:190969750 | C/A | 0.9944/0.0056 | [4](https://guolab.wchscu.cn/miRNASNP/#!/snp?snp_id=rs41324145&location=UTR3&four='1') | [1](https://guolab.wchscu.cn/miRNASNP/#!/snp?snp_id=rs41324145&location=UTR3&five='1') |
| [rs184180073](https://guolab.wchscu.cn/miRNASNP/#!/snp?snp_id=rs184180073&location=UTR3&one=1) | chr2:190969751 | T/C | 1/- | [10](https://guolab.wchscu.cn/miRNASNP/#!/snp?snp_id=rs184180073&location=UTR3&four='1') | [2](https://guolab.wchscu.cn/miRNASNP/#!/snp?snp_id=rs184180073&location=UTR3&five='1') |
| [rs1356062539](https://guolab.wchscu.cn/miRNASNP/#!/snp?snp_id=rs1356062539&location=UTR3&one=1) | chr2:190969752 | T/G | 1/- | [6](https://guolab.wchscu.cn/miRNASNP/#!/snp?snp_id=rs1356062539&location=UTR3&four='1') | [2](https://guolab.wchscu.cn/miRNASNP/#!/snp?snp_id=rs1356062539&location=UTR3&five='1') |
| [rs569019701](https://guolab.wchscu.cn/miRNASNP/#!/snp?snp_id=rs569019701&location=UTR3&one=1) | chr2:190969754 | C/T | 1/- | [1](https://guolab.wchscu.cn/miRNASNP/#!/snp?snp_id=rs569019701&location=UTR3&four='1') | [2](https://guolab.wchscu.cn/miRNASNP/#!/snp?snp_id=rs569019701&location=UTR3&five='1') |
| [rs976014229](https://guolab.wchscu.cn/miRNASNP/#!/snp?snp_id=rs976014229&location=UTR3&one=1) | chr2:190969759 | T/C | -/- | 0 | [1](https://guolab.wchscu.cn/miRNASNP/#!/snp?snp_id=rs976014229&location=UTR3&five='1') |
| [rs41363648](https://guolab.wchscu.cn/miRNASNP/#!/snp?snp_id=rs41363648&location=UTR3&one=1) | chr2:190969761 | T/C | 0.9828/0.0173 | [6](https://guolab.wchscu.cn/miRNASNP/#!/snp?snp_id=rs41363648&location=UTR3&four='1') | [3](https://guolab.wchscu.cn/miRNASNP/#!/snp?snp_id=rs41363648&location=UTR3&five='1') |
| [rs551312847](https://guolab.wchscu.cn/miRNASNP/#!/snp?snp_id=rs551312847&location=UTR3&one=1) | chr2:190969763 | G/A | -/- | [4](https://guolab.wchscu.cn/miRNASNP/#!/snp?snp_id=rs551312847&location=UTR3&four='1') | [4](https://guolab.wchscu.cn/miRNASNP/#!/snp?snp_id=rs551312847&location=UTR3&five='1') |
| [rs1345840617](https://guolab.wchscu.cn/miRNASNP/#!/snp?snp_id=rs1345840617&location=UTR3&one=1) | chr2:190969769 | A/G | -/- | [3](https://guolab.wchscu.cn/miRNASNP/#!/snp?snp_id=rs1345840617&location=UTR3&four='1') | [1](https://guolab.wchscu.cn/miRNASNP/#!/snp?snp_id=rs1345840617&location=UTR3&five='1') |
| [rs1276749157](https://guolab.wchscu.cn/miRNASNP/#!/snp?snp_id=rs1276749157&location=UTR3&one=1) | chr2:190969771 | T/TA | -/- | [5](https://guolab.wchscu.cn/miRNASNP/#!/snp?snp_id=rs1276749157&location=UTR3&four='1') | [2](https://guolab.wchscu.cn/miRNASNP/#!/snp?snp_id=rs1276749157&location=UTR3&five='1') |
| [rs886055375](https://guolab.wchscu.cn/miRNASNP/#!/snp?snp_id=rs886055375&location=UTR3&one=1) | chr2:190969771 | T/C | 1/- | [2](https://guolab.wchscu.cn/miRNASNP/#!/snp?snp_id=rs886055375&location=UTR3&four='1') | [2](https://guolab.wchscu.cn/miRNASNP/#!/snp?snp_id=rs886055375&location=UTR3&five='1') |
| [rs886055375](https://guolab.wchscu.cn/miRNASNP/#!/snp?snp_id=rs886055375&location=UTR3&one=1) | chr2:190969771 | T/G | 1/- | [1](https://guolab.wchscu.cn/miRNASNP/#!/snp?snp_id=rs886055375&location=UTR3&four='1') | [2](https://guolab.wchscu.cn/miRNASNP/#!/snp?snp_id=rs886055375&location=UTR3&five='1') |
| [rs772033315](https://guolab.wchscu.cn/miRNASNP/#!/snp?snp_id=rs772033315&location=UTR3&one=1) | chr2:190969775 | CTAAA/C | -/0.0001 | [1](https://guolab.wchscu.cn/miRNASNP/#!/snp?snp_id=rs772033315&location=UTR3&four='1') | [2](https://guolab.wchscu.cn/miRNASNP/#!/snp?snp_id=rs772033315&location=UTR3&five='1') |
| [rs886055376](https://guolab.wchscu.cn/miRNASNP/#!/snp?snp_id=rs886055376&location=UTR3&one=1) | chr2:190969782 | C/G | -/- | [6](https://guolab.wchscu.cn/miRNASNP/#!/snp?snp_id=rs886055376&location=UTR3&four='1') | 0 |
| [rs1326360631](https://guolab.wchscu.cn/miRNASNP/#!/snp?snp_id=rs1326360631&location=UTR3&one=1) | chr2:190969801 | G/A | 1/- | [4](https://guolab.wchscu.cn/miRNASNP/#!/snp?snp_id=rs1326360631&location=UTR3&four='1') | [3](https://guolab.wchscu.cn/miRNASNP/#!/snp?snp_id=rs1326360631&location=UTR3&five='1') |
| [rs566313103](https://guolab.wchscu.cn/miRNASNP/#!/snp?snp_id=rs566313103&location=UTR3&one=1) | chr2:190969824 | C/A | -/- | [3](https://guolab.wchscu.cn/miRNASNP/#!/snp?snp_id=rs566313103&location=UTR3&four='1') | [2](https://guolab.wchscu.cn/miRNASNP/#!/snp?snp_id=rs566313103&location=UTR3&five='1') |
| [rs1299140983](https://guolab.wchscu.cn/miRNASNP/#!/snp?snp_id=rs1299140983&location=UTR3&one=1) | chr2:190969827 | G/T | 1/- | [2](https://guolab.wchscu.cn/miRNASNP/#!/snp?snp_id=rs1299140983&location=UTR3&four='1') | [3](https://guolab.wchscu.cn/miRNASNP/#!/snp?snp_id=rs1299140983&location=UTR3&five='1') |
| [rs1048669825](https://guolab.wchscu.cn/miRNASNP/#!/snp?snp_id=rs1048669825&location=UTR3&one=1) | chr2:190969829 | A/G | -/- | [1](https://guolab.wchscu.cn/miRNASNP/#!/snp?snp_id=rs1048669825&location=UTR3&four='1') | [3](https://guolab.wchscu.cn/miRNASNP/#!/snp?snp_id=rs1048669825&location=UTR3&five='1') |
| [rs1374073561](https://guolab.wchscu.cn/miRNASNP/#!/snp?snp_id=rs1374073561&location=UTR3&one=1) | chr2:190969836 | T/G | 1/- | [5](https://guolab.wchscu.cn/miRNASNP/#!/snp?snp_id=rs1374073561&location=UTR3&four='1') | [4](https://guolab.wchscu.cn/miRNASNP/#!/snp?snp_id=rs1374073561&location=UTR3&five='1') |
| [rs908972162](https://guolab.wchscu.cn/miRNASNP/#!/snp?snp_id=rs908972162&location=UTR3&one=1) | chr2:190969839 | T/C | -/- | 0 | [4](https://guolab.wchscu.cn/miRNASNP/#!/snp?snp_id=rs908972162&location=UTR3&five='1') |
| [rs1171326918](https://guolab.wchscu.cn/miRNASNP/#!/snp?snp_id=rs1171326918&location=UTR3&one=1) | chr2:190969840 | G/C | 1/- | [3](https://guolab.wchscu.cn/miRNASNP/#!/snp?snp_id=rs1171326918&location=UTR3&four='1') | [4](https://guolab.wchscu.cn/miRNASNP/#!/snp?snp_id=rs1171326918&location=UTR3&five='1') |
| [rs1413622801](https://guolab.wchscu.cn/miRNASNP/#!/snp?snp_id=rs1413622801&location=UTR3&one=1) | chr2:190969845 | T/C | 1/- | [4](https://guolab.wchscu.cn/miRNASNP/#!/snp?snp_id=rs1413622801&location=UTR3&four='1') | 0 |
| [rs1425238981](https://guolab.wchscu.cn/miRNASNP/#!/snp?snp_id=rs1425238981&location=UTR3&one=1) | chr2:190969847 | T/C | 1/- | 0 | 0 |
| [rs188557905](https://guolab.wchscu.cn/miRNASNP/#!/snp?snp_id=rs188557905&location=UTR3&one=1) | chr2:190969848 | C/T | 0.9995/0.0005 | [1](https://guolab.wchscu.cn/miRNASNP/#!/snp?snp_id=rs188557905&location=UTR3&four='1') | 0 |
| [rs1214688680](https://guolab.wchscu.cn/miRNASNP/#!/snp?snp_id=rs1214688680&location=UTR3&one=1) | chr2:190969852 | A/G | -/- | [1](https://guolab.wchscu.cn/miRNASNP/#!/snp?snp_id=rs1214688680&location=UTR3&four='1') | 0 |
| [rs1474363657](https://guolab.wchscu.cn/miRNASNP/#!/snp?snp_id=rs1474363657&location=UTR3&one=1) | chr2:190969856 | A/G | 1/- | [3](https://guolab.wchscu.cn/miRNASNP/#!/snp?snp_id=rs1474363657&location=UTR3&four='1') | [3](https://guolab.wchscu.cn/miRNASNP/#!/snp?snp_id=rs1474363657&location=UTR3&five='1') |
| [rs1288525683](https://guolab.wchscu.cn/miRNASNP/#!/snp?snp_id=rs1288525683&location=UTR3&one=1) | chr2:190969873 | T/C | -/- | [5](https://guolab.wchscu.cn/miRNASNP/#!/snp?snp_id=rs1288525683&location=UTR3&four='1') | [1](https://guolab.wchscu.cn/miRNASNP/#!/snp?snp_id=rs1288525683&location=UTR3&five='1') |
| [rs1422860396](https://guolab.wchscu.cn/miRNASNP/#!/snp?snp_id=rs1422860396&location=UTR3&one=1) | chr2:190969875 | G/C | 1/- | [2](https://guolab.wchscu.cn/miRNASNP/#!/snp?snp_id=rs1422860396&location=UTR3&four='1') | 0 |
| [rs1194364922](https://guolab.wchscu.cn/miRNASNP/#!/snp?snp_id=rs1194364922&location=UTR3&one=1) | chr2:190969878 | T/C | 1/- | [1](https://guolab.wchscu.cn/miRNASNP/#!/snp?snp_id=rs1194364922&location=UTR3&four='1') | [4](https://guolab.wchscu.cn/miRNASNP/#!/snp?snp_id=rs1194364922&location=UTR3&five='1') |
| [rs903239098](https://guolab.wchscu.cn/miRNASNP/#!/snp?snp_id=rs903239098&location=UTR3&one=1) | chr2:190969882 | T/C | -/- | [2](https://guolab.wchscu.cn/miRNASNP/#!/snp?snp_id=rs903239098&location=UTR3&four='1') | [10](https://guolab.wchscu.cn/miRNASNP/#!/snp?snp_id=rs903239098&location=UTR3&five='1') |
| [rs1036975160](https://guolab.wchscu.cn/miRNASNP/#!/snp?snp_id=rs1036975160&location=UTR3&one=1) | chr2:190969884 | G/A | 1/- | [10](https://guolab.wchscu.cn/miRNASNP/#!/snp?snp_id=rs1036975160&location=UTR3&four='1') | [9](https://guolab.wchscu.cn/miRNASNP/#!/snp?snp_id=rs1036975160&location=UTR3&five='1') |
| [rs1319612138](https://guolab.wchscu.cn/miRNASNP/#!/snp?snp_id=rs1319612138&location=UTR3&one=1) | chr2:190969889 | GA/G | -/- | [5](https://guolab.wchscu.cn/miRNASNP/#!/snp?snp_id=rs1319612138&location=UTR3&four='1') | [3](https://guolab.wchscu.cn/miRNASNP/#!/snp?snp_id=rs1319612138&location=UTR3&five='1') |
| [rs907349445](https://guolab.wchscu.cn/miRNASNP/#!/snp?snp_id=rs907349445&location=UTR3&one=1) | chr2:190969889 | G/T | 1/- | [3](https://guolab.wchscu.cn/miRNASNP/#!/snp?snp_id=rs907349445&location=UTR3&four='1') | [4](https://guolab.wchscu.cn/miRNASNP/#!/snp?snp_id=rs907349445&location=UTR3&five='1') |
| [rs1033141232](https://guolab.wchscu.cn/miRNASNP/#!/snp?snp_id=rs1033141232&location=UTR3&one=1) | chr2:190969894 | G/A | 1/- | 0 | [1](https://guolab.wchscu.cn/miRNASNP/#!/snp?snp_id=rs1033141232&location=UTR3&five='1') |
| [rs958955361](https://guolab.wchscu.cn/miRNASNP/#!/snp?snp_id=rs958955361&location=UTR3&one=1) | chr2:190969896 | C/G | 1/- | 0 | [1](https://guolab.wchscu.cn/miRNASNP/#!/snp?snp_id=rs958955361&location=UTR3&five='1') |
| [rs1013154846](https://guolab.wchscu.cn/miRNASNP/#!/snp?snp_id=rs1013154846&location=UTR3&one=1) | chr2:190969898 | A/G | 1/- | [3](https://guolab.wchscu.cn/miRNASNP/#!/snp?snp_id=rs1013154846&location=UTR3&four='1') | [1](https://guolab.wchscu.cn/miRNASNP/#!/snp?snp_id=rs1013154846&location=UTR3&five='1') |
| [rs1024508098](https://guolab.wchscu.cn/miRNASNP/#!/snp?snp_id=rs1024508098&location=UTR3&one=1) | chr2:190969899 | A/C | 1/- | [3](https://guolab.wchscu.cn/miRNASNP/#!/snp?snp_id=rs1024508098&location=UTR3&four='1') | [1](https://guolab.wchscu.cn/miRNASNP/#!/snp?snp_id=rs1024508098&location=UTR3&five='1') |
| [rs1455797633](https://guolab.wchscu.cn/miRNASNP/#!/snp?snp_id=rs1455797633&location=UTR3&one=1) | chr2:190969900 | C/T | -/- | [4](https://guolab.wchscu.cn/miRNASNP/#!/snp?snp_id=rs1455797633&location=UTR3&four='1') | [5](https://guolab.wchscu.cn/miRNASNP/#!/snp?snp_id=rs1455797633&location=UTR3&five='1') |
| [rs1270055315](https://guolab.wchscu.cn/miRNASNP/#!/snp?snp_id=rs1270055315&location=UTR3&one=1) | chr2:190969906 | T/G | -/- | [21](https://guolab.wchscu.cn/miRNASNP/#!/snp?snp_id=rs1270055315&location=UTR3&four='1') | [24](https://guolab.wchscu.cn/miRNASNP/#!/snp?snp_id=rs1270055315&location=UTR3&five='1') |
| [rs1003564535](https://guolab.wchscu.cn/miRNASNP/#!/snp?snp_id=rs1003564535&location=UTR3&one=1) | chr2:190969916 | G/C | 1/- | [5](https://guolab.wchscu.cn/miRNASNP/#!/snp?snp_id=rs1003564535&location=UTR3&four='1') | [1](https://guolab.wchscu.cn/miRNASNP/#!/snp?snp_id=rs1003564535&location=UTR3&five='1') |
| [rs1221044628](https://guolab.wchscu.cn/miRNASNP/#!/snp?snp_id=rs1221044628&location=UTR3&one=1) | chr2:190969922 | A/G | 1/- | [30](https://guolab.wchscu.cn/miRNASNP/#!/snp?snp_id=rs1221044628&location=UTR3&four='1') | [2](https://guolab.wchscu.cn/miRNASNP/#!/snp?snp_id=rs1221044628&location=UTR3&five='1') |
| [rs180904823](https://guolab.wchscu.cn/miRNASNP/#!/snp?snp_id=rs180904823&location=UTR3&one=1) | chr2:190969926 | T/C | 0.9967/0.0033 | [1](https://guolab.wchscu.cn/miRNASNP/#!/snp?snp_id=rs180904823&location=UTR3&four='1') | [1](https://guolab.wchscu.cn/miRNASNP/#!/snp?snp_id=rs180904823&location=UTR3&five='1') |
| [rs982454382](https://guolab.wchscu.cn/miRNASNP/#!/snp?snp_id=rs982454382&location=UTR3&one=1) | chr2:190969931 | T/A | 1/- | 0 | [6](https://guolab.wchscu.cn/miRNASNP/#!/snp?snp_id=rs982454382&location=UTR3&five='1') |
| [rs982454382](https://guolab.wchscu.cn/miRNASNP/#!/snp?snp_id=rs982454382&location=UTR3&one=1) | chr2:190969931 | T/C | 1/- | 0 | [6](https://guolab.wchscu.cn/miRNASNP/#!/snp?snp_id=rs982454382&location=UTR3&five='1') |
| [rs1266031406](https://guolab.wchscu.cn/miRNASNP/#!/snp?snp_id=rs1266031406&location=UTR3&one=1) | chr2:190969934 | G/C | -/- | [2](https://guolab.wchscu.cn/miRNASNP/#!/snp?snp_id=rs1266031406&location=UTR3&four='1') | [3](https://guolab.wchscu.cn/miRNASNP/#!/snp?snp_id=rs1266031406&location=UTR3&five='1') |
| [rs1478415376](https://guolab.wchscu.cn/miRNASNP/#!/snp?snp_id=rs1478415376&location=UTR3&one=1) | chr2:190969937 | C/T | -/- | [5](https://guolab.wchscu.cn/miRNASNP/#!/snp?snp_id=rs1478415376&location=UTR3&four='1') | [1](https://guolab.wchscu.cn/miRNASNP/#!/snp?snp_id=rs1478415376&location=UTR3&five='1') |
| [rs908300239](https://guolab.wchscu.cn/miRNASNP/#!/snp?snp_id=rs908300239&location=UTR3&one=1) | chr2:190969943 | G/A | 1/- | [4](https://guolab.wchscu.cn/miRNASNP/#!/snp?snp_id=rs908300239&location=UTR3&four='1') | [4](https://guolab.wchscu.cn/miRNASNP/#!/snp?snp_id=rs908300239&location=UTR3&five='1') |
| [rs894706086](https://guolab.wchscu.cn/miRNASNP/#!/snp?snp_id=rs894706086&location=UTR3&one=1) | chr2:190969946 | A/G | 0.9995/0.0005 | [16](https://guolab.wchscu.cn/miRNASNP/#!/snp?snp_id=rs894706086&location=UTR3&four='1') | [3](https://guolab.wchscu.cn/miRNASNP/#!/snp?snp_id=rs894706086&location=UTR3&five='1') |
| [rs1398973007](https://guolab.wchscu.cn/miRNASNP/#!/snp?snp_id=rs1398973007&location=UTR3&one=1) | chr2:190969947 | G/A | 1/- | [1](https://guolab.wchscu.cn/miRNASNP/#!/snp?snp_id=rs1398973007&location=UTR3&four='1') | [4](https://guolab.wchscu.cn/miRNASNP/#!/snp?snp_id=rs1398973007&location=UTR3&five='1') |
| [rs1407303390](https://guolab.wchscu.cn/miRNASNP/#!/snp?snp_id=rs1407303390&location=UTR3&one=1) | chr2:190969961 | G/A | 0.9999/0.0001 | [1](https://guolab.wchscu.cn/miRNASNP/#!/snp?snp_id=rs1407303390&location=UTR3&four='1') | [5](https://guolab.wchscu.cn/miRNASNP/#!/snp?snp_id=rs1407303390&location=UTR3&five='1') |
| [rs1165708718](https://guolab.wchscu.cn/miRNASNP/#!/snp?snp_id=rs1165708718&location=UTR3&one=1) | chr2:190969981 | C/T | 1/- | [3](https://guolab.wchscu.cn/miRNASNP/#!/snp?snp_id=rs1165708718&location=UTR3&four='1') | [2](https://guolab.wchscu.cn/miRNASNP/#!/snp?snp_id=rs1165708718&location=UTR3&five='1') |
| [rs772206364](https://guolab.wchscu.cn/miRNASNP/#!/snp?snp_id=rs772206364&location=UTR3&one=1) | chr2:190969982 | T/G | 1/- | 0 | [2](https://guolab.wchscu.cn/miRNASNP/#!/snp?snp_id=rs772206364&location=UTR3&five='1') |
| [rs566925477](https://guolab.wchscu.cn/miRNASNP/#!/snp?snp_id=rs566925477&location=UTR3&one=1) | chr2:190969985 | T/G | 0.9999/0.0001 | [10](https://guolab.wchscu.cn/miRNASNP/#!/snp?snp_id=rs566925477&location=UTR3&four='1') | [6](https://guolab.wchscu.cn/miRNASNP/#!/snp?snp_id=rs566925477&location=UTR3&five='1') |
| [rs537402240](https://guolab.wchscu.cn/miRNASNP/#!/snp?snp_id=rs537402240&location=UTR3&one=1) | chr2:190969988 | A/G | -/- | [1](https://guolab.wchscu.cn/miRNASNP/#!/snp?snp_id=rs537402240&location=UTR3&four='1') | [5](https://guolab.wchscu.cn/miRNASNP/#!/snp?snp_id=rs537402240&location=UTR3&five='1') |
| [rs921134748](https://guolab.wchscu.cn/miRNASNP/#!/snp?snp_id=rs921134748&location=UTR3&one=1) | chr2:190969990 | T/C | 1/- | [8](https://guolab.wchscu.cn/miRNASNP/#!/snp?snp_id=rs921134748&location=UTR3&four='1') | [4](https://guolab.wchscu.cn/miRNASNP/#!/snp?snp_id=rs921134748&location=UTR3&five='1') |
| [rs1239449774](https://guolab.wchscu.cn/miRNASNP/#!/snp?snp_id=rs1239449774&location=UTR3&one=1) | chr2:190970003 | T/C | 1/- | [2](https://guolab.wchscu.cn/miRNASNP/#!/snp?snp_id=rs1239449774&location=UTR3&four='1') | [6](https://guolab.wchscu.cn/miRNASNP/#!/snp?snp_id=rs1239449774&location=UTR3&five='1') |
| [rs1365987295](https://guolab.wchscu.cn/miRNASNP/#!/snp?snp_id=rs1365987295&location=UTR3&one=1) | chr2:190970010 | A/G | -/- | [5](https://guolab.wchscu.cn/miRNASNP/#!/snp?snp_id=rs1365987295&location=UTR3&four='1') | [2](https://guolab.wchscu.cn/miRNASNP/#!/snp?snp_id=rs1365987295&location=UTR3&five='1') |
| [rs1011592854](https://guolab.wchscu.cn/miRNASNP/#!/snp?snp_id=rs1011592854&location=UTR3&one=1) | chr2:190970011 | T/C | 0.9999/- | 0 | [2](https://guolab.wchscu.cn/miRNASNP/#!/snp?snp_id=rs1011592854&location=UTR3&five='1') |
| [rs1011592854](https://guolab.wchscu.cn/miRNASNP/#!/snp?snp_id=rs1011592854&location=UTR3&one=1) | chr2:190970011 | T/G | 0.9999/0.0001 | [9](https://guolab.wchscu.cn/miRNASNP/#!/snp?snp_id=rs1011592854&location=UTR3&four='1') | [2](https://guolab.wchscu.cn/miRNASNP/#!/snp?snp_id=rs1011592854&location=UTR3&five='1') |
| [rs1051032536](https://guolab.wchscu.cn/miRNASNP/#!/snp?snp_id=rs1051032536&location=UTR3&one=1) | chr2:190970017 | T/A | 1/- | [6](https://guolab.wchscu.cn/miRNASNP/#!/snp?snp_id=rs1051032536&location=UTR3&four='1') | [2](https://guolab.wchscu.cn/miRNASNP/#!/snp?snp_id=rs1051032536&location=UTR3&five='1') |
| [rs1258870044](https://guolab.wchscu.cn/miRNASNP/#!/snp?snp_id=rs1258870044&location=UTR3&one=1) | chr2:190970018 | T/C | 1/- | [4](https://guolab.wchscu.cn/miRNASNP/#!/snp?snp_id=rs1258870044&location=UTR3&four='1') | [2](https://guolab.wchscu.cn/miRNASNP/#!/snp?snp_id=rs1258870044&location=UTR3&five='1') |
| [rs1201056639](https://guolab.wchscu.cn/miRNASNP/#!/snp?snp_id=rs1201056639&location=UTR3&one=1) | chr2:190970028 | C/A | 1/- | [2](https://guolab.wchscu.cn/miRNASNP/#!/snp?snp_id=rs1201056639&location=UTR3&four='1') | [2](https://guolab.wchscu.cn/miRNASNP/#!/snp?snp_id=rs1201056639&location=UTR3&five='1') |
| [rs186033487](https://guolab.wchscu.cn/miRNASNP/#!/snp?snp_id=rs186033487&location=UTR3&one=1) | chr2:190970033 | A/C | 0.9997/0.0003 | [1](https://guolab.wchscu.cn/miRNASNP/#!/snp?snp_id=rs186033487&location=UTR3&four='1') | [1](https://guolab.wchscu.cn/miRNASNP/#!/snp?snp_id=rs186033487&location=UTR3&five='1') |
| [rs575893899](https://guolab.wchscu.cn/miRNASNP/#!/snp?snp_id=rs575893899&location=UTR3&one=1) | chr2:190970038 | T/A | -/- | [1](https://guolab.wchscu.cn/miRNASNP/#!/snp?snp_id=rs575893899&location=UTR3&four='1') | 0 |
| [rs575893899](https://guolab.wchscu.cn/miRNASNP/#!/snp?snp_id=rs575893899&location=UTR3&one=1) | chr2:190970038 | T/C | -/- | [4](https://guolab.wchscu.cn/miRNASNP/#!/snp?snp_id=rs575893899&location=UTR3&four='1') | 0 |
| [rs45449693](https://guolab.wchscu.cn/miRNASNP/#!/snp?snp_id=rs45449693&location=UTR3&one=1) | chr2:190970042 | A/G | 0.9999/0.0001 | 0 | [4](https://guolab.wchscu.cn/miRNASNP/#!/snp?snp_id=rs45449693&location=UTR3&five='1') |
| [rs1041695115](https://guolab.wchscu.cn/miRNASNP/#!/snp?snp_id=rs1041695115&location=UTR3&one=1) | chr2:190970050 | AAGCTAAT/A | -/0.0001 | [7](https://guolab.wchscu.cn/miRNASNP/#!/snp?snp_id=rs1041695115&location=UTR3&four='1') | [7](https://guolab.wchscu.cn/miRNASNP/#!/snp?snp_id=rs1041695115&location=UTR3&five='1') |
| [rs1381858784](https://guolab.wchscu.cn/miRNASNP/#!/snp?snp_id=rs1381858784&location=UTR3&one=1) | chr2:190970055 | A/T | 1/- | [4](https://guolab.wchscu.cn/miRNASNP/#!/snp?snp_id=rs1381858784&location=UTR3&four='1') | [1](https://guolab.wchscu.cn/miRNASNP/#!/snp?snp_id=rs1381858784&location=UTR3&five='1') |
| [rs1285562768](https://guolab.wchscu.cn/miRNASNP/#!/snp?snp_id=rs1285562768&location=UTR3&one=1) | chr2:190970058 | A/T | 1/- | [3](https://guolab.wchscu.cn/miRNASNP/#!/snp?snp_id=rs1285562768&location=UTR3&four='1') | [2](https://guolab.wchscu.cn/miRNASNP/#!/snp?snp_id=rs1285562768&location=UTR3&five='1') |
| [rs56000300](https://guolab.wchscu.cn/miRNASNP/#!/snp?snp_id=rs56000300&location=UTR3&one=1) | chr2:190970058 | ATTCTC/A | -/- | [2](https://guolab.wchscu.cn/miRNASNP/#!/snp?snp_id=rs56000300&location=UTR3&four='1') | [6](https://guolab.wchscu.cn/miRNASNP/#!/snp?snp_id=rs56000300&location=UTR3&five='1') |
| [rs1398907439](https://guolab.wchscu.cn/miRNASNP/#!/snp?snp_id=rs1398907439&location=UTR3&one=1) | chr2:190970059 | T/G | 1/- | [4](https://guolab.wchscu.cn/miRNASNP/#!/snp?snp_id=rs1398907439&location=UTR3&four='1') | [5](https://guolab.wchscu.cn/miRNASNP/#!/snp?snp_id=rs1398907439&location=UTR3&five='1') |
| [rs1031282966](https://guolab.wchscu.cn/miRNASNP/#!/snp?snp_id=rs1031282966&location=UTR3&one=1) | chr2:190970064 | T/C | 0.9999/0.0001 | [9](https://guolab.wchscu.cn/miRNASNP/#!/snp?snp_id=rs1031282966&location=UTR3&four='1') | [7](https://guolab.wchscu.cn/miRNASNP/#!/snp?snp_id=rs1031282966&location=UTR3&five='1') |
| [rs1467062599](https://guolab.wchscu.cn/miRNASNP/#!/snp?snp_id=rs1467062599&location=UTR3&one=1) | chr2:190970066 | CTCAAGAAA/C | 1.0/- | [4](https://guolab.wchscu.cn/miRNASNP/#!/snp?snp_id=rs1467062599&location=UTR3&four='1') | [4](https://guolab.wchscu.cn/miRNASNP/#!/snp?snp_id=rs1467062599&location=UTR3&five='1') |
| [rs1392245161](https://guolab.wchscu.cn/miRNASNP/#!/snp?snp_id=rs1392245161&location=UTR3&one=1) | chr2:190970068 | C/G | 1/- | [5](https://guolab.wchscu.cn/miRNASNP/#!/snp?snp_id=rs1392245161&location=UTR3&four='1') | [4](https://guolab.wchscu.cn/miRNASNP/#!/snp?snp_id=rs1392245161&location=UTR3&five='1') |
| [rs965920313](https://guolab.wchscu.cn/miRNASNP/#!/snp?snp_id=rs965920313&location=UTR3&one=1) | chr2:190970070 | A/G | -/- | [8](https://guolab.wchscu.cn/miRNASNP/#!/snp?snp_id=rs965920313&location=UTR3&four='1') | [3](https://guolab.wchscu.cn/miRNASNP/#!/snp?snp_id=rs965920313&location=UTR3&five='1') |
| [rs1353563398](https://guolab.wchscu.cn/miRNASNP/#!/snp?snp_id=rs1353563398&location=UTR3&one=1) | chr2:190970071 | GA/G | -/- | [3](https://guolab.wchscu.cn/miRNASNP/#!/snp?snp_id=rs1353563398&location=UTR3&four='1') | [1](https://guolab.wchscu.cn/miRNASNP/#!/snp?snp_id=rs1353563398&location=UTR3&five='1') |
| [rs975866030](https://guolab.wchscu.cn/miRNASNP/#!/snp?snp_id=rs975866030&location=UTR3&one=1) | chr2:190970075 | C/T | -/- | [1](https://guolab.wchscu.cn/miRNASNP/#!/snp?snp_id=rs975866030&location=UTR3&four='1') | [1](https://guolab.wchscu.cn/miRNASNP/#!/snp?snp_id=rs975866030&location=UTR3&five='1') |
| [rs1163743171](https://guolab.wchscu.cn/miRNASNP/#!/snp?snp_id=rs1163743171&location=UTR3&one=1) | chr2:190970078 | A/T | -/- | [5](https://guolab.wchscu.cn/miRNASNP/#!/snp?snp_id=rs1163743171&location=UTR3&four='1') | 0 |
| [rs770420100](https://guolab.wchscu.cn/miRNASNP/#!/snp?snp_id=rs770420100&location=UTR3&one=1) | chr2:190970081 | TTG/T | -/- | 0 | [1](https://guolab.wchscu.cn/miRNASNP/#!/snp?snp_id=rs770420100&location=UTR3&five='1') |
| [rs936066002](https://guolab.wchscu.cn/miRNASNP/#!/snp?snp_id=rs936066002&location=UTR3&one=1) | chr2:190970085 | CTT/C | -/- | [5](https://guolab.wchscu.cn/miRNASNP/#!/snp?snp_id=rs936066002&location=UTR3&four='1') | [1](https://guolab.wchscu.cn/miRNASNP/#!/snp?snp_id=rs936066002&location=UTR3&five='1') |
| [rs1415740340](https://guolab.wchscu.cn/miRNASNP/#!/snp?snp_id=rs1415740340&location=UTR3&one=1) | chr2:190970091 | C/G | 1/- | [3](https://guolab.wchscu.cn/miRNASNP/#!/snp?snp_id=rs1415740340&location=UTR3&four='1') | [2](https://guolab.wchscu.cn/miRNASNP/#!/snp?snp_id=rs1415740340&location=UTR3&five='1') |
| [rs921851886](https://guolab.wchscu.cn/miRNASNP/#!/snp?snp_id=rs921851886&location=UTR3&one=1) | chr2:190970098 | T/C | 1/- | [2](https://guolab.wchscu.cn/miRNASNP/#!/snp?snp_id=rs921851886&location=UTR3&four='1') | [3](https://guolab.wchscu.cn/miRNASNP/#!/snp?snp_id=rs921851886&location=UTR3&five='1') |
| [rs760945547](https://guolab.wchscu.cn/miRNASNP/#!/snp?snp_id=rs760945547&location=UTR3&one=1) | chr2:190970101 | C/A | 1/- | [1](https://guolab.wchscu.cn/miRNASNP/#!/snp?snp_id=rs760945547&location=UTR3&four='1') | [2](https://guolab.wchscu.cn/miRNASNP/#!/snp?snp_id=rs760945547&location=UTR3&five='1') |
| [rs79073086](https://guolab.wchscu.cn/miRNASNP/#!/snp?snp_id=rs79073086&location=UTR3&one=1) | chr2:190970106 | G/C | -/- | [2](https://guolab.wchscu.cn/miRNASNP/#!/snp?snp_id=rs79073086&location=UTR3&four='1') | [3](https://guolab.wchscu.cn/miRNASNP/#!/snp?snp_id=rs79073086&location=UTR3&five='1') |
| [rs146036682](https://guolab.wchscu.cn/miRNASNP/#!/snp?snp_id=rs146036682&location=UTR3&one=1) | chr2:190970112 | G/A | 1/- | [2](https://guolab.wchscu.cn/miRNASNP/#!/snp?snp_id=rs146036682&location=UTR3&four='1') | [6](https://guolab.wchscu.cn/miRNASNP/#!/snp?snp_id=rs146036682&location=UTR3&five='1') |
| [rs1180591844](https://guolab.wchscu.cn/miRNASNP/#!/snp?snp_id=rs1180591844&location=UTR3&one=1) | chr2:190970114 | T/C | 1/- | [3](https://guolab.wchscu.cn/miRNASNP/#!/snp?snp_id=rs1180591844&location=UTR3&four='1') | [10](https://guolab.wchscu.cn/miRNASNP/#!/snp?snp_id=rs1180591844&location=UTR3&five='1') |
| [rs984455097](https://guolab.wchscu.cn/miRNASNP/#!/snp?snp_id=rs984455097&location=UTR3&one=1) | chr2:190970115 | G/A | -/- | [7](https://guolab.wchscu.cn/miRNASNP/#!/snp?snp_id=rs984455097&location=UTR3&four='1') | [10](https://guolab.wchscu.cn/miRNASNP/#!/snp?snp_id=rs984455097&location=UTR3&five='1') |
| [rs1467489667](https://guolab.wchscu.cn/miRNASNP/#!/snp?snp_id=rs1467489667&location=UTR3&one=1) | chr2:190970128 | CA/C | 1.0/- | [2](https://guolab.wchscu.cn/miRNASNP/#!/snp?snp_id=rs1467489667&location=UTR3&four='1') | [7](https://guolab.wchscu.cn/miRNASNP/#!/snp?snp_id=rs1467489667&location=UTR3&five='1') |
| [rs190508584](https://guolab.wchscu.cn/miRNASNP/#!/snp?snp_id=rs190508584&location=UTR3&one=1) | chr2:190970129 | A/G | -/- | 0 | [7](https://guolab.wchscu.cn/miRNASNP/#!/snp?snp_id=rs190508584&location=UTR3&five='1') |
| [rs114360225](https://guolab.wchscu.cn/miRNASNP/#!/snp?snp_id=rs114360225&location=UTR3&one=1) | chr2:190970130 | T/C | 0.9997/0.0003 | [6](https://guolab.wchscu.cn/miRNASNP/#!/snp?snp_id=rs114360225&location=UTR3&four='1') | [4](https://guolab.wchscu.cn/miRNASNP/#!/snp?snp_id=rs114360225&location=UTR3&five='1') |
| [rs1223001198](https://guolab.wchscu.cn/miRNASNP/#!/snp?snp_id=rs1223001198&location=UTR3&one=1) | chr2:190970133 | T/TC | -/- | 0 | 0 |
| [rs1323876080](https://guolab.wchscu.cn/miRNASNP/#!/snp?snp_id=rs1323876080&location=UTR3&one=1) | chr2:190970144 | G/T | 1/- | 0 | [2](https://guolab.wchscu.cn/miRNASNP/#!/snp?snp_id=rs1323876080&location=UTR3&five='1') |
| [rs907433798](https://guolab.wchscu.cn/miRNASNP/#!/snp?snp_id=rs907433798&location=UTR3&one=1) | chr2:190970146 | T/C | 1/- | [3](https://guolab.wchscu.cn/miRNASNP/#!/snp?snp_id=rs907433798&location=UTR3&four='1') | [4](https://guolab.wchscu.cn/miRNASNP/#!/snp?snp_id=rs907433798&location=UTR3&five='1') |
| [rs1237438235](https://guolab.wchscu.cn/miRNASNP/#!/snp?snp_id=rs1237438235&location=UTR3&one=1) | chr2:190970150 | T/C | 1/- | [1](https://guolab.wchscu.cn/miRNASNP/#!/snp?snp_id=rs1237438235&location=UTR3&four='1') | [5](https://guolab.wchscu.cn/miRNASNP/#!/snp?snp_id=rs1237438235&location=UTR3&five='1') |
| [rs1004290548](https://guolab.wchscu.cn/miRNASNP/#!/snp?snp_id=rs1004290548&location=UTR3&one=1) | chr2:190970155 | G/T | 1/- | [4](https://guolab.wchscu.cn/miRNASNP/#!/snp?snp_id=rs1004290548&location=UTR3&four='1') | [4](https://guolab.wchscu.cn/miRNASNP/#!/snp?snp_id=rs1004290548&location=UTR3&five='1') |
| [rs41476445](https://guolab.wchscu.cn/miRNASNP/#!/snp?snp_id=rs41476445&location=UTR3&one=1) | chr2:190970168 | G/A | 0.9902/0.0098 | [3](https://guolab.wchscu.cn/miRNASNP/#!/snp?snp_id=rs41476445&location=UTR3&four='1') | [9](https://guolab.wchscu.cn/miRNASNP/#!/snp?snp_id=rs41476445&location=UTR3&five='1') |
| [rs1448379540](https://guolab.wchscu.cn/miRNASNP/#!/snp?snp_id=rs1448379540&location=UTR3&one=1) | chr2:190970170 | G/C | 1/- | [2](https://guolab.wchscu.cn/miRNASNP/#!/snp?snp_id=rs1448379540&location=UTR3&four='1') | [14](https://guolab.wchscu.cn/miRNASNP/#!/snp?snp_id=rs1448379540&location=UTR3&five='1') |
| [rs1036098684](https://guolab.wchscu.cn/miRNASNP/#!/snp?snp_id=rs1036098684&location=UTR3&one=1) | chr2:190970171 | G/A | 0.9998/0.0002 | [6](https://guolab.wchscu.cn/miRNASNP/#!/snp?snp_id=rs1036098684&location=UTR3&four='1') | [14](https://guolab.wchscu.cn/miRNASNP/#!/snp?snp_id=rs1036098684&location=UTR3&five='1') |
| [rs1292838083](https://guolab.wchscu.cn/miRNASNP/#!/snp?snp_id=rs1292838083&location=UTR3&one=1) | chr2:190970172 | A/G | 1/- | [15](https://guolab.wchscu.cn/miRNASNP/#!/snp?snp_id=rs1292838083&location=UTR3&four='1') | [14](https://guolab.wchscu.cn/miRNASNP/#!/snp?snp_id=rs1292838083&location=UTR3&five='1') |
| [rs1437185939](https://guolab.wchscu.cn/miRNASNP/#!/snp?snp_id=rs1437185939&location=UTR3&one=1) | chr2:190970179 | T/C | -/- | [4](https://guolab.wchscu.cn/miRNASNP/#!/snp?snp_id=rs1437185939&location=UTR3&four='1') | [3](https://guolab.wchscu.cn/miRNASNP/#!/snp?snp_id=rs1437185939&location=UTR3&five='1') |
| [rs1351391769](https://guolab.wchscu.cn/miRNASNP/#!/snp?snp_id=rs1351391769&location=UTR3&one=1) | chr2:190970182 | T/C | 1/- | [2](https://guolab.wchscu.cn/miRNASNP/#!/snp?snp_id=rs1351391769&location=UTR3&four='1') | [3](https://guolab.wchscu.cn/miRNASNP/#!/snp?snp_id=rs1351391769&location=UTR3&five='1') |
| [rs1156626637](https://guolab.wchscu.cn/miRNASNP/#!/snp?snp_id=rs1156626637&location=UTR3&one=1) | chr2:190970184 | G/A | 1/- | [3](https://guolab.wchscu.cn/miRNASNP/#!/snp?snp_id=rs1156626637&location=UTR3&four='1') | [2](https://guolab.wchscu.cn/miRNASNP/#!/snp?snp_id=rs1156626637&location=UTR3&five='1') |
| [rs1448097711](https://guolab.wchscu.cn/miRNASNP/#!/snp?snp_id=rs1448097711&location=UTR3&one=1) | chr2:190970188 | A/G | 1/- | [13](https://guolab.wchscu.cn/miRNASNP/#!/snp?snp_id=rs1448097711&location=UTR3&four='1') | [3](https://guolab.wchscu.cn/miRNASNP/#!/snp?snp_id=rs1448097711&location=UTR3&five='1') |
| [rs1381185396](https://guolab.wchscu.cn/miRNASNP/#!/snp?snp_id=rs1381185396&location=UTR3&one=1) | chr2:190970193 | C/T | 1/- | [4](https://guolab.wchscu.cn/miRNASNP/#!/snp?snp_id=rs1381185396&location=UTR3&four='1') | [2](https://guolab.wchscu.cn/miRNASNP/#!/snp?snp_id=rs1381185396&location=UTR3&five='1') |
| [rs1160732426](https://guolab.wchscu.cn/miRNASNP/#!/snp?snp_id=rs1160732426&location=UTR3&one=1) | chr2:190970198 | C/T | 1/- | [6](https://guolab.wchscu.cn/miRNASNP/#!/snp?snp_id=rs1160732426&location=UTR3&four='1') | [2](https://guolab.wchscu.cn/miRNASNP/#!/snp?snp_id=rs1160732426&location=UTR3&five='1') |
| [rs754155168](https://guolab.wchscu.cn/miRNASNP/#!/snp?snp_id=rs754155168&location=UTR3&one=1) | chr2:190970200 | T/C | -/- | [11](https://guolab.wchscu.cn/miRNASNP/#!/snp?snp_id=rs754155168&location=UTR3&four='1') | [2](https://guolab.wchscu.cn/miRNASNP/#!/snp?snp_id=rs754155168&location=UTR3&five='1') |
| [rs1452625062](https://guolab.wchscu.cn/miRNASNP/#!/snp?snp_id=rs1452625062&location=UTR3&one=1) | chr2:190970201 | T/C | 1/- | [1](https://guolab.wchscu.cn/miRNASNP/#!/snp?snp_id=rs1452625062&location=UTR3&four='1') | [4](https://guolab.wchscu.cn/miRNASNP/#!/snp?snp_id=rs1452625062&location=UTR3&five='1') |
| [rs1187379026](https://guolab.wchscu.cn/miRNASNP/#!/snp?snp_id=rs1187379026&location=UTR3&one=1) | chr2:190970205 | C/A | -/- | [2](https://guolab.wchscu.cn/miRNASNP/#!/snp?snp_id=rs1187379026&location=UTR3&four='1') | [3](https://guolab.wchscu.cn/miRNASNP/#!/snp?snp_id=rs1187379026&location=UTR3&five='1') |
| [rs938880825](https://guolab.wchscu.cn/miRNASNP/#!/snp?snp_id=rs938880825&location=UTR3&one=1) | chr2:190970221 | C/T | -/- | [1](https://guolab.wchscu.cn/miRNASNP/#!/snp?snp_id=rs938880825&location=UTR3&four='1') | [6](https://guolab.wchscu.cn/miRNASNP/#!/snp?snp_id=rs938880825&location=UTR3&five='1') |
| [rs1266849259](https://guolab.wchscu.cn/miRNASNP/#!/snp?snp_id=rs1266849259&location=UTR3&one=1) | chr2:190970227 | GA/G | -/- | [3](https://guolab.wchscu.cn/miRNASNP/#!/snp?snp_id=rs1266849259&location=UTR3&four='1') | [10](https://guolab.wchscu.cn/miRNASNP/#!/snp?snp_id=rs1266849259&location=UTR3&five='1') |
| [rs1192270071](https://guolab.wchscu.cn/miRNASNP/#!/snp?snp_id=rs1192270071&location=UTR3&one=1) | chr2:190970234 | G/A | 1/- | [2](https://guolab.wchscu.cn/miRNASNP/#!/snp?snp_id=rs1192270071&location=UTR3&four='1') | [7](https://guolab.wchscu.cn/miRNASNP/#!/snp?snp_id=rs1192270071&location=UTR3&five='1') |
| [rs201656062](https://guolab.wchscu.cn/miRNASNP/#!/snp?snp_id=rs201656062&location=UTR3&one=1) | chr2:190970241 | AC/A | 0.9994/0.0006 | 0 | 0 |
| [rs200344731](https://guolab.wchscu.cn/miRNASNP/#!/snp?snp_id=rs200344731&location=UTR3&one=1) | chr2:190970242 | C/A | -/- | 0 | 0 |
| [rs1240659511](https://guolab.wchscu.cn/miRNASNP/#!/snp?snp_id=rs1240659511&location=UTR3&one=1) | chr2:190970250 | G/A | 1/- | [4](https://guolab.wchscu.cn/miRNASNP/#!/snp?snp_id=rs1240659511&location=UTR3&four='1') | [5](https://guolab.wchscu.cn/miRNASNP/#!/snp?snp_id=rs1240659511&location=UTR3&five='1') |
| [rs1055938745](https://guolab.wchscu.cn/miRNASNP/#!/snp?snp_id=rs1055938745&location=UTR3&one=1) | chr2:190970260 | C/T | -/- | [3](https://guolab.wchscu.cn/miRNASNP/#!/snp?snp_id=rs1055938745&location=UTR3&four='1') | [1](https://guolab.wchscu.cn/miRNASNP/#!/snp?snp_id=rs1055938745&location=UTR3&five='1') |
| [rs1473792625](https://guolab.wchscu.cn/miRNASNP/#!/snp?snp_id=rs1473792625&location=UTR3&one=1) | chr2:190970261 | A/G | -/- | [2](https://guolab.wchscu.cn/miRNASNP/#!/snp?snp_id=rs1473792625&location=UTR3&four='1') | [4](https://guolab.wchscu.cn/miRNASNP/#!/snp?snp_id=rs1473792625&location=UTR3&five='1') |
| [rs529316850](https://guolab.wchscu.cn/miRNASNP/#!/snp?snp_id=rs529316850&location=UTR3&one=1) | chr2:190970263 | G/T | 1/- | [4](https://guolab.wchscu.cn/miRNASNP/#!/snp?snp_id=rs529316850&location=UTR3&four='1') | [4](https://guolab.wchscu.cn/miRNASNP/#!/snp?snp_id=rs529316850&location=UTR3&five='1') |
| [rs1352469364](https://guolab.wchscu.cn/miRNASNP/#!/snp?snp_id=rs1352469364&location=UTR3&one=1) | chr2:190970266 | A/C | 1/- | [1](https://guolab.wchscu.cn/miRNASNP/#!/snp?snp_id=rs1352469364&location=UTR3&four='1') | [5](https://guolab.wchscu.cn/miRNASNP/#!/snp?snp_id=rs1352469364&location=UTR3&five='1') |
| [rs1286254077](https://guolab.wchscu.cn/miRNASNP/#!/snp?snp_id=rs1286254077&location=UTR3&one=1) | chr2:190970274 | C/T | 1/- | [4](https://guolab.wchscu.cn/miRNASNP/#!/snp?snp_id=rs1286254077&location=UTR3&four='1') | [10](https://guolab.wchscu.cn/miRNASNP/#!/snp?snp_id=rs1286254077&location=UTR3&five='1') |
| [rs139958571](https://guolab.wchscu.cn/miRNASNP/#!/snp?snp_id=rs139958571&location=UTR3&one=1) | chr2:190970275 | C/G | 0.9992/0.0008 | [2](https://guolab.wchscu.cn/miRNASNP/#!/snp?snp_id=rs139958571&location=UTR3&four='1') | [10](https://guolab.wchscu.cn/miRNASNP/#!/snp?snp_id=rs139958571&location=UTR3&five='1') |
| [rs562899714](https://guolab.wchscu.cn/miRNASNP/#!/snp?snp_id=rs562899714&location=UTR3&one=1) | chr2:190970276 | C/A | -/- | [5](https://guolab.wchscu.cn/miRNASNP/#!/snp?snp_id=rs562899714&location=UTR3&four='1') | [9](https://guolab.wchscu.cn/miRNASNP/#!/snp?snp_id=rs562899714&location=UTR3&five='1') |
| [rs562899714](https://guolab.wchscu.cn/miRNASNP/#!/snp?snp_id=rs562899714&location=UTR3&one=1) | chr2:190970276 | C/T | -/- | [3](https://guolab.wchscu.cn/miRNASNP/#!/snp?snp_id=rs562899714&location=UTR3&four='1') | [9](https://guolab.wchscu.cn/miRNASNP/#!/snp?snp_id=rs562899714&location=UTR3&five='1') |
| [rs1309927096](https://guolab.wchscu.cn/miRNASNP/#!/snp?snp_id=rs1309927096&location=UTR3&one=1) | chr2:190970286 | T/C | 1/- | [2](https://guolab.wchscu.cn/miRNASNP/#!/snp?snp_id=rs1309927096&location=UTR3&four='1') | [5](https://guolab.wchscu.cn/miRNASNP/#!/snp?snp_id=rs1309927096&location=UTR3&five='1') |
| [rs1300061520](https://guolab.wchscu.cn/miRNASNP/#!/snp?snp_id=rs1300061520&location=UTR3&one=1) | chr2:190970288 | C/T | 1/- | 0 | [1](https://guolab.wchscu.cn/miRNASNP/#!/snp?snp_id=rs1300061520&location=UTR3&five='1') |
| [rs1406977266](https://guolab.wchscu.cn/miRNASNP/#!/snp?snp_id=rs1406977266&location=UTR3&one=1) | chr2:190970289 | C/T | -/- | [5](https://guolab.wchscu.cn/miRNASNP/#!/snp?snp_id=rs1406977266&location=UTR3&four='1') | 0 |
| [rs1156677557](https://guolab.wchscu.cn/miRNASNP/#!/snp?snp_id=rs1156677557&location=UTR3&one=1) | chr2:190970290 | G/A | 1/- | [1](https://guolab.wchscu.cn/miRNASNP/#!/snp?snp_id=rs1156677557&location=UTR3&four='1') | [1](https://guolab.wchscu.cn/miRNASNP/#!/snp?snp_id=rs1156677557&location=UTR3&five='1') |
| [rs182394503](https://guolab.wchscu.cn/miRNASNP/#!/snp?snp_id=rs182394503&location=UTR3&one=1) | chr2:190970297 | C/T | 0.9999/0.0001 | [2](https://guolab.wchscu.cn/miRNASNP/#!/snp?snp_id=rs182394503&location=UTR3&four='1') | [3](https://guolab.wchscu.cn/miRNASNP/#!/snp?snp_id=rs182394503&location=UTR3&five='1') |
| [rs1296073393](https://guolab.wchscu.cn/miRNASNP/#!/snp?snp_id=rs1296073393&location=UTR3&one=1) | chr2:190970298 | G/A | 1/- | [2](https://guolab.wchscu.cn/miRNASNP/#!/snp?snp_id=rs1296073393&location=UTR3&four='1') | [2](https://guolab.wchscu.cn/miRNASNP/#!/snp?snp_id=rs1296073393&location=UTR3&five='1') |
| [rs1401221651](https://guolab.wchscu.cn/miRNASNP/#!/snp?snp_id=rs1401221651&location=UTR3&one=1) | chr2:190970307 | T/C | 1/- | [15](https://guolab.wchscu.cn/miRNASNP/#!/snp?snp_id=rs1401221651&location=UTR3&four='1') | [7](https://guolab.wchscu.cn/miRNASNP/#!/snp?snp_id=rs1401221651&location=UTR3&five='1') |
| [rs775525919](https://guolab.wchscu.cn/miRNASNP/#!/snp?snp_id=rs775525919&location=UTR3&one=1) | chr2:190970313 | AAAT/A | -/- | [2](https://guolab.wchscu.cn/miRNASNP/#!/snp?snp_id=rs775525919&location=UTR3&four='1') | [2](https://guolab.wchscu.cn/miRNASNP/#!/snp?snp_id=rs775525919&location=UTR3&five='1') |
| [rs1360976235](https://guolab.wchscu.cn/miRNASNP/#!/snp?snp_id=rs1360976235&location=UTR3&one=1) | chr2:190970321 | ACT/A | -/- | [2](https://guolab.wchscu.cn/miRNASNP/#!/snp?snp_id=rs1360976235&location=UTR3&four='1') | [6](https://guolab.wchscu.cn/miRNASNP/#!/snp?snp_id=rs1360976235&location=UTR3&five='1') |
| [rs1300275650](https://guolab.wchscu.cn/miRNASNP/#!/snp?snp_id=rs1300275650&location=UTR3&one=1) | chr2:190970337 | A/T | 1/- | [4](https://guolab.wchscu.cn/miRNASNP/#!/snp?snp_id=rs1300275650&location=UTR3&four='1') | [3](https://guolab.wchscu.cn/miRNASNP/#!/snp?snp_id=rs1300275650&location=UTR3&five='1') |
| [rs186032149](https://guolab.wchscu.cn/miRNASNP/#!/snp?snp_id=rs186032149&location=UTR3&one=1) | chr2:190970339 | C/G | 0.9995/0.0005 | 0 | [1](https://guolab.wchscu.cn/miRNASNP/#!/snp?snp_id=rs186032149&location=UTR3&five='1') |
| [rs886055377](https://guolab.wchscu.cn/miRNASNP/#!/snp?snp_id=rs886055377&location=UTR3&one=1) | chr2:190970341 | A/G | 1/- | [4](https://guolab.wchscu.cn/miRNASNP/#!/snp?snp_id=rs886055377&location=UTR3&four='1') | 0 |
| [rs886055377](https://guolab.wchscu.cn/miRNASNP/#!/snp?snp_id=rs886055377&location=UTR3&one=1) | chr2:190970341 | A/T | 1/- | [5](https://guolab.wchscu.cn/miRNASNP/#!/snp?snp_id=rs886055377&location=UTR3&four='1') | 0 |
| [rs903172029](https://guolab.wchscu.cn/miRNASNP/#!/snp?snp_id=rs903172029&location=UTR3&one=1) | chr2:190970342 | T/C | -/- | [3](https://guolab.wchscu.cn/miRNASNP/#!/snp?snp_id=rs903172029&location=UTR3&four='1') | 0 |
| [rs566349352](https://guolab.wchscu.cn/miRNASNP/#!/snp?snp_id=rs566349352&location=UTR3&one=1) | chr2:190970346 | A/G | 1/- | [1](https://guolab.wchscu.cn/miRNASNP/#!/snp?snp_id=rs566349352&location=UTR3&four='1') | [4](https://guolab.wchscu.cn/miRNASNP/#!/snp?snp_id=rs566349352&location=UTR3&five='1') |
| [rs566349352](https://guolab.wchscu.cn/miRNASNP/#!/snp?snp_id=rs566349352&location=UTR3&one=1) | chr2:190970346 | A/T | 1/- | [4](https://guolab.wchscu.cn/miRNASNP/#!/snp?snp_id=rs566349352&location=UTR3&four='1') | [4](https://guolab.wchscu.cn/miRNASNP/#!/snp?snp_id=rs566349352&location=UTR3&five='1') |
| [rs977832294](https://guolab.wchscu.cn/miRNASNP/#!/snp?snp_id=rs977832294&location=UTR3&one=1) | chr2:190970348 | T/C | 1/- | [9](https://guolab.wchscu.cn/miRNASNP/#!/snp?snp_id=rs977832294&location=UTR3&four='1') | [6](https://guolab.wchscu.cn/miRNASNP/#!/snp?snp_id=rs977832294&location=UTR3&five='1') |
| [rs1193632643](https://guolab.wchscu.cn/miRNASNP/#!/snp?snp_id=rs1193632643&location=UTR3&one=1) | chr2:190970364 | T/G | 1/- | [1](https://guolab.wchscu.cn/miRNASNP/#!/snp?snp_id=rs1193632643&location=UTR3&four='1') | [4](https://guolab.wchscu.cn/miRNASNP/#!/snp?snp_id=rs1193632643&location=UTR3&five='1') |
| [rs777014385](https://guolab.wchscu.cn/miRNASNP/#!/snp?snp_id=rs777014385&location=UTR3&one=1) | chr2:190970367 | A/G | -/- | [1](https://guolab.wchscu.cn/miRNASNP/#!/snp?snp_id=rs777014385&location=UTR3&four='1') | [3](https://guolab.wchscu.cn/miRNASNP/#!/snp?snp_id=rs777014385&location=UTR3&five='1') |
| [rs1395511430](https://guolab.wchscu.cn/miRNASNP/#!/snp?snp_id=rs1395511430&location=UTR3&one=1) | chr2:190970437 | G/T | -/- | [6](https://guolab.wchscu.cn/miRNASNP/#!/snp?snp_id=rs1395511430&location=UTR3&four='1') | [8](https://guolab.wchscu.cn/miRNASNP/#!/snp?snp_id=rs1395511430&location=UTR3&five='1') |
| [rs182725919](https://guolab.wchscu.cn/miRNASNP/#!/snp?snp_id=rs182725919&location=UTR3&one=1) | chr2:190970440 | T/C | 1/- | 0 | [4](https://guolab.wchscu.cn/miRNASNP/#!/snp?snp_id=rs182725919&location=UTR3&five='1') |
| [rs1401808260](https://guolab.wchscu.cn/miRNASNP/#!/snp?snp_id=rs1401808260&location=UTR3&one=1) | chr2:190970442 | T/C | 1/- | [5](https://guolab.wchscu.cn/miRNASNP/#!/snp?snp_id=rs1401808260&location=UTR3&four='1') | [7](https://guolab.wchscu.cn/miRNASNP/#!/snp?snp_id=rs1401808260&location=UTR3&five='1') |
| [rs1303456711](https://guolab.wchscu.cn/miRNASNP/#!/snp?snp_id=rs1303456711&location=UTR3&one=1) | chr2:190970446 | G/A | 1/- | [6](https://guolab.wchscu.cn/miRNASNP/#!/snp?snp_id=rs1303456711&location=UTR3&four='1') | [10](https://guolab.wchscu.cn/miRNASNP/#!/snp?snp_id=rs1303456711&location=UTR3&five='1') |
| [rs1045851098](https://guolab.wchscu.cn/miRNASNP/#!/snp?snp_id=rs1045851098&location=UTR3&one=1) | chr2:190970457 | C/CT | -/- | [2](https://guolab.wchscu.cn/miRNASNP/#!/snp?snp_id=rs1045851098&location=UTR3&four='1') | [3](https://guolab.wchscu.cn/miRNASNP/#!/snp?snp_id=rs1045851098&location=UTR3&five='1') |
| [rs984653102](https://guolab.wchscu.cn/miRNASNP/#!/snp?snp_id=rs984653102&location=UTR3&one=1) | chr2:190970457 | C/G | -/- | [3](https://guolab.wchscu.cn/miRNASNP/#!/snp?snp_id=rs984653102&location=UTR3&four='1') | [3](https://guolab.wchscu.cn/miRNASNP/#!/snp?snp_id=rs984653102&location=UTR3&five='1') |
| [rs1204464083](https://guolab.wchscu.cn/miRNASNP/#!/snp?snp_id=rs1204464083&location=UTR3&one=1) | chr2:190970466 | C/A | -/- | [10](https://guolab.wchscu.cn/miRNASNP/#!/snp?snp_id=rs1204464083&location=UTR3&four='1') | [2](https://guolab.wchscu.cn/miRNASNP/#!/snp?snp_id=rs1204464083&location=UTR3&five='1') |
| [rs1398246936](https://guolab.wchscu.cn/miRNASNP/#!/snp?snp_id=rs1398246936&location=UTR3&one=1) | chr2:190970467 | C/G | 1/- | [3](https://guolab.wchscu.cn/miRNASNP/#!/snp?snp_id=rs1398246936&location=UTR3&four='1') | [3](https://guolab.wchscu.cn/miRNASNP/#!/snp?snp_id=rs1398246936&location=UTR3&five='1') |
| [rs1237627664](https://guolab.wchscu.cn/miRNASNP/#!/snp?snp_id=rs1237627664&location=UTR3&one=1) | chr2:190970488 | T/C | -/- | [4](https://guolab.wchscu.cn/miRNASNP/#!/snp?snp_id=rs1237627664&location=UTR3&four='1') | [1](https://guolab.wchscu.cn/miRNASNP/#!/snp?snp_id=rs1237627664&location=UTR3&five='1') |
| [rs1016371601](https://guolab.wchscu.cn/miRNASNP/#!/snp?snp_id=rs1016371601&location=UTR3&one=1) | chr2:190970489 | C/T | -/- | [1](https://guolab.wchscu.cn/miRNASNP/#!/snp?snp_id=rs1016371601&location=UTR3&four='1') | [1](https://guolab.wchscu.cn/miRNASNP/#!/snp?snp_id=rs1016371601&location=UTR3&five='1') |
| [rs907458830](https://guolab.wchscu.cn/miRNASNP/#!/snp?snp_id=rs907458830&location=UTR3&one=1) | chr2:190970491 | A/G | 1/- | [4](https://guolab.wchscu.cn/miRNASNP/#!/snp?snp_id=rs907458830&location=UTR3&four='1') | [2](https://guolab.wchscu.cn/miRNASNP/#!/snp?snp_id=rs907458830&location=UTR3&five='1') |
| [rs961749006](https://guolab.wchscu.cn/miRNASNP/#!/snp?snp_id=rs961749006&location=UTR3&one=1) | chr2:190970492 | T/A | -/- | [6](https://guolab.wchscu.cn/miRNASNP/#!/snp?snp_id=rs961749006&location=UTR3&four='1') | [2](https://guolab.wchscu.cn/miRNASNP/#!/snp?snp_id=rs961749006&location=UTR3&five='1') |
| [rs1004504808](https://guolab.wchscu.cn/miRNASNP/#!/snp?snp_id=rs1004504808&location=UTR3&one=1) | chr2:190970511 | C/T | 1/- | [4](https://guolab.wchscu.cn/miRNASNP/#!/snp?snp_id=rs1004504808&location=UTR3&four='1') | [5](https://guolab.wchscu.cn/miRNASNP/#!/snp?snp_id=rs1004504808&location=UTR3&five='1') |
| [rs1180667225](https://guolab.wchscu.cn/miRNASNP/#!/snp?snp_id=rs1180667225&location=UTR3&one=1) | chr2:190970511 | CA/C | -/- | [5](https://guolab.wchscu.cn/miRNASNP/#!/snp?snp_id=rs1180667225&location=UTR3&four='1') | [6](https://guolab.wchscu.cn/miRNASNP/#!/snp?snp_id=rs1180667225&location=UTR3&five='1') |
| [rs1426152930](https://guolab.wchscu.cn/miRNASNP/#!/snp?snp_id=rs1426152930&location=UTR3&one=1) | chr2:190970520 | CT/C | -/- | [2](https://guolab.wchscu.cn/miRNASNP/#!/snp?snp_id=rs1426152930&location=UTR3&four='1') | [4](https://guolab.wchscu.cn/miRNASNP/#!/snp?snp_id=rs1426152930&location=UTR3&five='1') |
| [rs537942049](https://guolab.wchscu.cn/miRNASNP/#!/snp?snp_id=rs537942049&location=UTR3&one=1) | chr2:190970522 | T/C | -/- | [7](https://guolab.wchscu.cn/miRNASNP/#!/snp?snp_id=rs537942049&location=UTR3&four='1') | [5](https://guolab.wchscu.cn/miRNASNP/#!/snp?snp_id=rs537942049&location=UTR3&five='1') |
| [rs750941607](https://guolab.wchscu.cn/miRNASNP/#!/snp?snp_id=rs750941607&location=UTR3&one=1) | chr2:190970523 | C/T | -/- | [2](https://guolab.wchscu.cn/miRNASNP/#!/snp?snp_id=rs750941607&location=UTR3&four='1') | [4](https://guolab.wchscu.cn/miRNASNP/#!/snp?snp_id=rs750941607&location=UTR3&five='1') |
| [rs1365351649](https://guolab.wchscu.cn/miRNASNP/#!/snp?snp_id=rs1365351649&location=UTR3&one=1) | chr2:190970529 | A/G | -/- | [13](https://guolab.wchscu.cn/miRNASNP/#!/snp?snp_id=rs1365351649&location=UTR3&four='1') | [1](https://guolab.wchscu.cn/miRNASNP/#!/snp?snp_id=rs1365351649&location=UTR3&five='1') |
| [rs1470359868](https://guolab.wchscu.cn/miRNASNP/#!/snp?snp_id=rs1470359868&location=UTR3&one=1) | chr2:190970531 | C/G | 1/- | [3](https://guolab.wchscu.cn/miRNASNP/#!/snp?snp_id=rs1470359868&location=UTR3&four='1') | [2](https://guolab.wchscu.cn/miRNASNP/#!/snp?snp_id=rs1470359868&location=UTR3&five='1') |
| [rs972115850](https://guolab.wchscu.cn/miRNASNP/#!/snp?snp_id=rs972115850&location=UTR3&one=1) | chr2:190970538 | T/A | -/- | [2](https://guolab.wchscu.cn/miRNASNP/#!/snp?snp_id=rs972115850&location=UTR3&four='1') | [2](https://guolab.wchscu.cn/miRNASNP/#!/snp?snp_id=rs972115850&location=UTR3&five='1') |
| [rs1421648128](https://guolab.wchscu.cn/miRNASNP/#!/snp?snp_id=rs1421648128&location=UTR3&one=1) | chr2:190970541 | T/C | 0.9999/0.0001 | 0 | [4](https://guolab.wchscu.cn/miRNASNP/#!/snp?snp_id=rs1421648128&location=UTR3&five='1') |
| [rs41481847](https://guolab.wchscu.cn/miRNASNP/#!/snp?snp_id=rs41481847&location=UTR3&one=1) | chr2:190970549 | A/G | 0.9907/0.0093 | [13](https://guolab.wchscu.cn/miRNASNP/#!/snp?snp_id=rs41481847&location=UTR3&four='1') | [4](https://guolab.wchscu.cn/miRNASNP/#!/snp?snp_id=rs41481847&location=UTR3&five='1') |
| [rs1200715284](https://guolab.wchscu.cn/miRNASNP/#!/snp?snp_id=rs1200715284&location=UTR3&one=1) | chr2:190970560 | C/A | 1/- | [2](https://guolab.wchscu.cn/miRNASNP/#!/snp?snp_id=rs1200715284&location=UTR3&four='1') | [2](https://guolab.wchscu.cn/miRNASNP/#!/snp?snp_id=rs1200715284&location=UTR3&five='1') |
| [rs768945585](https://guolab.wchscu.cn/miRNASNP/#!/snp?snp_id=rs768945585&location=UTR3&one=1) | chr2:190970562 | T/G | 1/- | [13](https://guolab.wchscu.cn/miRNASNP/#!/snp?snp_id=rs768945585&location=UTR3&four='1') | [2](https://guolab.wchscu.cn/miRNASNP/#!/snp?snp_id=rs768945585&location=UTR3&five='1') |
| [rs571207686](https://guolab.wchscu.cn/miRNASNP/#!/snp?snp_id=rs571207686&location=UTR3&one=1) | chr2:190970566 | A/G | 1/- | [5](https://guolab.wchscu.cn/miRNASNP/#!/snp?snp_id=rs571207686&location=UTR3&four='1') | 0 |
| [rs1197872838](https://guolab.wchscu.cn/miRNASNP/#!/snp?snp_id=rs1197872838&location=UTR3&one=1) | chr2:190970573 | G/T | 1/- | [2](https://guolab.wchscu.cn/miRNASNP/#!/snp?snp_id=rs1197872838&location=UTR3&four='1') | [3](https://guolab.wchscu.cn/miRNASNP/#!/snp?snp_id=rs1197872838&location=UTR3&five='1') |
| [rs1240953570](https://guolab.wchscu.cn/miRNASNP/#!/snp?snp_id=rs1240953570&location=UTR3&one=1) | chr2:190970581 | C/T | -/- | [5](https://guolab.wchscu.cn/miRNASNP/#!/snp?snp_id=rs1240953570&location=UTR3&four='1') | [3](https://guolab.wchscu.cn/miRNASNP/#!/snp?snp_id=rs1240953570&location=UTR3&five='1') |
| [rs938403650](https://guolab.wchscu.cn/miRNASNP/#!/snp?snp_id=rs938403650&location=UTR3&one=1) | chr2:190970582 | G/A | 0.9997/0.0003 | [7](https://guolab.wchscu.cn/miRNASNP/#!/snp?snp_id=rs938403650&location=UTR3&four='1') | 0 |
| [rs1259708984](https://guolab.wchscu.cn/miRNASNP/#!/snp?snp_id=rs1259708984&location=UTR3&one=1) | chr2:190970584 | A/G | 1/- | 0 | 0 |
| [rs1242374810](https://guolab.wchscu.cn/miRNASNP/#!/snp?snp_id=rs1242374810&location=UTR3&one=1) | chr2:190970593 | CCTTT/C | -/- | 0 | 0 |
| [rs991696488](https://guolab.wchscu.cn/miRNASNP/#!/snp?snp_id=rs991696488&location=UTR3&one=1) | chr2:190970601 | T/C | 1/- | [4](https://guolab.wchscu.cn/miRNASNP/#!/snp?snp_id=rs991696488&location=UTR3&four='1') | [3](https://guolab.wchscu.cn/miRNASNP/#!/snp?snp_id=rs991696488&location=UTR3&five='1') |
| [rs1313664856](https://guolab.wchscu.cn/miRNASNP/#!/snp?snp_id=rs1313664856&location=UTR3&one=1) | chr2:190970610 | T/C | 1/- | [3](https://guolab.wchscu.cn/miRNASNP/#!/snp?snp_id=rs1313664856&location=UTR3&four='1') | [5](https://guolab.wchscu.cn/miRNASNP/#!/snp?snp_id=rs1313664856&location=UTR3&five='1') |
| [rs1398072235](https://guolab.wchscu.cn/miRNASNP/#!/snp?snp_id=rs1398072235&location=UTR3&one=1) | chr2:190970614 | AAC/A | -/- | [1](https://guolab.wchscu.cn/miRNASNP/#!/snp?snp_id=rs1398072235&location=UTR3&four='1') | [1](https://guolab.wchscu.cn/miRNASNP/#!/snp?snp_id=rs1398072235&location=UTR3&five='1') |
| [rs1028090653](https://guolab.wchscu.cn/miRNASNP/#!/snp?snp_id=rs1028090653&location=UTR3&one=1) | chr2:190970618 | C/A | 1/- | [5](https://guolab.wchscu.cn/miRNASNP/#!/snp?snp_id=rs1028090653&location=UTR3&four='1') | [2](https://guolab.wchscu.cn/miRNASNP/#!/snp?snp_id=rs1028090653&location=UTR3&five='1') |
| [rs1028090653](https://guolab.wchscu.cn/miRNASNP/#!/snp?snp_id=rs1028090653&location=UTR3&one=1) | chr2:190970618 | C/T | 1/- | [2](https://guolab.wchscu.cn/miRNASNP/#!/snp?snp_id=rs1028090653&location=UTR3&four='1') | [2](https://guolab.wchscu.cn/miRNASNP/#!/snp?snp_id=rs1028090653&location=UTR3&five='1') |
| [rs1330776493](https://guolab.wchscu.cn/miRNASNP/#!/snp?snp_id=rs1330776493&location=UTR3&one=1) | chr2:190970623 | T/C | 1/- | [2](https://guolab.wchscu.cn/miRNASNP/#!/snp?snp_id=rs1330776493&location=UTR3&four='1') | [6](https://guolab.wchscu.cn/miRNASNP/#!/snp?snp_id=rs1330776493&location=UTR3&five='1') |
| [rs1288692855](https://guolab.wchscu.cn/miRNASNP/#!/snp?snp_id=rs1288692855&location=UTR3&one=1) | chr2:190970630 | G/T | -/- | [6](https://guolab.wchscu.cn/miRNASNP/#!/snp?snp_id=rs1288692855&location=UTR3&four='1') | [6](https://guolab.wchscu.cn/miRNASNP/#!/snp?snp_id=rs1288692855&location=UTR3&five='1') |
| [rs1444165759](https://guolab.wchscu.cn/miRNASNP/#!/snp?snp_id=rs1444165759&location=UTR3&one=1) | chr2:190970632 | A/G | 1/- | [10](https://guolab.wchscu.cn/miRNASNP/#!/snp?snp_id=rs1444165759&location=UTR3&four='1') | [4](https://guolab.wchscu.cn/miRNASNP/#!/snp?snp_id=rs1444165759&location=UTR3&five='1') |
| [rs1403123349](https://guolab.wchscu.cn/miRNASNP/#!/snp?snp_id=rs1403123349&location=UTR3&one=1) | chr2:190970646 | G/A | -/- | [5](https://guolab.wchscu.cn/miRNASNP/#!/snp?snp_id=rs1403123349&location=UTR3&four='1') | [18](https://guolab.wchscu.cn/miRNASNP/#!/snp?snp_id=rs1403123349&location=UTR3&five='1') |
| [rs916064299](https://guolab.wchscu.cn/miRNASNP/#!/snp?snp_id=rs916064299&location=UTR3&one=1) | chr2:190970650 | A/G | -/- | [8](https://guolab.wchscu.cn/miRNASNP/#!/snp?snp_id=rs916064299&location=UTR3&four='1') | [7](https://guolab.wchscu.cn/miRNASNP/#!/snp?snp_id=rs916064299&location=UTR3&five='1') |
| [rs548086115](https://guolab.wchscu.cn/miRNASNP/#!/snp?snp_id=rs548086115&location=UTR3&one=1) | chr2:190970652 | C/A | -/- | 0 | [6](https://guolab.wchscu.cn/miRNASNP/#!/snp?snp_id=rs548086115&location=UTR3&five='1') |
| [rs749380631](https://guolab.wchscu.cn/miRNASNP/#!/snp?snp_id=rs749380631&location=UTR3&one=1) | chr2:190970655 | A/G | -/- | [4](https://guolab.wchscu.cn/miRNASNP/#!/snp?snp_id=rs749380631&location=UTR3&four='1') | [6](https://guolab.wchscu.cn/miRNASNP/#!/snp?snp_id=rs749380631&location=UTR3&five='1') |
| [rs1161774047](https://guolab.wchscu.cn/miRNASNP/#!/snp?snp_id=rs1161774047&location=UTR3&one=1) | chr2:190970659 | G/A | 1/- | [1](https://guolab.wchscu.cn/miRNASNP/#!/snp?snp_id=rs1161774047&location=UTR3&four='1') | [5](https://guolab.wchscu.cn/miRNASNP/#!/snp?snp_id=rs1161774047&location=UTR3&five='1') |
| [rs953752983](https://guolab.wchscu.cn/miRNASNP/#!/snp?snp_id=rs953752983&location=UTR3&one=1) | chr2:190970661 | G/C | 1/- | [3](https://guolab.wchscu.cn/miRNASNP/#!/snp?snp_id=rs953752983&location=UTR3&four='1') | [6](https://guolab.wchscu.cn/miRNASNP/#!/snp?snp_id=rs953752983&location=UTR3&five='1') |
| [rs768700585](https://guolab.wchscu.cn/miRNASNP/#!/snp?snp_id=rs768700585&location=UTR3&one=1) | chr2:190970664 | G/C | -/- | [4](https://guolab.wchscu.cn/miRNASNP/#!/snp?snp_id=rs768700585&location=UTR3&four='1') | [10](https://guolab.wchscu.cn/miRNASNP/#!/snp?snp_id=rs768700585&location=UTR3&five='1') |
| [rs1218764714](https://guolab.wchscu.cn/miRNASNP/#!/snp?snp_id=rs1218764714&location=UTR3&one=1) | chr2:190970669 | A/T | -/- | [2](https://guolab.wchscu.cn/miRNASNP/#!/snp?snp_id=rs1218764714&location=UTR3&four='1') | [12](https://guolab.wchscu.cn/miRNASNP/#!/snp?snp_id=rs1218764714&location=UTR3&five='1') |
| [rs1369475742](https://guolab.wchscu.cn/miRNASNP/#!/snp?snp_id=rs1369475742&location=UTR3&one=1) | chr2:190970670 | C/T | 1/- | [1](https://guolab.wchscu.cn/miRNASNP/#!/snp?snp_id=rs1369475742&location=UTR3&four='1') | [10](https://guolab.wchscu.cn/miRNASNP/#!/snp?snp_id=rs1369475742&location=UTR3&five='1') |
| [rs774654212](https://guolab.wchscu.cn/miRNASNP/#!/snp?snp_id=rs774654212&location=UTR3&one=1) | chr2:190970674 | C/T | 1/- | [11](https://guolab.wchscu.cn/miRNASNP/#!/snp?snp_id=rs774654212&location=UTR3&four='1') | [1](https://guolab.wchscu.cn/miRNASNP/#!/snp?snp_id=rs774654212&location=UTR3&five='1') |
| [rs762042700](https://guolab.wchscu.cn/miRNASNP/#!/snp?snp_id=rs762042700&location=UTR3&one=1) | chr2:190970675 | G/A | 1/- | [6](https://guolab.wchscu.cn/miRNASNP/#!/snp?snp_id=rs762042700&location=UTR3&four='1') | [1](https://guolab.wchscu.cn/miRNASNP/#!/snp?snp_id=rs762042700&location=UTR3&five='1') |
| [rs1235760340](https://guolab.wchscu.cn/miRNASNP/#!/snp?snp_id=rs1235760340&location=UTR3&one=1) | chr2:190970678 | AGAG/A | 1.0/- | [4](https://guolab.wchscu.cn/miRNASNP/#!/snp?snp_id=rs1235760340&location=UTR3&four='1') | [6](https://guolab.wchscu.cn/miRNASNP/#!/snp?snp_id=rs1235760340&location=UTR3&five='1') |
| [rs1255205369](https://guolab.wchscu.cn/miRNASNP/#!/snp?snp_id=rs1255205369&location=UTR3&one=1) | chr2:190970679 | G/C | -/- | 0 | [4](https://guolab.wchscu.cn/miRNASNP/#!/snp?snp_id=rs1255205369&location=UTR3&five='1') |
| [rs1177503647](https://guolab.wchscu.cn/miRNASNP/#!/snp?snp_id=rs1177503647&location=UTR3&one=1) | chr2:190970684 | G/A | 1/- | [1](https://guolab.wchscu.cn/miRNASNP/#!/snp?snp_id=rs1177503647&location=UTR3&four='1') | [3](https://guolab.wchscu.cn/miRNASNP/#!/snp?snp_id=rs1177503647&location=UTR3&five='1') |
| [rs947648115](https://guolab.wchscu.cn/miRNASNP/#!/snp?snp_id=rs947648115&location=UTR3&one=1) | chr2:190970686 | T/C | -/- | 0 | [2](https://guolab.wchscu.cn/miRNASNP/#!/snp?snp_id=rs947648115&location=UTR3&five='1') |
| [rs767892470](https://guolab.wchscu.cn/miRNASNP/#!/snp?snp_id=rs767892470&location=UTR3&one=1) | chr2:190970694 | A/G | 1/- | 0 | 0 |
| [rs773624561](https://guolab.wchscu.cn/miRNASNP/#!/snp?snp_id=rs773624561&location=UTR3&one=1) | chr2:190970696 | T/C | -/- | 0 | 0 |
| [rs983566520](https://guolab.wchscu.cn/miRNASNP/#!/snp?snp_id=rs983566520&location=UTR3&one=1) | chr2:190975544 | T/G | -/- | [5](https://guolab.wchscu.cn/miRNASNP/#!/snp?snp_id=rs983566520&location=UTR3&four='1') | [1](https://guolab.wchscu.cn/miRNASNP/#!/snp?snp_id=rs983566520&location=UTR3&five='1') |
| [rs1412164181](https://guolab.wchscu.cn/miRNASNP/#!/snp?snp_id=rs1412164181&location=UTR3&one=1) | chr2:190975561 | T/C | 1/- | [1](https://guolab.wchscu.cn/miRNASNP/#!/snp?snp_id=rs1412164181&location=UTR3&four='1') | 0 |
| [rs1335503966](https://guolab.wchscu.cn/miRNASNP/#!/snp?snp_id=rs1335503966&location=UTR3&one=1) | chr2:190975573 | G/C | -/- | [2](https://guolab.wchscu.cn/miRNASNP/#!/snp?snp_id=rs1335503966&location=UTR3&four='1') | [4](https://guolab.wchscu.cn/miRNASNP/#!/snp?snp_id=rs1335503966&location=UTR3&five='1') |
| [rs1412318644](https://guolab.wchscu.cn/miRNASNP/#!/snp?snp_id=rs1412318644&location=UTR3&one=1) | chr2:190975574 | G/A | 1/- | [4](https://guolab.wchscu.cn/miRNASNP/#!/snp?snp_id=rs1412318644&location=UTR3&four='1') | [4](https://guolab.wchscu.cn/miRNASNP/#!/snp?snp_id=rs1412318644&location=UTR3&five='1') |
| [rs1389517862](https://guolab.wchscu.cn/miRNASNP/#!/snp?snp_id=rs1389517862&location=UTR3&one=1) | chr2:190975578 | T/A | -/- | [4](https://guolab.wchscu.cn/miRNASNP/#!/snp?snp_id=rs1389517862&location=UTR3&four='1') | [4](https://guolab.wchscu.cn/miRNASNP/#!/snp?snp_id=rs1389517862&location=UTR3&five='1') |
| [rs77910835](https://guolab.wchscu.cn/miRNASNP/#!/snp?snp_id=rs77910835&location=UTR3&one=1) | chr2:190975643 | A/T | 0.9921/0.0079 | 0 | [28](https://guolab.wchscu.cn/miRNASNP/#!/snp?snp_id=rs77910835&location=UTR3&five='1') |
| [rs1280181707](https://guolab.wchscu.cn/miRNASNP/#!/snp?snp_id=rs1280181707&location=UTR3&one=1) | chr2:190975645 | G/T | -/- | 0 | [29](https://guolab.wchscu.cn/miRNASNP/#!/snp?snp_id=rs1280181707&location=UTR3&five='1') |
| [rs112071828](https://guolab.wchscu.cn/miRNASNP/#!/snp?snp_id=rs112071828&location=UTR3&one=1) | chr2:190975648 | G/A | 0.9999/0.0001 | [1](https://guolab.wchscu.cn/miRNASNP/#!/snp?snp_id=rs112071828&location=UTR3&four='1') | [2](https://guolab.wchscu.cn/miRNASNP/#!/snp?snp_id=rs112071828&location=UTR3&five='1') |
| [rs1047419091](https://guolab.wchscu.cn/miRNASNP/#!/snp?snp_id=rs1047419091&location=UTR3&one=1) | chr2:190975655 | G/A | 1/- | [4](https://guolab.wchscu.cn/miRNASNP/#!/snp?snp_id=rs1047419091&location=UTR3&four='1') | [5](https://guolab.wchscu.cn/miRNASNP/#!/snp?snp_id=rs1047419091&location=UTR3&five='1') |
| [rs566136866](https://guolab.wchscu.cn/miRNASNP/#!/snp?snp_id=rs566136866&location=UTR3&one=1) | chr2:190975672 | C/A | 1/- | [3](https://guolab.wchscu.cn/miRNASNP/#!/snp?snp_id=rs566136866&location=UTR3&four='1') | [4](https://guolab.wchscu.cn/miRNASNP/#!/snp?snp_id=rs566136866&location=UTR3&five='1') |
| [rs566136866](https://guolab.wchscu.cn/miRNASNP/#!/snp?snp_id=rs566136866&location=UTR3&one=1) | chr2:190975672 | C/T | 1/- | [10](https://guolab.wchscu.cn/miRNASNP/#!/snp?snp_id=rs566136866&location=UTR3&four='1') | [4](https://guolab.wchscu.cn/miRNASNP/#!/snp?snp_id=rs566136866&location=UTR3&five='1') |
| [rs940656467](https://guolab.wchscu.cn/miRNASNP/#!/snp?snp_id=rs940656467&location=UTR3&one=1) | chr2:190975673 | G/A | 1/- | [6](https://guolab.wchscu.cn/miRNASNP/#!/snp?snp_id=rs940656467&location=UTR3&four='1') | [5](https://guolab.wchscu.cn/miRNASNP/#!/snp?snp_id=rs940656467&location=UTR3&five='1') |
| [rs116554639](https://guolab.wchscu.cn/miRNASNP/#!/snp?snp_id=rs116554639&location=UTR3&one=1) | chr2:190975678 | T/A | 0.9991/0.0009 | [9](https://guolab.wchscu.cn/miRNASNP/#!/snp?snp_id=rs116554639&location=UTR3&four='1') | [7](https://guolab.wchscu.cn/miRNASNP/#!/snp?snp_id=rs116554639&location=UTR3&five='1') |
| [rs1471423016](https://guolab.wchscu.cn/miRNASNP/#!/snp?snp_id=rs1471423016&location=UTR3&one=1) | chr2:190975683 | A/T | 1/- | [5](https://guolab.wchscu.cn/miRNASNP/#!/snp?snp_id=rs1471423016&location=UTR3&four='1') | [6](https://guolab.wchscu.cn/miRNASNP/#!/snp?snp_id=rs1471423016&location=UTR3&five='1') |
| [rs554936429](https://guolab.wchscu.cn/miRNASNP/#!/snp?snp_id=rs554936429&location=UTR3&one=1) | chr2:190975684 | C/G | -/- | [2](https://guolab.wchscu.cn/miRNASNP/#!/snp?snp_id=rs554936429&location=UTR3&four='1') | [4](https://guolab.wchscu.cn/miRNASNP/#!/snp?snp_id=rs554936429&location=UTR3&five='1') |
| [rs1410817426](https://guolab.wchscu.cn/miRNASNP/#!/snp?snp_id=rs1410817426&location=UTR3&one=1) | chr2:190975698 | A/T | 1/- | 0 | [5](https://guolab.wchscu.cn/miRNASNP/#!/snp?snp_id=rs1410817426&location=UTR3&five='1') |
| [rs899224935](https://guolab.wchscu.cn/miRNASNP/#!/snp?snp_id=rs899224935&location=UTR3&one=1) | chr2:190975702 | C/T | -/- | [5](https://guolab.wchscu.cn/miRNASNP/#!/snp?snp_id=rs899224935&location=UTR3&four='1') | [3](https://guolab.wchscu.cn/miRNASNP/#!/snp?snp_id=rs899224935&location=UTR3&five='1') |
| [rs575057281](https://guolab.wchscu.cn/miRNASNP/#!/snp?snp_id=rs575057281&location=UTR3&one=1) | chr2:190975703 | G/A | 1/- | [2](https://guolab.wchscu.cn/miRNASNP/#!/snp?snp_id=rs575057281&location=UTR3&four='1') | [3](https://guolab.wchscu.cn/miRNASNP/#!/snp?snp_id=rs575057281&location=UTR3&five='1') |
| [rs575057281](https://guolab.wchscu.cn/miRNASNP/#!/snp?snp_id=rs575057281&location=UTR3&one=1) | chr2:190975703 | G/C | 1/- | [7](https://guolab.wchscu.cn/miRNASNP/#!/snp?snp_id=rs575057281&location=UTR3&four='1') | [3](https://guolab.wchscu.cn/miRNASNP/#!/snp?snp_id=rs575057281&location=UTR3&five='1') |
| [rs961200796](https://guolab.wchscu.cn/miRNASNP/#!/snp?snp_id=rs961200796&location=UTR3&one=1) | chr2:190975705 | T/C | -/- | [6](https://guolab.wchscu.cn/miRNASNP/#!/snp?snp_id=rs961200796&location=UTR3&four='1') | [3](https://guolab.wchscu.cn/miRNASNP/#!/snp?snp_id=rs961200796&location=UTR3&five='1') |
| [rs45623541](https://guolab.wchscu.cn/miRNASNP/#!/snp?snp_id=rs45623541&location=UTR3&one=1) | chr2:190975708 | C/T | 0.9987/0.0013 | 0 | [5](https://guolab.wchscu.cn/miRNASNP/#!/snp?snp_id=rs45623541&location=UTR3&five='1') |
| [rs1029197409](https://guolab.wchscu.cn/miRNASNP/#!/snp?snp_id=rs1029197409&location=UTR3&one=1) | chr2:190975715 | A/G | 1/- | [3](https://guolab.wchscu.cn/miRNASNP/#!/snp?snp_id=rs1029197409&location=UTR3&four='1') | [3](https://guolab.wchscu.cn/miRNASNP/#!/snp?snp_id=rs1029197409&location=UTR3&five='1') |
| [rs1274851520](https://guolab.wchscu.cn/miRNASNP/#!/snp?snp_id=rs1274851520&location=UTR3&one=1) | chr2:190975723 | A/C | 1/- | [2](https://guolab.wchscu.cn/miRNASNP/#!/snp?snp_id=rs1274851520&location=UTR3&four='1') | [32](https://guolab.wchscu.cn/miRNASNP/#!/snp?snp_id=rs1274851520&location=UTR3&five='1') |
| [rs927902093](https://guolab.wchscu.cn/miRNASNP/#!/snp?snp_id=rs927902093&location=UTR3&one=1) | chr2:190975729 | A/G | -/- | 0 | [3](https://guolab.wchscu.cn/miRNASNP/#!/snp?snp_id=rs927902093&location=UTR3&five='1') |
| [rs1179362530](https://guolab.wchscu.cn/miRNASNP/#!/snp?snp_id=rs1179362530&location=UTR3&one=1) | chr2:190975734 | A/G | -/- | [7](https://guolab.wchscu.cn/miRNASNP/#!/snp?snp_id=rs1179362530&location=UTR3&four='1') | [3](https://guolab.wchscu.cn/miRNASNP/#!/snp?snp_id=rs1179362530&location=UTR3&five='1') |
| [rs890617468](https://guolab.wchscu.cn/miRNASNP/#!/snp?snp_id=rs890617468&location=UTR3&one=1) | chr2:190975745 | C/T | 1/- | [4](https://guolab.wchscu.cn/miRNASNP/#!/snp?snp_id=rs890617468&location=UTR3&four='1') | [6](https://guolab.wchscu.cn/miRNASNP/#!/snp?snp_id=rs890617468&location=UTR3&five='1') |
| [rs1328729536](https://guolab.wchscu.cn/miRNASNP/#!/snp?snp_id=rs1328729536&location=UTR3&one=1) | chr2:190975752 | C/T | 1/- | [5](https://guolab.wchscu.cn/miRNASNP/#!/snp?snp_id=rs1328729536&location=UTR3&four='1') | [6](https://guolab.wchscu.cn/miRNASNP/#!/snp?snp_id=rs1328729536&location=UTR3&five='1') |
| [rs201391478](https://guolab.wchscu.cn/miRNASNP/#!/snp?snp_id=rs201391478&location=UTR3&one=1) | chr2:190975760 | A/G | -/- | [11](https://guolab.wchscu.cn/miRNASNP/#!/snp?snp_id=rs201391478&location=UTR3&four='1') | [8](https://guolab.wchscu.cn/miRNASNP/#!/snp?snp_id=rs201391478&location=UTR3&five='1') |
| [rs752978951](https://guolab.wchscu.cn/miRNASNP/#!/snp?snp_id=rs752978951&location=UTR3&one=1) | chr2:190975761 | GC/G | -/- | [9](https://guolab.wchscu.cn/miRNASNP/#!/snp?snp_id=rs752978951&location=UTR3&four='1') | [7](https://guolab.wchscu.cn/miRNASNP/#!/snp?snp_id=rs752978951&location=UTR3&five='1') |
| [rs753796178](https://guolab.wchscu.cn/miRNASNP/#!/snp?snp_id=rs753796178&location=UTR3&one=1) | chr2:190975764 | A/G | -/- | [20](https://guolab.wchscu.cn/miRNASNP/#!/snp?snp_id=rs753796178&location=UTR3&four='1') | [5](https://guolab.wchscu.cn/miRNASNP/#!/snp?snp_id=rs753796178&location=UTR3&five='1') |
| [rs11549698](https://guolab.wchscu.cn/miRNASNP/#!/snp?snp_id=rs11549698&location=UTR3&one=1) | chr2:190975766 | G/C | 0.9986/0.0014 | [7](https://guolab.wchscu.cn/miRNASNP/#!/snp?snp_id=rs11549698&location=UTR3&four='1') | [10](https://guolab.wchscu.cn/miRNASNP/#!/snp?snp_id=rs11549698&location=UTR3&five='1') |
| [rs779046006](https://guolab.wchscu.cn/miRNASNP/#!/snp?snp_id=rs779046006&location=UTR3&one=1) | chr2:190975770 | G/T | -/- | [8](https://guolab.wchscu.cn/miRNASNP/#!/snp?snp_id=rs779046006&location=UTR3&four='1') | [12](https://guolab.wchscu.cn/miRNASNP/#!/snp?snp_id=rs779046006&location=UTR3&five='1') |
| [rs369953295](https://guolab.wchscu.cn/miRNASNP/#!/snp?snp_id=rs369953295&location=UTR3&one=1) | chr2:190975772 | T/A | 1/- | [9](https://guolab.wchscu.cn/miRNASNP/#!/snp?snp_id=rs369953295&location=UTR3&four='1') | [7](https://guolab.wchscu.cn/miRNASNP/#!/snp?snp_id=rs369953295&location=UTR3&five='1') |
| [rs1284802601](https://guolab.wchscu.cn/miRNASNP/#!/snp?snp_id=rs1284802601&location=UTR3&one=1) | chr2:190975773 | T/G | -/- | [8](https://guolab.wchscu.cn/miRNASNP/#!/snp?snp_id=rs1284802601&location=UTR3&four='1') | [1](https://guolab.wchscu.cn/miRNASNP/#!/snp?snp_id=rs1284802601&location=UTR3&five='1') |
| [rs748181005](https://guolab.wchscu.cn/miRNASNP/#!/snp?snp_id=rs748181005&location=UTR3&one=1) | chr2:190975776 | G/A | -/- | [3](https://guolab.wchscu.cn/miRNASNP/#!/snp?snp_id=rs748181005&location=UTR3&four='1') | [8](https://guolab.wchscu.cn/miRNASNP/#!/snp?snp_id=rs748181005&location=UTR3&five='1') |
| [rs1365587496](https://guolab.wchscu.cn/miRNASNP/#!/snp?snp_id=rs1365587496&location=UTR3&one=1) | chr2:190975777 | G/GT | -/- | [4](https://guolab.wchscu.cn/miRNASNP/#!/snp?snp_id=rs1365587496&location=UTR3&four='1') | [2](https://guolab.wchscu.cn/miRNASNP/#!/snp?snp_id=rs1365587496&location=UTR3&five='1') |
| [rs1211217813](https://guolab.wchscu.cn/miRNASNP/#!/snp?snp_id=rs1211217813&location=UTR3&one=1) | chr2:190975786 | C/T | -/- | [1](https://guolab.wchscu.cn/miRNASNP/#!/snp?snp_id=rs1211217813&location=UTR3&four='1') | [5](https://guolab.wchscu.cn/miRNASNP/#!/snp?snp_id=rs1211217813&location=UTR3&five='1') |
| [rs372657991](https://guolab.wchscu.cn/miRNASNP/#!/snp?snp_id=rs372657991&location=UTR3&one=1) | chr2:190975792 | A/C | 1/- | [1](https://guolab.wchscu.cn/miRNASNP/#!/snp?snp_id=rs372657991&location=UTR3&four='1') | 0 |
| [rs777991780](https://guolab.wchscu.cn/miRNASNP/#!/snp?snp_id=rs777991780&location=UTR3&one=1) | chr2:190975796 | T/C | -/- | 0 | 0 |
| [rs1442310621](https://guolab.wchscu.cn/miRNASNP/#!/snp?snp_id=rs1442310621&location=UTR3&one=1) | chr2:190975799 | T/C | 1/- | 0 | 0 |
| [rs1204145559](https://guolab.wchscu.cn/miRNASNP/#!/snp?snp_id=rs1204145559&location=UTR3&one=1) | chr2:191029830 | G/GTT | -/- | 0 | 0 |
| [rs1413622801](https://guolab.wchscu.cn/miRNASNP/#!/snp?snp_id=rs1413622801&location=UTR3&one=1) | chr2:190969845 | T/C | 1/- | [4](https://guolab.wchscu.cn/miRNASNP/#!/snp?snp_id=rs1413622801&location=UTR3&four='1') | 0 |
| [rs1425238981](https://guolab.wchscu.cn/miRNASNP/#!/snp?snp_id=rs1425238981&location=UTR3&one=1) | chr2:190969847 | T/C | 1/- | 0 | 0 |
| [rs188557905](https://guolab.wchscu.cn/miRNASNP/#!/snp?snp_id=rs188557905&location=UTR3&one=1) | chr2:190969848 | C/T | 0.9995/0.0005 | [1](https://guolab.wchscu.cn/miRNASNP/#!/snp?snp_id=rs188557905&location=UTR3&four='1') | 0 |
| [rs1214688680](https://guolab.wchscu.cn/miRNASNP/#!/snp?snp_id=rs1214688680&location=UTR3&one=1) | chr2:190969852 | A/G | -/- | [1](https://guolab.wchscu.cn/miRNASNP/#!/snp?snp_id=rs1214688680&location=UTR3&four='1') | 0 |
| [rs1474363657](https://guolab.wchscu.cn/miRNASNP/#!/snp?snp_id=rs1474363657&location=UTR3&one=1) | chr2:190969856 | A/G | 1/- | [3](https://guolab.wchscu.cn/miRNASNP/#!/snp?snp_id=rs1474363657&location=UTR3&four='1') | [3](https://guolab.wchscu.cn/miRNASNP/#!/snp?snp_id=rs1474363657&location=UTR3&five='1') |
| [rs1288525683](https://guolab.wchscu.cn/miRNASNP/#!/snp?snp_id=rs1288525683&location=UTR3&one=1) | chr2:190969873 | T/C | -/- | [5](https://guolab.wchscu.cn/miRNASNP/#!/snp?snp_id=rs1288525683&location=UTR3&four='1') | [1](https://guolab.wchscu.cn/miRNASNP/#!/snp?snp_id=rs1288525683&location=UTR3&five='1') |
| [rs1422860396](https://guolab.wchscu.cn/miRNASNP/#!/snp?snp_id=rs1422860396&location=UTR3&one=1) | chr2:190969875 | G/C | 1/- | [2](https://guolab.wchscu.cn/miRNASNP/#!/snp?snp_id=rs1422860396&location=UTR3&four='1') | 0 |
| [rs1194364922](https://guolab.wchscu.cn/miRNASNP/#!/snp?snp_id=rs1194364922&location=UTR3&one=1) | chr2:190969878 | T/C | 1/- | [1](https://guolab.wchscu.cn/miRNASNP/#!/snp?snp_id=rs1194364922&location=UTR3&four='1') | [4](https://guolab.wchscu.cn/miRNASNP/#!/snp?snp_id=rs1194364922&location=UTR3&five='1') |
| [rs903239098](https://guolab.wchscu.cn/miRNASNP/#!/snp?snp_id=rs903239098&location=UTR3&one=1) | chr2:190969882 | T/C | -/- | [2](https://guolab.wchscu.cn/miRNASNP/#!/snp?snp_id=rs903239098&location=UTR3&four='1') | [10](https://guolab.wchscu.cn/miRNASNP/#!/snp?snp_id=rs903239098&location=UTR3&five='1') |
| [rs1036975160](https://guolab.wchscu.cn/miRNASNP/#!/snp?snp_id=rs1036975160&location=UTR3&one=1) | chr2:190969884 | G/A | 1/- | [10](https://guolab.wchscu.cn/miRNASNP/#!/snp?snp_id=rs1036975160&location=UTR3&four='1') | [9](https://guolab.wchscu.cn/miRNASNP/#!/snp?snp_id=rs1036975160&location=UTR3&five='1') |
| [rs1319612138](https://guolab.wchscu.cn/miRNASNP/#!/snp?snp_id=rs1319612138&location=UTR3&one=1) | chr2:190969889 | GA/G | -/- | [5](https://guolab.wchscu.cn/miRNASNP/#!/snp?snp_id=rs1319612138&location=UTR3&four='1') | [3](https://guolab.wchscu.cn/miRNASNP/#!/snp?snp_id=rs1319612138&location=UTR3&five='1') |
| [rs907349445](https://guolab.wchscu.cn/miRNASNP/#!/snp?snp_id=rs907349445&location=UTR3&one=1) | chr2:190969889 | G/T | 1/- | [3](https://guolab.wchscu.cn/miRNASNP/#!/snp?snp_id=rs907349445&location=UTR3&four='1') | [4](https://guolab.wchscu.cn/miRNASNP/#!/snp?snp_id=rs907349445&location=UTR3&five='1') |
| [rs1033141232](https://guolab.wchscu.cn/miRNASNP/#!/snp?snp_id=rs1033141232&location=UTR3&one=1) | chr2:190969894 | G/A | 1/- | 0 | [1](https://guolab.wchscu.cn/miRNASNP/#!/snp?snp_id=rs1033141232&location=UTR3&five='1') |
| [rs958955361](https://guolab.wchscu.cn/miRNASNP/#!/snp?snp_id=rs958955361&location=UTR3&one=1) | chr2:190969896 | C/G | 1/- | 0 | [1](https://guolab.wchscu.cn/miRNASNP/#!/snp?snp_id=rs958955361&location=UTR3&five='1') |
| [rs1013154846](https://guolab.wchscu.cn/miRNASNP/#!/snp?snp_id=rs1013154846&location=UTR3&one=1) | chr2:190969898 | A/G | 1/- | [3](https://guolab.wchscu.cn/miRNASNP/#!/snp?snp_id=rs1013154846&location=UTR3&four='1') | [1](https://guolab.wchscu.cn/miRNASNP/#!/snp?snp_id=rs1013154846&location=UTR3&five='1') |

Table 6. Functionally significant 3 prime UTR SNP analysis by MicroSNIper.

| SNP ID | Range | MiRNAs |
| --- | --- | --- |
| rs11305 | chr2:191833762-191835428 | hsa-miR-656  hsa-miR-4795-3p  hsa-miR-4528  hsa-miR-6083  hsa-miR-556-3p  hsa-miR-3669  hsa-miR-3145-3p  hsa-miR-556-3p |
| rs189030575 | chr2:191833762-191835428 | hsa-miR-5692b  hsa-miR-5692c  hsa-miR-4703-3p  hsa-miR-3121-3p  hsa-miR-219-1-3p |
| rs41324145 | chr2:191833762-191835428 | hsa-miR-4448  hsa-miR-483-3p  hsa-miR-934  hsa-miR-335-3p  hsa-miR-4482-3p  hsa-miR-501-5p  hsa-miR-3124-3p  hsa-miR-508-5p  hsa-miR-617  hsa-miR-3680-5p  hsa-miR-510  hsa-miR-1304-3p  hsa-miR-4685-3p  hsa-miR-4287  hsa-miR-4755-5p  hsa-miR-5006-3p  hsa-miR-4469  hsa-miR-934  hsa-miR-211-5p  hsa-miR-3667-3p  hsa-miR-5581-5p  hsa-miR-623  hsa-miR-204-5p  hsa-miR-5006-3p  hsa-miR-3127-3p  hsa-miR-4446-5p  hsa-miR-579  hsa-miR-4297 |
| rs184180073 | chr2:191833762-191835428 | hsa-miR-150-5p  hsa-miR-4755-5p  hsa-miR-3679-3p  hsa-miR-5006-3p |
| rs41363648 | chr2:191833762-191835428 | hsa-miR-4474-3p  hsa-miR-5010-3p  hsa-miR-32-3p  hsa-miR-4752  hsa-miR-758-3p  hsa-miR-4778-5p  hsa-miR-5571-5p  hsa-miR-4733-5p  hsa-miR-3182  hsa-miR-4265  hsa-miR-4322  hsa-miR-4652-3p  hsa-miR-150-5p |
| rs188557905 | chr2:191833762-191835428 | hsa-miR-222-3p  hsa-miR-3606-3p  hsa-miR-3618  hsa-miR-4743-3p  hsa-miR-139-5p  hsa-miR-4699-3p |
| rs180904823 | chr2:191833762-191835428 | hsa-miR-548g-3p  [hsa-miR-302f](http://microrna.sanger.ac.uk/cgi-bin/mirna_entry.pl?acc=hsa-miR-302f)  [hsa-miR-369-3p](http://microrna.sanger.ac.uk/cgi-bin/mirna_entry.pl?acc=hsa-miR-369-3p)  [hsa-miR-548g-3p](http://microrna.sanger.ac.uk/cgi-bin/mirna_entry.pl?acc=hsa-miR-548g-3p)  [hsa-miR-675-3p](http://microrna.sanger.ac.uk/cgi-bin/mirna_entry.pl?acc=hsa-miR-675-3p)  [hsa-miR-203b-3p](http://microrna.sanger.ac.uk/cgi-bin/mirna_entry.pl?acc=hsa-miR-203b-3p)  [hsa-miR-20b-3p](http://microrna.sanger.ac.uk/cgi-bin/mirna_entry.pl?acc=hsa-miR-20b-3p) |
| rs186033487 |  |  |
| rs146036682 | chr2:191833762-191835428 | [hsa-miR-4766-5p](http://microrna.sanger.ac.uk/cgi-bin/mirna_entry.pl?acc=hsa-miR-4766-5p)  [hsa-miR-4662b](http://microrna.sanger.ac.uk/cgi-bin/mirna_entry.pl?acc=hsa-miR-4662b)  [hsa-miR-3171](http://microrna.sanger.ac.uk/cgi-bin/mirna_entry.pl?acc=hsa-miR-3171)  [hsa-miR-4426](http://microrna.sanger.ac.uk/cgi-bin/mirna_entry.pl?acc=hsa-miR-4426)  [hsa-miR-4311](http://microrna.sanger.ac.uk/cgi-bin/mirna_entry.pl?acc=hsa-miR-4311)  [hsa-miR-4657](http://microrna.sanger.ac.uk/cgi-bin/mirna_entry.pl?acc=hsa-miR-4657)  [hsa-miR-4647](http://microrna.sanger.ac.uk/cgi-bin/mirna_entry.pl?acc=hsa-miR-4647)  [hsa-miR-4698](http://microrna.sanger.ac.uk/cgi-bin/mirna_entry.pl?acc=hsa-miR-4698)  [hsa-miR-4495](http://microrna.sanger.ac.uk/cgi-bin/mirna_entry.pl?acc=hsa-miR-4495)  [hsa-miR-522-3p](http://microrna.sanger.ac.uk/cgi-bin/mirna_entry.pl?acc=hsa-miR-522-3p)  [hsa-miR-3607-5p](http://microrna.sanger.ac.uk/cgi-bin/mirna_entry.pl?acc=hsa-miR-3607-5p)  [hsa-miR-103b](http://microrna.sanger.ac.uk/cgi-bin/mirna_entry.pl?acc=hsa-miR-103b)  [hsa-miR-224-3p](http://microrna.sanger.ac.uk/cgi-bin/mirna_entry.pl?acc=hsa-miR-224-3p) |
| rs190508584 | chr2:191833762-191835428 | hsa-miR-380-3p  [hsa-miR-642b-3p](http://microrna.sanger.ac.uk/cgi-bin/mirna_entry.pl?acc=hsa-miR-642b-3p)  [hsa-miR-642a-3p](http://microrna.sanger.ac.uk/cgi-bin/mirna_entry.pl?acc=hsa-miR-642a-3p)  [hsa-miR-466](http://microrna.sanger.ac.uk/cgi-bin/mirna_entry.pl?acc=hsa-miR-466)  [hsa-miR-23b-3p](http://microrna.sanger.ac.uk/cgi-bin/mirna_entry.pl?acc=hsa-miR-23b-3p)  [hsa-miR-4717-3p](http://microrna.sanger.ac.uk/cgi-bin/mirna_entry.pl?acc=hsa-miR-4717-3p)  [hsa-miR-323a-3p](http://microrna.sanger.ac.uk/cgi-bin/mirna_entry.pl?acc=hsa-miR-323a-3p)  [hsa-miR-4672](http://microrna.sanger.ac.uk/cgi-bin/mirna_entry.pl?acc=hsa-miR-4672)  [hsa-miR-23a-3p](http://microrna.sanger.ac.uk/cgi-bin/mirna_entry.pl?acc=hsa-miR-23a-3p)  [hsa-miR-130a-5p](http://microrna.sanger.ac.uk/cgi-bin/mirna_entry.pl?acc=hsa-miR-130a-5p)  [hsa-miR-4643](http://microrna.sanger.ac.uk/cgi-bin/mirna_entry.pl?acc=hsa-miR-4643)  [hsa-miR-380-3p](http://microrna.sanger.ac.uk/cgi-bin/mirna_entry.pl?acc=hsa-miR-380-3p)  [hsa-miR-5590-3p](http://microrna.sanger.ac.uk/cgi-bin/mirna_entry.pl?acc=hsa-miR-5590-3p) |
| rs114360225 | chr2:191833762-191835428 | [hsa-miR-380-3p](http://microrna.sanger.ac.uk/cgi-bin/mirna_entry.pl?acc=hsa-miR-380-3p)  [hsa-miR-642b-3p](http://microrna.sanger.ac.uk/cgi-bin/mirna_entry.pl?acc=hsa-miR-642b-3p)  [hsa-miR-642a-3p](http://microrna.sanger.ac.uk/cgi-bin/mirna_entry.pl?acc=hsa-miR-642a-3p)  [hsa-miR-466](http://microrna.sanger.ac.uk/cgi-bin/mirna_entry.pl?acc=hsa-miR-466)  [hsa-miR-23b-3p](http://microrna.sanger.ac.uk/cgi-bin/mirna_entry.pl?acc=hsa-miR-23b-3p)  [hsa-miR-4717-3p](http://microrna.sanger.ac.uk/cgi-bin/mirna_entry.pl?acc=hsa-miR-4717-3p)  [hsa-miR-323a-3p](http://microrna.sanger.ac.uk/cgi-bin/mirna_entry.pl?acc=hsa-miR-323a-3p)  [hsa-miR-23a-3p](http://microrna.sanger.ac.uk/cgi-bin/mirna_entry.pl?acc=hsa-miR-23a-3p)  [hsa-miR-130a-5p](http://microrna.sanger.ac.uk/cgi-bin/mirna_entry.pl?acc=hsa-miR-130a-5p)  [hsa-miR-4643](http://microrna.sanger.ac.uk/cgi-bin/mirna_entry.pl?acc=hsa-miR-4643)  [hsa-miR-380-3p](http://microrna.sanger.ac.uk/cgi-bin/mirna_entry.pl?acc=hsa-miR-380-3p)  [hsa-miR-603](http://microrna.sanger.ac.uk/cgi-bin/mirna_entry.pl?acc=hsa-miR-603)  [hsa-miR-3124-3p](http://microrna.sanger.ac.uk/cgi-bin/mirna_entry.pl?acc=hsa-miR-3124-3p)  [hsa-miR-466](http://microrna.sanger.ac.uk/cgi-bin/mirna_entry.pl?acc=hsa-miR-466)  [hsa-miR-6500-3p](http://microrna.sanger.ac.uk/cgi-bin/mirna_entry.pl?acc=hsa-miR-6500-3p) |
| rs139958571 | chr2:191833762-191835428 | [hsa-miR-186-5p](http://microrna.sanger.ac.uk/cgi-bin/mirna_entry.pl?acc=hsa-miR-186-5p)  [hsa-miR-4713-5p](http://microrna.sanger.ac.uk/cgi-bin/mirna_entry.pl?acc=hsa-miR-4713-5p)  [hsa-miR-188-3p](http://microrna.sanger.ac.uk/cgi-bin/mirna_entry.pl?acc=hsa-miR-188-3p)  [hsa-miR-629-3p](http://microrna.sanger.ac.uk/cgi-bin/mirna_entry.pl?acc=hsa-miR-629-3p)  [hsa-miR-532-3p](http://microrna.sanger.ac.uk/cgi-bin/mirna_entry.pl?acc=hsa-miR-532-3p)  [hsa-miR-3156-3p](http://microrna.sanger.ac.uk/cgi-bin/mirna_entry.pl?acc=hsa-miR-3156-3p)  [hsa-miR-150-5p](http://microrna.sanger.ac.uk/cgi-bin/mirna_entry.pl?acc=hsa-miR-150-5p)  [hsa-miR-4433-5p](http://microrna.sanger.ac.uk/cgi-bin/mirna_entry.pl?acc=hsa-miR-4433-5p)  [hsa-miR-4452](http://microrna.sanger.ac.uk/cgi-bin/mirna_entry.pl?acc=hsa-miR-4452) |
| rs182394503 | chr2:191833762-191835428 | [hsa-miR-598](http://microrna.sanger.ac.uk/cgi-bin/mirna_entry.pl?acc=hsa-miR-598)  [hsa-miR-505-3p](http://microrna.sanger.ac.uk/cgi-bin/mirna_entry.pl?acc=hsa-miR-505-3p)  [hsa-miR-337-3p](http://microrna.sanger.ac.uk/cgi-bin/mirna_entry.pl?acc=hsa-miR-337-3p)  [hsa-miR-654-3p](http://microrna.sanger.ac.uk/cgi-bin/mirna_entry.pl?acc=hsa-miR-654-3p) |
| rs186032149 | chr2:191833762-191835428 | [hsa-miR-4714-3p](http://microrna.sanger.ac.uk/cgi-bin/mirna_entry.pl?acc=hsa-miR-4714-3p)  [hsa-miR-337-3p](http://microrna.sanger.ac.uk/cgi-bin/mirna_entry.pl?acc=hsa-miR-337-3p)  [hsa-miR-3150b-5p](http://microrna.sanger.ac.uk/cgi-bin/mirna_entry.pl?acc=hsa-miR-3150b-5p) |
| rs190542524 | chr2:191833762-191835428 | [hsa-miR-4804-3p](http://microrna.sanger.ac.uk/cgi-bin/mirna_entry.pl?acc=hsa-miR-4804-3p)  [hsa-miR-4279](http://microrna.sanger.ac.uk/cgi-bin/mirna_entry.pl?acc=hsa-miR-4279)  [hsa-miR-4448](http://microrna.sanger.ac.uk/cgi-bin/mirna_entry.pl?acc=hsa-miR-4448)  [hsa-miR-4804-3p](http://microrna.sanger.ac.uk/cgi-bin/mirna_entry.pl?acc=hsa-miR-4804-3p)  [hsa-miR-4720-3p](http://microrna.sanger.ac.uk/cgi-bin/mirna_entry.pl?acc=hsa-miR-4720-3p) |
| rs182725919 | chr2:191833762-191835428 | [hsa-miR-3126-3p](http://microrna.sanger.ac.uk/cgi-bin/mirna_entry.pl?acc=hsa-miR-3126-3p)  [hsa-miR-4466](http://microrna.sanger.ac.uk/cgi-bin/mirna_entry.pl?acc=hsa-miR-4466)  [hsa-miR-4701-3p](http://microrna.sanger.ac.uk/cgi-bin/mirna_entry.pl?acc=hsa-miR-4701-3p)  [hsa-miR-4711-5p](http://microrna.sanger.ac.uk/cgi-bin/mirna_entry.pl?acc=hsa-miR-4711-5p) |
| rs41481847 | chr2:191833762-191835428 | [hsa-miR-4303](http://microrna.sanger.ac.uk/cgi-bin/mirna_entry.pl?acc=hsa-miR-4303)  [hsa-miR-1303](http://microrna.sanger.ac.uk/cgi-bin/mirna_entry.pl?acc=hsa-miR-1303)  [hsa-miR-5680](http://microrna.sanger.ac.uk/cgi-bin/mirna_entry.pl?acc=hsa-miR-5680)  [hsa-miR-4668-5p](http://microrna.sanger.ac.uk/cgi-bin/mirna_entry.pl?acc=hsa-miR-4668-5p)  [hsa-miR-4303](http://microrna.sanger.ac.uk/cgi-bin/mirna_entry.pl?acc=hsa-miR-4303)  [hsa-miR-1258](http://microrna.sanger.ac.uk/cgi-bin/mirna_entry.pl?acc=hsa-miR-1258)  [hsa-miR-3162-5p](http://microrna.sanger.ac.uk/cgi-bin/mirna_entry.pl?acc=hsa-miR-3162-5p)  [hsa-miR-135a-3p](http://microrna.sanger.ac.uk/cgi-bin/mirna_entry.pl?acc=hsa-miR-135a-3p) |
